# Conformational transitions of the mitotic adaptor Spindly underlie its interaction with Dynein and Dynactin

**DOI:** 10.1101/2022.02.02.478874

**Authors:** Ennio d’Amico, Misbha Ud Din Ahmad, Verena Cmentowski, Mathias Girbig, Franziska Müller, Sabine Wohlgemuth, Andreas Brockmeyer, Stefano Maffini, Petra Janning, Ingrid R. Vetter, Andrew P. Carter, Anastassis Perrakis, Andrea Musacchio

**Author notes:** ZoBio BV, J.H. Oortweg19, 2333 CH Leiden, The Netherlands. European Molecular Biology Laboratory (EMBL), Structural and Computational Biology Unit, Meyerhofstrasse 1, 69117 Heidelberg, Germany. These authors contributed equally (Misbha Ud Din Ahmad, Verena Cmentowski).

## Abstract

Cytoplasmic Dynein 1, or Dynein, is a microtubule minus-end directed motor. Dynein motility requires Dynactin and a family of activating adaptors that stabilize the Dynein-Dynactin complex and promote regulated interactions with cargo in space and time. How activating adaptors limit Dynein activation to specialized subcellular locales is unclear. Here, we reveal that Spindly, a mitotic Dynein adaptor at the kinetochore corona, exists natively in a closed conformation that occludes binding of Dynein-Dynactin to its CC1 box and Spindly motif. A structure-based analysis identified various mutations promoting an open conformation of Spindly that binds Dynein-Dynactin. A region of Spindly downstream from the Spindly motif and not required for cargo binding faces the CC1 box and stabilizes the intramolecular closed conformation. This region is also required for robust kinetochore localization of Spindly, suggesting that kinetochores promote Spindly activation to recruit Dynein. This mechanism may be paradigmatic for Dynein activation by other adaptors at various cellular locales.

## Introduction

Eukaryotic cells maintain their internal order through the concerted action of a variety of functionally diverse energy-harnessing enzymes. Among these are molecular motors that convert the chemical energy of adenosine 5′-triphosphate (ATP) into mechanical work to dispatch various cargoes to different subcellular locations (Klinman and Holzbaur, 2018). Many molecular motors move along microtubules, polarized cellular tracks with plus- and minus-ends, the latter normally localized near microtubule-organizing centers, such as centrosomes. Motors use microtubules to transport cargoes of various size, ranging from individual protein complexes, to viruses, to organelles. Molecular motors also move chromosomes and transport, crosslink, and reciprocally slide microtubules to promote the assembly of the mitotic spindle in mitotic cells (Pavin and Tolic, 2021). The two main classes of intra-cellular molecular motors are the kinesins and Dynein. Kinesins populate a wide and diverse family of motors, prevalently with plus-end-directed polarity, with each family member specializing in the transport of distinct cargoes (Klinman and Holzbaur, 2018). Cytoplasmic Dynein-1 (Dynein), on the other hand, is a minus-end-directed multi-subunit assembly whose motor subunit, Dynein heavy chain (DHC), is encoded by a single gene. Its association with different cargoes relies therefore on various activating adaptors, each with a distinct cargo preference (Canty et al., 2021; Reck-Peterson et al., 2018; Roberts et al., 2013).

In humans, the 1.4 MDa Dynein complex consists of 12 subunits, with six different polypeptides all present in two copies, including the DHC, the intermediate chains (DIC), the light intermediate chains (LIC), and the three Dynein light chains LC8, Roadblock, and Tctex (Reck-Peterson et al., 2018). The motility of Dynein requires the 1.1 MDa complex Dynactin (Carter et al., 2016; Reck-Peterson et al., 2018). Dynactin consists of 23 polypeptides and 11 individual subunits, organized in four main structural domains: 1) a central actin-like filament consisting of eight ARP1 subunits and one actin; 2) a four-subunit pointed-end capping complex including the subunits p25, p27, p62, Arp11; 3) a barbed-end capping complex containing CapZαβ; and 4) a shoulder domain containing p24, p150^glued^, and p50/dynamitin (Chowdhury et al., 2015; Lau et al., 2021; Urnavicius et al., 2015).

The interaction of Dynein and Dynactin (DD) is weak but is strongly promoted by different classes of activating adaptors that also promote interactions with various cargoes (Hoogenraad and Akhmanova, 2016; Olenick and Holzbaur, 2019; Reck-Peterson et al., 2018). The prototypical activating adaptor, BICD2 (bicaudal D homologue 2), favors the incorporation of Dynein and Dynactin into a single complex with greatly increased motility and processivity in comparison to the isolated Dynein (McKenney et al., 2014; Schlager et al., 2014a; Schlager et al., 2014b; Splinter et al., 2012; Urnavicius et al., 2015). The DD-binding segment maps to the BICD2 N-terminal region, which forms an apparently uninterrupted dimeric coiled-coil of ≈250 residues that binds alongside the Dynactin filament. The BICD2 N-terminal region makes contacts near the barbed end, while a more C-terminal region of BICD2 makes contacts near the pointed end. This arrangement promotes binding to Dynein through several contacts with the N-terminal tail domain of the DHC as well as with the C-terminal region of the LICs (Celestino et al., 2019; Gama et al., 2017; Lee et al., 2020; Lee et al., 2018; Renna et al., 2020; Schroeder et al., 2014; Schroeder and Vale, 2016; Urnavicius et al., 2018; Urnavicius et al., 2015; Yeh et al., 2012).

In addition to the paradigmatic BICD2 adaptor and its family, several other proteins are or are likely to be activating adaptors, including CCDC88B, FIP3, HAP1, HOOK1-3, JIP3, Ninein and Ninein-like (NIN and NINL), NuMa, RAB11FIP3, RILP, Spindly (SPDL1), and TRAK1 (Gama et al., 2017; Hueschen et al., 2017; McKenney et al., 2014; Olenick et al., 2016; Redwine et al., 2017; Renna et al., 2020; Schroeder and Vale, 2016). At least some of these adaptors promote the interaction of Dynactin with two Dynein dimers, which increases processivity, speed, force production, and unidirectional movement (Grotjahn et al., 2018; Urnavicius et al., 2018). Collectively, the interactions of Dynein with Dynactin and adaptors appear to induce large conformational changes that align the motor domains for concomitant binding to microtubules (Zhang et al., 2017), an effect that likely extends to both Dynein dimers, when present.

While diverse, adaptors also share certain structural and functional features. These include 1) a propensity to form long coiled-coils with a dimeric parallel organization (Olenick and Holzbaur, 2019); 2) the ability of the N-terminal region to interact specifically with a conserved helix of LIC isoforms (Celestino et al., 2019; Gama et al., 2017; Lee et al., 2020; Lee et al., 2018). LIC-interacting sequences on adaptors belong to at least three different subfamilies, containing either a CC1 box, a HOOK domain, or EF-hand pairs (Olenick and Holzbaur, 2019; Reck-Peterson et al., 2018), which bind the LIC in different manners (Lee et al., 2020; Lee et al., 2018). The CC1 box (Figure 1A and Figure 1 – Supplement 1A) encompasses a highly conserved AAXXG sequence, where X denotes any amino acid (Gama et al., 2017; Hoogenraad and Akhmanova, 2016; Lee et al., 2018; Schlager et al., 2014b); 3) a second conserved motif, adjacent to the CC1 box and possibly also implicated in Dynein binding (Sacristan et al., 2018) (Figure 1A and Figure 1 – Supplement 1A). Even if it lies within the CC1 coiled-coil, this motif has been named the CC2 box (Sacristan et al., 2018). Here, we refer to it as the CC1’s second conserved motif (CC1 SCM); 4) a more C-terminal Spindly box motif (LΘXEΘ, where Θ indicates an aliphatic or aromatic side chain; Figure 1A and Figure 1 – Supplement 1B), conserved in many adaptors, that mediates binding to the four subunits of the Dynactin pointed end subcomplex (p25, p27, p62, Arp11) (Gama et al., 2017; Lau et al., 2021); 5) an extended stretch of coiled coil (corresponding to CC1), positioned between the CC1 box and the Spindly motif, that lodges between Dynein and the Dynactin mini-filament, strongly enhancing complex formation (Chowdhury et al., 2015; Urnavicius et al., 2018; Urnavicius et al., 2015); and 6) a C-terminal domain, positioned after the Dynein-Dynactin binding region, shown or hypothesized to bind to cargo (Akhmanova and Hoogenraad, 2015; Hoogenraad et al., 2003).

**Figure 1.**
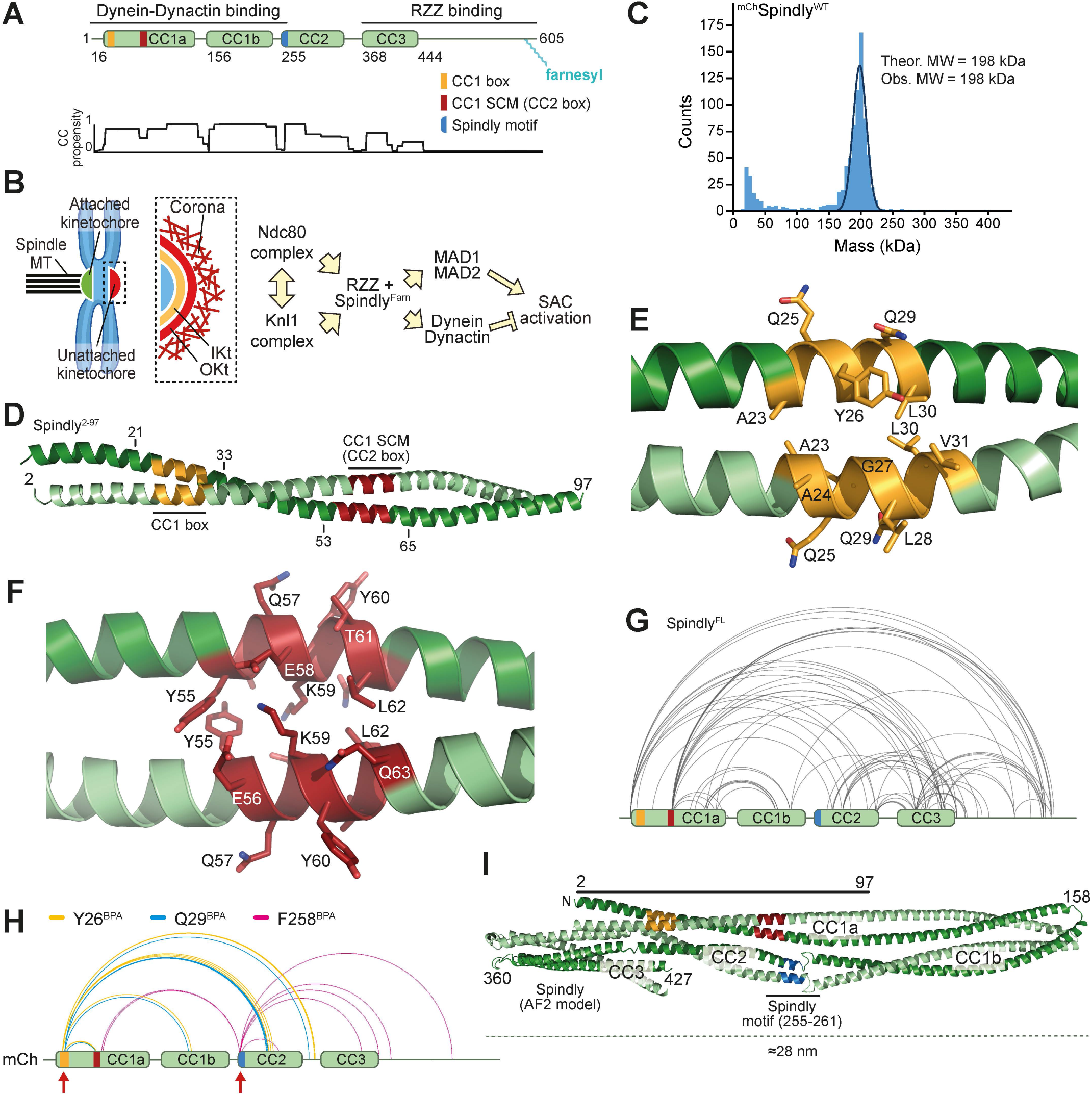
Spindly is a folded adaptor. (**A**) Schematic representation of the organisation of the coiled-coil regions of Spindly, and relevant coiled coil prediction (COILS, ExPaSy suite). (**B**) Organisation of the kinetochore and corona. (**C**) Results of mass photometry measurement of a sample of mCherry-tagged Spindly (^mCh^Spindly^FL^). The measurements are consistent with Spindly being a dimer in solution, even at very low concentration (10 nM). (**D**) Crystallographic structure of a Spindly^1-100^ construct. Only residues 2-97 were visible in the electron density. (**E**) Structure of the Spindly CC1 box (orange) and surrounding sequence. (**F**) Structure of the Spindly CC1 SCM (also known as CC2 box) (red) and surrounding sequence. (**G**) Summary of XL-MS data reporting Spindly intramolecular crosslinks. For ease of viewing, only crosslinks detected ≥3 times and involving sites ≥40 residues apart are depicted. See also Table S2 for a detailed list of all crosslinks. (**H**) Summary of XL-MS data reporting Spindly intramolecular crosslinks found through amber codon suppression experiments. Red arrows indicate the sites where the BPA residues were introduced. Crosslinking results from three mutants are merged: Y26BPA (orange), Q29BPA (cyan), and F258BPA (magenta). A few crosslinks identified between the BPA residues and the mCherry tag were considered spurious and not displayed. See also Table S2 for a detailed list of crosslinks. (**I**) AlphaFold2 Multimer prediction of Spindly structure. The CC1 box is in orange, the CC1 SCM in red, the Spindly motif in cyan. The C-terminal unstructured tail of Spindly (aa 440-605) was omitted from the model due to the very low confidence index (pLDDT) of the prediction for this region (unpublished results).

The exquisitely specific regulation of Dynein at different subcellular structures implies that the interaction with adaptors is tightly regulated, stimulated to occur only at the appropriate time and subcellular locale. Thus, in solution the majority of Dynein adopts the so-called phi-particle conformation, unable to interact with Dynactin and adaptors (Amos, 1989; Zhang et al., 2017). The Dynactin p150 subunit can dock onto the pointed end subcomplex, sterically preventing adaptor binding (Lau et al., 2021; Urnavicius et al., 2015). Similarly, the cargo-binding domains of several adaptors, and prominently of BICD2, have been shown to fold back onto the Dynein-binding domains, forcing an autoinhibited conformation that can be relieved either by cargo binding, or by removing the cargo-binding domain in recombinant protein (Hoogenraad et al., 2003; Liu et al., 2013; Splinter et al., 2012; Stuurman et al., 1999; Terawaki et al., 2015; Urnavicius et al., 2015).

Here, we have addressed the organization and regulation of the 605-residue adaptor Spindly (Figure 1A), a farnesylated mitotic regulator of DD. In early mitosis, Spindly promotes the recruitment of DD to kinetochores, the structures that connect chromosomes to spindle microtubules (Musacchio and Desai, 2017). This function of Spindly is enabled by the 800 kDa hexameric ROD-Zwilch-ZW10 (RZZ) cargo complex, to which Spindly binds directly through its farnesylated C-terminal region (Holland et al., 2015; Mosalaganti et al., 2017; Moudgil et al., 2015). At kinetochores, phosphorylation by the MPS1 kinase promotes the polymerization of the RZZ-Spindly (RZZS) complex into a mesh (Figure 1B), the kinetochore corona, which adopts a characteristic crescent shape and contributes to the initial phases of chromosome alignment that precede end-on microtubule attachment and chromosome bi-orientation (Kops and Gassmann, 2020; Magidson et al., 2015; Pereira et al., 2018; Raisch et al., 2021; Rodriguez-Rodriguez et al., 2018; Sacristan et al., 2018). Besides DD, the corona also recruits the MAD1:MAD2 complex (Barisic et al., 2010; Chan et al., 2009; Cheerambathur et al., 2013; Gassmann et al., 2008; Gassmann et al., 2010; Griffis et al., 2007; Rodriguez-Rodriguez et al., 2018; Starr et al., 1998; Yamamoto et al., 2008), a central component of the spindle assembly checkpoint (SAC), which synchronizes cell cycle progression with the completion of chromosome alignment (Kops and Gassmann, 2020; Musacchio, 2015). Upon achievement of end-on attachment, the DD-RZZS complex becomes activated and begins to travel away from the kinetochore and toward the spindle poles, in a process of corona disassembly referred to as “corona shedding” or “stripping” (Auckland et al., 2020; Basto et al., 2004; Howell et al., 2001; Mische et al., 2008; Sivaram et al., 2009; Varma et al., 2008; Williams et al., 1996; Wojcik et al., 2001). As the MAD1:MAD2 complex is transported away from kinetochores together with DD-RZZS, this also suppresses SAC signaling. Mutations in the Spindly motif abrogate kinetochore recruitment of DD, blocking concomitantly corona shedding and SAC silencing (Cheerambathur et al., 2013; Gassmann et al., 2010).

Spindly has all the sequence credentials of a *bona fide* DD activating adaptor (Gama et al., 2017) (Figure 1A); however, it activates DD only weakly in motility assays *in vitro* (McKenney et al., 2014). This suggests that Spindly may exist in autoinhibited and active forms, as its presence alone is not sufficient for DD activation. Previous studies have also highlighted interactions between the N-terminal and C-terminal domains on Spindly, supporting the idea of an autoinhibitory interaction that hinders DD binding (Mosalaganti et al., 2017; Sacristan et al., 2018). Here, we demonstrate that Spindly is subject to a complex regulation involving large conformational changes, and identify crucial intramolecular contacts that regulate Spindly autoinhibition and prevent Spindly interaction with DD. Our data suggest that relief of Spindly autoinhibition requires the interaction with the RZZ and an additional kinetochore trigger that can be bypassed mutationally. Our results describe an in-depth characterization of Dynein adaptor autoinhibition, and have important general implications for the mechanism of DD activation.

## Results

### Spindly adopts a complex dimeric structural organization

We have previously shown through solution scattering studies and hydrodynamic analyses that Spindly is an elongated molecule, and could be a dimer in solution (Mosalaganti et al., 2017; Sacristan et al., 2018). To conclusively show the stoichiometry of Spindly in solution, we employed mass photometry, which identified recombinant full-length Spindly (indicated as Spindly^FL^ or Spindly ^WT^) as a dimer, with an excellent agreement between theoretical and observed molecular masses (Figure 1C). We then determined the crystal structure of human Spindly^1-100^ (Table S1; only residues 2-97 were clearly resolved in the electron density), a fragment containing both the CC1 box and the CC1 SCM (Figure 1D-F). The structure demonstrated that Spindly forms a parallel dimeric coiled-coil, similar in its outline to that observed in structures of other adaptors captured in complex with DD (Lau et al., 2021; Urnavicius et al., 2018; Urnavicius et al., 2015). Thus, both Spindly^FL^ and an N-terminal segment of Spindly exist as stable dimers. The structure of Spindly^2-97^ is closely reminiscent of the structure of BICD2^1-98^ in complex with a peptide encompassing the LIC1 helix (residues 433-458, PDB ID 6PSE (Lee et al., 2020); Figure 1 – Supplement 1C). In Spindly, Ala23 and Gly27 in the CC1 box occupy *a* and *d* positions within the heptad repeats of the coiled-coil, similarly to Ala43 and Gly47 in BicD2, a pattern also conserved in BICDL1 and BICD1 (Figure 1 – Supplement 1A). This unusual composition for *a* and *d* residues generates a cavity along the BICD2 coiled-coil axis that interacts with aromatic and hydrophobic LIC side chains (Lee et al., 2020). Its conservation in Spindly was confirmed by high-confidence prediction by AlphaFold2 (AF2) in the variants Colabfold and AF2-Multimer (Evans et al., 2021; Jumper et al., 2021; Mirdita et al., 2021), which only became available during the final phases of this study (Figure 1 – Supplement 1D). Indeed, Spindly and BICD2 interact with the LIC with similar affinity (Lee et al., 2020; Lee et al., 2018).

Structural work on the complex of DD with various adaptors has demonstrated the existence of an uninterrupted coiled-coil spanning the distance between the LIC-binding CC1 box in BICD2 and a coiled-coil break that immediately precedes the Spindly motif. The coiled-coil, referred to as coiled-coil 1 (CC1), is cradled between Dynein and Dynactin (Urnavicius et al., 2018; Urnavicius et al., 2015). In Spindly, the coiled-coil propensity between the CC1 box and the break immediately preceding the Spindly motif (around residue 256) is generally high, but there is a conserved 2- or 3-residue insertion around residue 155 that maps to an interruption of the register of CC1 not expected in BICD1 and BICD2 (Figure 1 – Supplement 1E-F). The conservation of this feature in the Spindly family prompted us to investigate the possibility that the CC1 coiled-coil of Spindly splits into distinct segments (CC1a and CC1b). For this, we subjected full-length Spindly to crosslinking-mass spectrometry experiments with the bifunctional crosslinker DSBU (disuccinimidyl dibutyric urea) (Pan et al., 2018). This revealed multiple intramolecular contacts within Spindly, including “concentric” crosslinks of the putative CC1b region with the second half of CC1a, consistent with the idea that they may be arranged as an anti-parallel pair separated by a loop containing the conserved insertion around residues 154-155. In addition, we observed crosslinks of the first half of CC1a with CC2 and CC3, and of CC2 with CC3 (Figure 1G, Figure 1 – Supplement G and Table S2). Essentially identical crosslinks were observed in experiments with Spindly^1-440^ (lacking the flexible C-terminal region that contributes to binding the RZZ complex (Mosalaganti et al., 2017)), with the expected exception of contacts involving the C-terminal disordered region (Figure 1 – Supplement 1H-I).

These observations suggested that Spindly, at least in isolation, may adopt a compact conformation relative to the extended conformations observed for adaptors bound to DD (Urnavicius et al., 2018; Urnavicius et al., 2015). To probe this, we harnessed amber codon suppression to introduce the UV-photoactivatable crosslinker *p*-Benzoyl-L-phenylalanine (BPA) (Ai et al., 2011; Davis and Chin, 2012) into selected positions of an mCherry (mCh)-tagged construct of Spindly (^mCh^Spindly). These included Tyr26 and Gln29 (Y26^BPA^ and Q29^BPA^, respectively) in the CC1 box and Phe258 (F258 ^BPA^) in the Spindly box. After irradiation with UV light (Figure 1 – Supplement 1J-K), Y26^BPA^ and Q29^BPA^ generated largely equivalent crosslinking patterns, with a majority of targets near the center of the predicted CC2 coiled-coil, around residue 300 (e.g. K297, L298, Q299, I300, L303, and M306; Figure 1H, Table S2, and Figure 1 – Supplement 1L). F258^BPA^, on the other hand, crosslinked to residues E74 and L76 immediately after the CC1 SCM in the central half of CC1a, as well as to residues in CC3 and the tail (Figure 1H, Table S2).

Collectively, these results establish that Spindly is a dimer, and that it may fold as four interacting coiled-coil segments (CC1a, CC1b, CC2, and CC3), with extensive interactions of the first half of CC1a with CC2 and CC3, and of the second half of CC1a with CC1b. A high-confidence AF2 prediction rationalized these observations (Figure 1I and Figure 1 – Supplement 2). AF2 models depict Spindly as having a complex organization, where CC1 is almost invariably predicted to be interrupted around residues 154-155, giving rise to CC1a and CC1b coiled-coil segments. The ≈23 nm CC1a coiled-coil, partly captured in our crystal structure (Figure 1D), is predicted to encompass the majority of the long-axis of the Spindly dimer (≈28 nm), in good agreement with values derived from 2D class averages of negatively stained Spindly samples and small angle X-ray scattering (SAXS) experiments for both Spindly^FL^ and Spindly^1-440^ (Sacristan et al., 2018). CC1b packs against CC1a in such a way that the Spindly motif, located at the beginning of CC2, is positioned roughly halfway along the complex, facing the segment immediately C-terminal to the CC1 SCM, in excellent agreement with the BPA crosslinking experiments (Figure 1I and Figure 1 – Supplement 2). A loop around residue 360 separates CC2 and CC3, in such a way that both CC2 and CC3 are in contact with the first part of CC1. The CC1 box faces precisely the region centered on residue 300 identified by the crosslinking experiments with Y26^BPA^ and Q29^BPA^. Importantly, the model predicts that both the CC1 box and the Spindly motif will be largely inaccessible to DD. While all parallel coiled-coils are roughly symmetric, residues in the two chains experience different environments due to the asymmetric intramolecular interactions of the coiled-coils. Thus, the folded structure of Spindly predicted by AF2 is inherently asymmetric.

### Spindly is not accessible to Dynein-Dynactin

In addition to structural work demonstrating that the adaptors showcase an uninterrupted CC1 to DD, a non-comprehensive AF2 analysis of a subset of adaptors (including BICD2, BICDL1, HOOK1, HOOK3, Spindly, and TRAK1) confirmed that they all have an N-terminal extended dimeric coiled-coil section of variable lengths. In these models, the CC1 continues more or less uninterrupted until a break of variable length (where CC propensity drops). This precedes CC2, which usually begins with, or is even preceded by, the Spindly motif (Figure 2 – Supplement 1A). This is true also of BICD2 (Figure 2A), even if a coiled-coil prediction algorithm suggested a ≈30-residue drop in coiled-coil propensity after the CC1 SCM (Figure 1 – Supplement 1E). The AF2 models show that the CC1 in all these adaptors has a rather regular length of 35-39 nm. In other adaptors, including JIP3 and RILP, CC1 is considerably shorter (approximately 20 nm) and there is no obvious Spindly motif (Reck-Peterson et al., 2018). As established in Figure 1 and Figure 1 – Supplement 2, Spindly can also adopt a closed compact conformation, but its constellation of DD binding motifs predicts that it opens as a canonical adaptor under appropriate conditions. Indeed, AF2 predicted a closed conformation for Spindly^1-309^ (Figure 2B, *right* and Figure 1 – Supplement 2), while it did not predict convincing intramolecular interactions for Spindly^1-275^ (Figure 2B, *left* and Figure 1 – Supplement 2). This suggests that residues 276-309, a region that according to our XL-MS analysis faces the CC1 box, contains determinants of a conformational transition from a closed to an open form.

**Figure 2.**
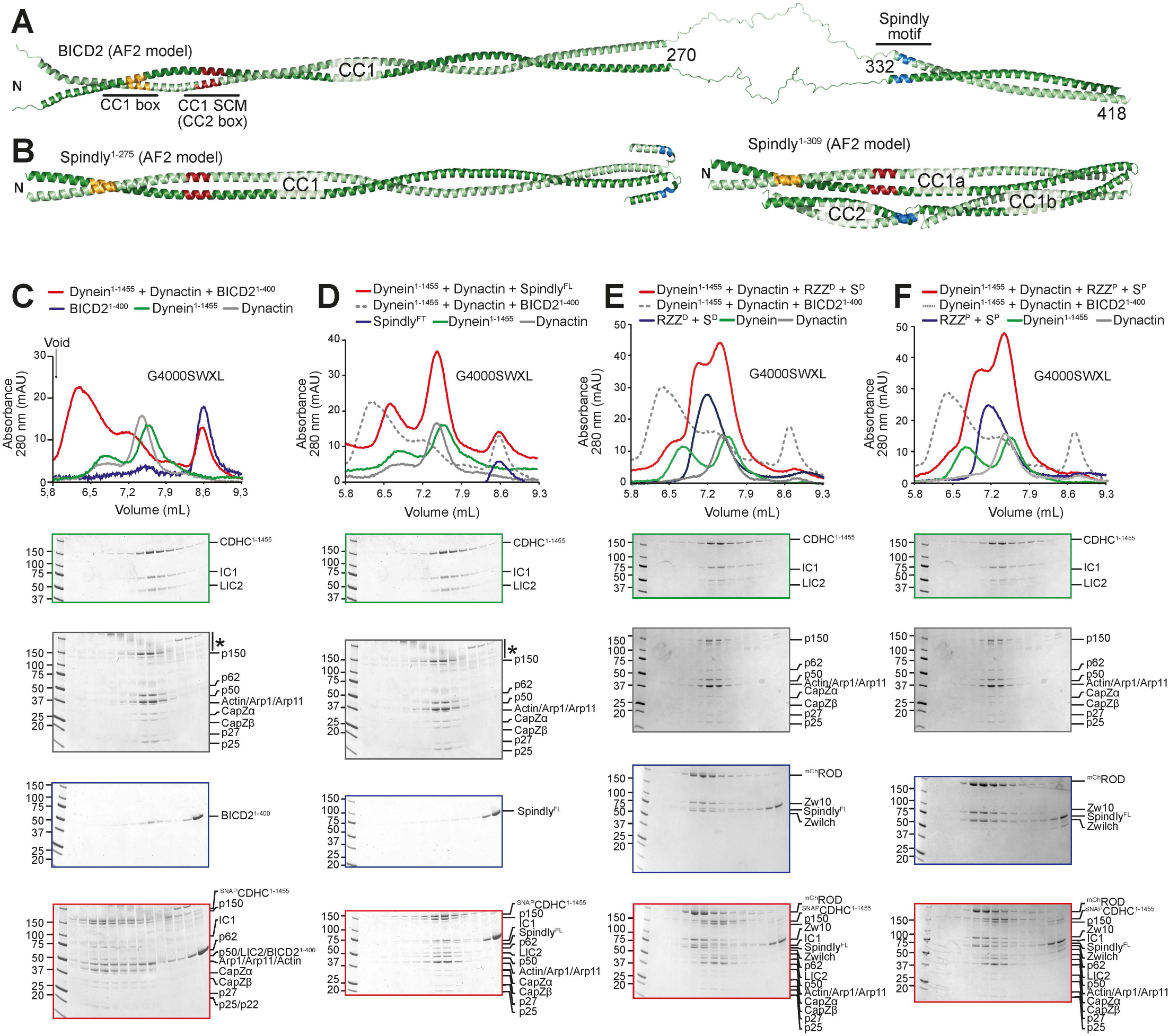
Spindly autoinhibition prevents its interaction with Dynein-Dynactin. (**A**) AlphaFold2 Multimer was used to predict a model of BicD2. The flexible region between 270 and 332 has very low reliability and has therefore been artificially linearized for visualization purposes (see Methods). The tail region has been omitted due to limited reliability of the predictions. The vertical like with an asterisk marks the accumulation of unknown contaminants in the upper part of the gel. (**B**) AlphaFold2 Multimer model of Spindly^1-275^. (**C**) AlphaFold2 Multimer model of Spindly^1-309^. The PAE plots and pLDDT scores are displayed in Figure 1 – Supplement 2. (**D-G**) Analytical size-exclusion chromatography on a G4000SWXL column (void volume ≈ 5.5 ml) to assess complex formation between Dynein tail (green), pig brain Dynactin (grey), and an adaptor -cargo/adaptor complex (blue), with the complex run shown in red. (**D**) BicD2^1-400^; (**E**) Spindly^FL^; (**F**) RZZ-Spindly^FL^ treated with λ-phosphatase; (**G**) RZZ-Spindly^FL^ pretreated with a mix of mitotic kinases (MPS1, Aurora B, CDK1/Cyclin B). Note that the Dynein tail and Dynactin controls are both shared between panels C and D, and panels E and F. In all shown SEC experiments, Spindly^FL^ was farnsylated.

In this context, a question of significant mechanistic relevance is whether binding of Spindly to its cargo is sufficient to relieve autoinhibition and trigger the formation of a complex with DD, as postulated for other adaptors (Olenick and Holzbaur, 2019; Terawaki et al., 2015). To address this question, we used analytical size-exclusion chromatography (SEC) to monitor the formation of complexes of DD with either Spindly or BICD2^1-400^, which served as positive control (Schlager et al., 2014a). Because the Dynein phi-particle might prevent Dynein from engaging into complexes with Dynactin and adaptors, we used a Dynein tail construct (CDCH^1-1455^) that does not form the phi-particle (Urnavicius et al., 2015; Zhang et al., 2017). As expected, Dynein, Dynactin, and BICD2^1-400^ interacted in a complex that eluted before any of the individual components, indicative of an increased Stokes’ radius (Figure 2C). Conversely, there was no evidence of binding of Dynein and Dynactin to full length Spindly (Figure 2D). This result is consistent with the hypothesis that Spindly adopts an auto-inhibited conformation refractory to interact with DD in the absence of adequate triggers.

The RZZ complex, to which farnesylated Spindly (Spindly^F^) binds directly, mediates Spindly’s kinetochore recruitment and can therefore be considered Spindly’s cargo, or a connector of Spindly to its chromosome cargo. We asked therefore if Spindly interacted with DD in presence of the RZZ complex. As the RZZ complex and Spindly are both known to be phosphorylated, with phosphorylation being critical for corona function (Raisch et al., 2021; Rodriguez-Rodriguez et al., 2018), we tested binding to DD after dephosphorylation (RZZ^D^ and Spindly^D^, Figure 2E), or after additional incubation with a mix of ATP and mitotic kinases, including CDK1/Cyclin B, MPS1, and Aurora B (RZZ^P^ and Spindly^P^, Figure 2F and Figure 2 – Supplement 1B). In either case, no interaction of the RZZS^F^ complex with DD was detected. Thus, the RZZ is not sufficient to relieve Spindly auto-inhibition and DD binding. Results presented below suggest that the kinetochore itself may play a role in the activation of Spindly required for DD binding.

### Relieving Spindly auto-inhibition

We next attempted to investigate what forces contribute to maintain Spindly’s autoinhibited state. The pointed end (PE) subcomplex of Dynactin, which interacts with the conserved Spindly motif, has previously been shown to bind adaptors with limited but measurable binding affinity (Gama et al., 2017; Lau et al., 2021; Yeh et al., 2012). Thus, we developed a minimal recombinant adaptor-binding subcomplex of Dynactin, containing only the subunits of the pointed end-capping complex, p25, p27, p62 and Arp11 (Figure 3A). After purification to homogeneity, we tested whether this pointed end (PE) complex interacted with different fragments of Spindly. In SEC experiments, the PE subcomplex did not bind ^mCh^Spindly^FL^, in agreement with the possibility that Spindly is auto-inhibited (Figure 3B). Conversely, Spindly^1-275^, a fragment that contains the Spindly box and that our AF2 predictions identified as having an open, elongated coiled-coil (Figure 2B, *left*), bound to the PE complex, as indicated by a clear, albeit only partial shift in its elution volume (Figure 3C). The interaction of Spindly^1-275^ with the PE complex required the Spindly motif, as Spindly^1-250^, a construct lacking it, was unable to bind the PE (Figure 3D). C-terminal deletions have been shown to relieve autoinhibition in BICD2, mimicking the effect of cargo binding (McKenney et al., 2014; Schlager et al., 2014b). Thus, we tested various Spindly C-terminal deletions for their ability to interact with the PE complex in the absence of cargo and other activators. Constructs lacking only the (disordered) C-terminal tail (Spindly^1-440^), or lacking the C-terminal tail and the CC3 (Spindly^1-354^) did not bind the PE complex in SEC experiments (Figure 3 – Supplement 1A), most likely because they adopt a closed conformation related to that predicted by AF2 for the Spindly^1-309^ construct (Figure 2B, *right*). We reasoned therefore that a Spindly deletion mutant lacking determinants of autoinhibition in the CC2 region ought to show features of the open complex and bind the PE complex even in the context of full-length Spindly. Indeed, an ^mCh^Spindly mutant lacking residues 276-306, a segment already identified for its interactions with the CC1 box, showed a clear level of binding to the PE complex (Figure 3E). Spindly^1-275^ and ^mCh^Spindly^Δ276-306^ did not fully co-elute with the PE complex, possibly indicative of low binding affinity. These constructs may be only partially open, or their binding site for the PE may be partly disrupted. We have also found that both constructs populate a dimer-tetramer equilibrium and that the tetramer, especially for Spindly^1-275^, may prevail at the micromolar concentration used in these experiments, possibly counteracting the interaction with the PE (unpublished observations and see Figure 4C,E, discussed below).

**Figure 3.**
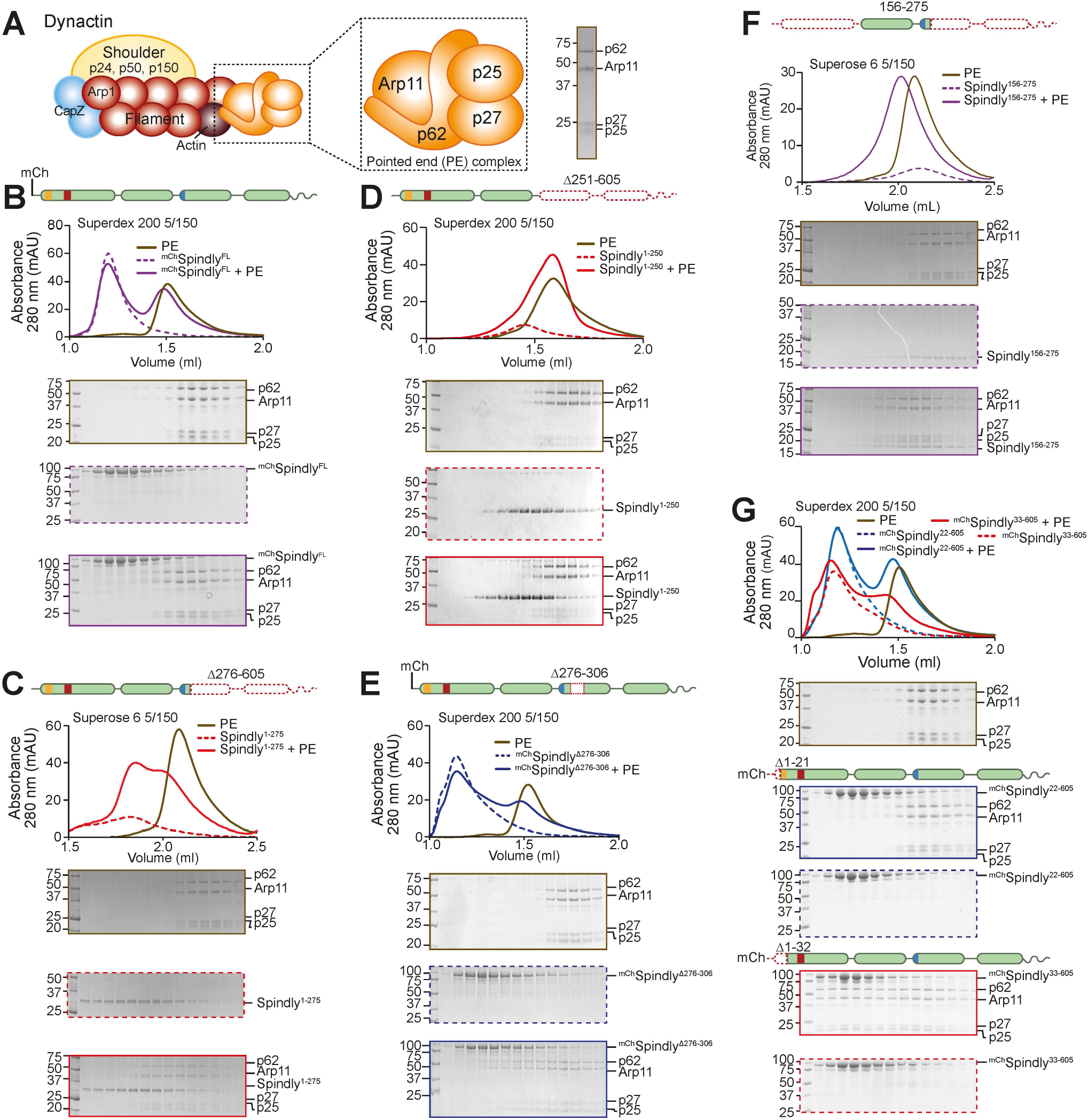
Spindly autoinhibition is relieved by N- and C-terminal deletions. (**A**) Schematic representation of the pointed end complex in the context of Dynactin, and SDS-PAGE of its chromatographic peak in gel filtration. (**B-G**) Analytical SEC binding assays between the Dynactin pointed end (brown) and Spindly constructs. The complex run is always represented with a continuous line, the Spindly construct with a dashed line. (**B**) ^mCh^Spindly (purple); (**C**) Spindly^1-275^ (red); (**D**) Spindly^1-250^ (red); (**E**) ^mCh^Spindly^Δ276-306^ (blue); (**F**) Spindly^156-275^ (purple); (**G**) ^mCh^Spindly^22-605^ (blue); ^mCh^Spindly^33-605^ (red). The control gels with the pointed-end (PE) alone are shared between panels B and G, C and F, D and Figure 3 – Supplement 1A.

**Figure 4.**
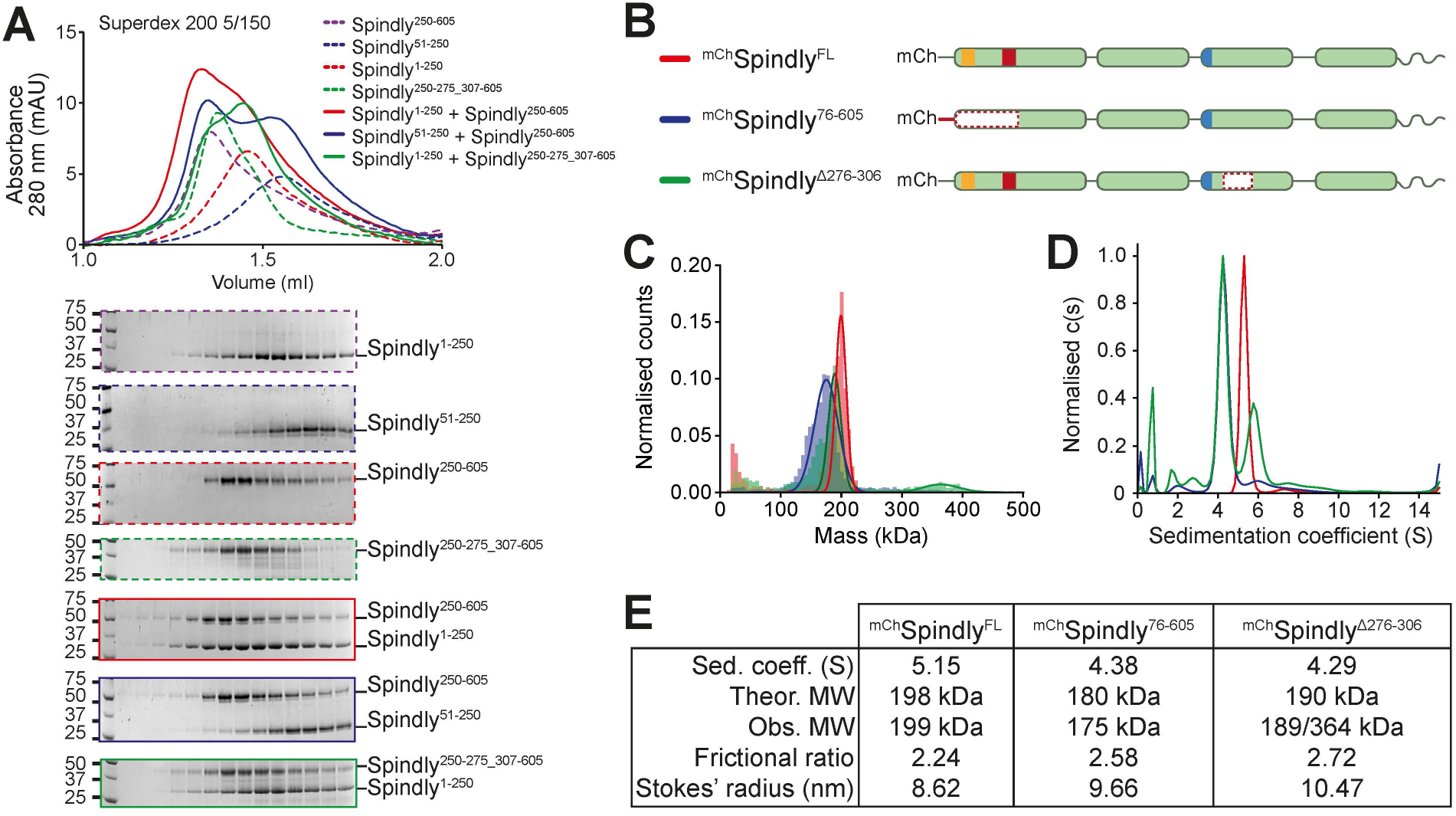
Spindly autoinhibition involves a direct interaction between N- and C-terminal regions. (**A**) Analytical SEC elution profile and SDS-PAGE analysis for interaction assays between the Spindly N-terminal and C-terminal domains. Spindly^1-250^ (red, dashed) interacts with Spindly^250-605^ (alone: purple, dashed; complex: red, continuous), but not with Spindly^250-275_307-605^ (alone: green, dashed; complex: green, continuous). Spindly^51-250^ (blue, dashed) does not interact with Spindly^250-605^ (complex: blue, continuous). This result shows that the Spindly regions 1-51 and 276-306 are required for Spindly folding. (**B**) schematic representation of Spindly constructs referred to in panels C,D and E. (**C**) Mass photometry results for the constructs in panel B. (**D**) Analytical ultracentrifugation results for the constructs in panel B. The smaller sedimentation coefficient indicates higher drag, which is caused by an increased Stokes’ radius. (**E**) Data table from hydrodynamic and mass photometry results in panels C-D. The Stokes’ radius and frictional ratio were estimated from the AUC-measured sedimentation coefficient and from the theoretical molecular weights.

We suspected that the N-terminal region of Spindly, which our experiments have suggested to be split in CC1a and CC1b segments, contributes to stabilize the autoinhibitory interaction that controls access of the PE complex to the Spindly motif. To test this idea, we asked if the ^mCh^Spindly^76-605^ construct, lacking the segment of CC1a where both the CC1 box and the CC1 SCM are located, bound the PE complex. Indeed, ^mCh^Spindly^76-605^ bound the Dynactin pointed end in SEC assays (Figure 3 – Supplement 1B), confirming that the N-terminal segment of Spindly contributes to maintain the autoinhibited state of Spindly. Finally, a mutant lacking the entire CC1a region and also truncated in the CC2 after the Spindly motif, Spindly^156-275^, interacted robustly with the PE complex (Figure 3F).

To identify the role of the CC1 box in this process, we designed two sequential N-terminal truncations. ^mCh^Spindly^22-605^, which retains the CC1 box, did not bind the PE complex. Conversely, ^mCh^Spindly^33-605^, which does not retain the CC1 box, bound the PE complex efficiently (Figure 3G). Two further short deletion mutants within the CC1, ^mCh^Spindly^Δ26-28^ and ^mCh^Spindly^Δ26-32^ respectively did not bind and bound weakly to the PE complex (Figure 3 – Supplement 1C-D), indicating that shorter deletions elicit a less penetrant effect on Spindly auto-inhibition. Collectively, these observations indicate that the Spindly auto-inhibition mechanism involves a tight intramolecular interaction of the conserved, Dynein-binding CC1 box with a regulatory segment of the Spindly CC2 coiled-coil roughly comprised between residues 276 and 306. Our results also imply that this inhibitory control cannot be readily relieved by DD, not even in presence of cargo (RZZ), possibly implying that an additional trigger at the kinetochore catalyzes opening.

### Testing structural predictions

The interactions between CC1 and CC2 in the AF2 model of Spindly in Figure 1I, together with our extensive analysis in Figure 3, explain why Spindly^1-250^ (CC1) and Spindly^250-605^ (CC2 and CC3) interact with high affinity and coelute from a SEC column (Sacristan et al., 2018), a result that we could readily reproduce (Figure 4A). When tested in the same assay, however, Spindly^51-250^, where the N-terminal segment of CC1a predicted to bind CC2 has been removed, did not interact with Spindly^250-605^. Similarly, Spindly^1-250^ seemed unable to bind Spindly^250-275_307-605^, where the CC2 segment predicted to face the CC1 box is deleted (Figure 4A). This is further supported by surface plasmon resonance experiments that showed that the region between residues 259 and 306 is essential for interaction with Spindly^1-250^ (Sacristan et al., 2018).

The disruption of interactions responsible for intramolecular folding may be expected to render the Spindly deletion mutants more elongated. To test this prediction, we first used mass photometry to verify that ^mCh^Spindly^76-605^ and ^mCh^Spindly^Δ276-306^ remained dimeric like wild type ^mCh^Spindly. This was confirmed for both constructs (with ^mCh^Spindly^Δ276-306^ being predominantly dimeric but also showing a tendency to form tetramers already at very low concentration, Figure 4C,E). We then analyzed the sedimentation behavior of these constructs by analytical ultracentrifugation, and derived their Stokes’ radii and frictional ratios (see Methods) (Figure 4D). For a given molecular mass, the product of the sedimentation coefficient and of the Stokes’ radius is a constant (Siegel and Monty, 1966). Indeed, the sedimentation coefficient of the deletion mutants was reduced in comparison to that of ^mCh^Spindly^FL^, indicative of larger Stokes’ radii and frictional ratios and therefore of a more elongated conformation (Figure 4E). Thus, deletion of regions in CC1a and CC2 predicted to interact with each other in the folded conformation of Spindly cause an at least partial opening of the Spindly structure.

### Opening up Spindly with point mutations

The region of Spindly downstream of the Spindly box (residues 281-322) is very conserved among Spindly orthologues, but not among other members of the BICD adaptor family (Figure 1 – Supplement 1L). Within this region, we mutated the positively charged residue pair R295-K297 to glutamate. The resulting construct, indicated as Spindly^CC2*^ (where CC2* indicates the R295E-K297E mutant in the CC2 coiled-coil) bound the PE complex in SEC (Figure 5A). The interaction with the PE complex was mediated by the Spindly motif, because combining the CC2* mutation with a mutation in the Spindly motif (F258A, indicated as SM*) to generate the ^mCh^Spindly^SM*-CC2*^ mutant, abolished the interaction (Figure 5B). In an orthogonal approach, we developed ^mCh^Spindly^ΔRV^, a construct deleted of the Spindly-specific 2-residue insert (residues R154-V155) between the CC1a and CC1b coiled-coil segments (Figure 1 – Supplement 1F) with the goal of favoring a full extension of CC1 like in BICD2, which lacks the insertion. ^mCh^Spindly^ΔRV^ bound the PE complex (Figure 5C), albeit more weakly, suggesting that that the 2-residue insertion into the CC1 of Spindly favors autoinhibition.

**Figure 5.**
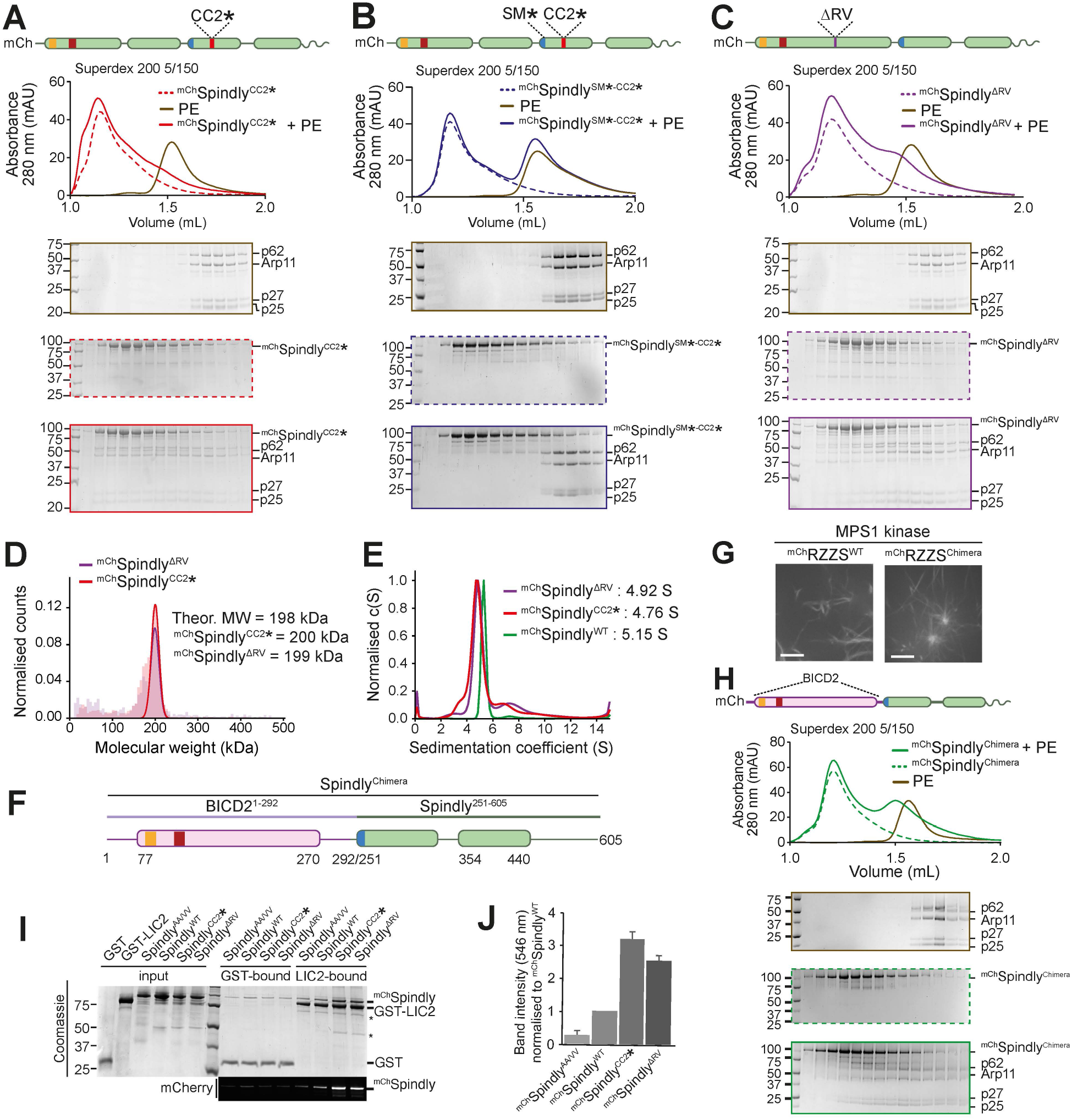
Point mutations relieve Spindly autoinhibition. (**A**-**C**, **H**) Analytical SEC analyses on a Superdex 200 5/150 column to assess complex formation between the Dynactin pointed end (brown) and various Spindly constructs. The complex run is always represented with a continuous line, the Spindly construct with a dashed line. (**A**) ^mCh^Spindly^CC2*^ (red); (**B**) ^mCh^Spindly^SM*-CC2*^ (blue); (**C**) ^mCh^Spindly^ΔRV^ (purple). (**H**) ^mCh^Spindly^Chimera^ (green). (**D**) Mass photometry results for ^mCh^Spindly^CC2*^ (red) and ^mCh^Spindly^ΔRV^ (purple). The main peaks’ “shoulders” are consistent with minor sample degradation. (**E**) AUC profile of ^mCh^Spindly^CC2*^ (red), ^mCh^Spindly^ΔRV^ (purple), and ^mCh^Spindly^WT^ (green). (**F**) schematic representation of the ^mCh^Spindly^Chimera^. (**G**) Spinning-disk confocal fluorescence microscopy-based filamentation assay at 561 nm shows the indicated ^mCh^RZZS species (4 µM RZZ, 8 µM Spindly) form filaments when incubated at 20°C with MPS1 kinase. Scale bar: 5 µm. (**I**) SDS-PAGE analysis of pulldown assay with either GST or GST-tagged LIC2 as bait, and mCherry-tagged Spindly as prey. Coomassie staining and fluorescent signal in the red channel are displayed. Asterisks mark contaminants or degradation products. (**J**) quantification of the ^mCh^Spindly fluorescent signal and SD calculated from three technical repeats. The PE alone controls in panels A and C are shared with the control in Figure 3E.

In mass photometry measurements, ^mCh^Spindly^CC2*^ and ^mCh^Spindly^ΔRV^ had masses expected of dimers (Figure 5D) and essentially indistinguishable from those of ^mCh^Spindly^FL^ (Figure 1C), suggesting that their ability to interact with the PE complex does not result from changes in stoichiometry. This was further confirmed by introducing a GST in Spindly^FL^ and Spindly^CC2*^ to reinforce dimerization of both. GST-Spindly^FL^ did not bind the PE complex, whereas GST-Spindly^CC2*^ did (Figure 5 – Supplement 1A-B). Analytical ultracentrifugation demonstrated a decreased sedimentation coefficient for ^mCh^Spindly^CC2*^ and ^mCh^Spindly^ΔRV^ (Figure 5E), indicative of a more extended conformation, as already shown for the Spindly deletion mutants in Figure 4D-E. Further, we developed an mCherry-tagged BICD2-Spindly chimeric construct that combined the BICD2 N-terminal region until the Spindly motif (residues 1-292^BICD2^), which is believed to contain an uninterrupted CC1, with the Spindly motif and C-terminal RZZ-binding domain of Spindly (residues 251-605^Spindly^; Figure 5F). Indeed, AF2 modelled this construct (Spindly^Chimera^) with a continuous CC1 until the flexible region that precedes the Spindly box (Figure 5 – Supplement 1C). Spindly^Chimera^ promoted Spindly-dependent oligomerization of the RZZ complex in filaments *in vitro* (Figure 5G), which requires MPS1 kinase and mimics kinetochore corona assembly (Raisch et al., 2021; Rodriguez-Rodriguez et al., 2018; Sacristan et al., 2018), thus indicating that the Spindly segment in Spindly^Chimera^ is sufficient for polymerization. In fact, Spindly^Chimera^ was more prone to promoting filaments than Spindly^WT^, and filaments were even observed at room temperature in absence of Mps1, when Spindly^WT^ did not stimulate filament formation (Figure 5 – Supplement 1D). In SEC experiments, ^mCh^Spindly^Chimera^ bound the PE complex (Figure 5H), in agreement with our expectation that an uninterrupted CC1 allows the PE to access the Spindly motif.

Finally, we asked whether relief of autoinhibition would increase the affinity of Spindly for Dynein in addition to increasing the affinity for the PE complex of Dynactin. As the CC1 box, required for the interaction with the Dynein LIC, is directly involved in the autoinhibitory interaction, we asked whether we could see increasing binding of LIC upon straightening Spindly to render the CC1 box more accessible. As the affinity of the LIC for CC1 box containing adaptors is too low for accurate study by SEC, we used a pull-down assay. We produced recombinantly a GST-tagged construct of the LIC2 isoform, and we tested its ability to pull down Spindly wild-type and “open” mutants. As a negative control, we used the ^mCh^Spindly^AA/VV^ (A23V-A24V) mutant, which has been previously shown to inhibit the interaction with the LIC2 in a similar assay (Gama et al., 2017). As open mutants, we used ^mCh^Spindly^CC2*^ and ^mCh^Spindly^ΔRV^. There was a clear increase of affinity for the LIC of the open mutants, with ^mCh^Spindly^CC2*^ showing an even slightly higher affinity than ^mCh^Spindly^ΔRV^, possibly due to an only partial restoration of coiled-coil continuity in the latter (Figure 5I-J), and in line with the lower apparent affinity of ^mCh^Spindly^ΔRV^ for the PE. Collectively, these results indicate that open mutants of Spindly interfering with the stability of the CC1-CC2 interaction or with the bending of the CC1 coiled-coil are more easily accessible to DD.

### Spindly^CC2*^ binds Dynein and Dynactin with higher affinity than Spindly^WT^

After showing that Spindly^WT^ does not interact with DD, we were eager to test whether the open Spindly mutants formed a super-complex with DD. For these assays, we were able to set up the production of recombinant human Dynactin, a significant technical challenge (Lau et al., 2021; Urnavicius et al., 2015). We produced Dynactin in the HEK293^expi^ expression system, using a modified version of the pBiG2 plasmid from the biGBac system (Weissmann et al., 2016) (Figure 6 – Supplement 1A). The resulting pBiG2 was used to transfect insect cells to produce a baculovirus and then to infect Expi293F cells (see Methods). The purified human Dynactin is biochemically pure and similar to the protein complex purified from pig brains, except for the absence of the p135 isoform of p150^glued^, which is not expressed in our system (a band in the same position is likely caused by degradation of the p150^glued^ subunits and is marked with an asterisk in Figure 6 – Supplement 1B). SEC combined with multiangle light scattering (SEC-MALS) measurements demonstrated that human recombinant purified Dynactin has the expected molecular mass (Figure 6 – Supplement 1C). Furthermore, recombinant Dynactin bound BICD2^1-400^ like pig brain Dynactin (Figure 6 – Supplement 1D-E). Finally, recombinant human Dynactin appeared morphologically indistinguishable from pig brain Dynactin, as judged by 2D classes from negative stain electron microscopy (Figure 6 – Supplement 1F; see also (Urnavicius et al., 2015)).

As we have shown that fragments from both Dynein (the LIC) or Dynactin (the PE) bind the CC2* mutant independently, we performed binding assays using stoichiometric ratios of Spindly, Dynein tail, and recombinant human Dynactin to maximize complex formation (Figure 6A). ^mCh^Spindly^WT^ was unable to form the super-complex with DD, as expected, but ^mCh^Spindly^CC2*^ and ^mCh^Spindly^Chimera^ were, as assessed by the shift of the adaptor into the expected super-complex peak (Figure 6A-B). We also found that ^mCh^Spindly^33-605^ interacted with Dynein-Dynactin, albeit apparently more weakly than ^mCh^Spindly^CC2*^ or ^mCh^Spindly^Chimera^. Both ^mCh^Spindly^CC2*^ and ^mCh^Spindly^Chimera^ were able to interact with the RZZ complex, indicating that the mutations do not affect the cargo-binding region of RZZ (Figure 6C-D).

**Figure 6.**
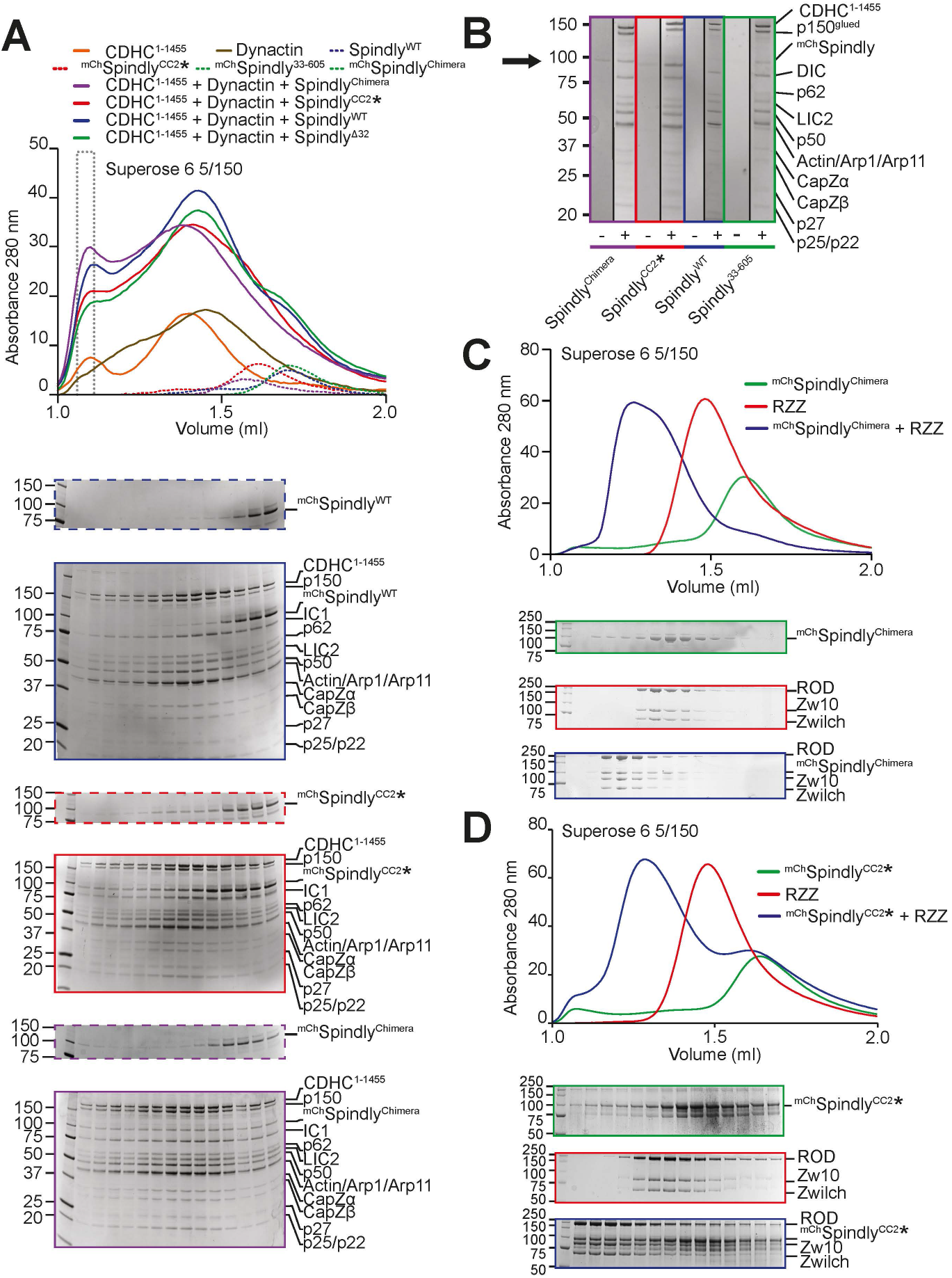
complex formation assay between Dynein-Dynactin and Spindly mutants. (**A**) Elution profiles and SDS-PAGE of complex formation assays between Dynein tail, recombinant Dynactin and Spindly constructs. Experiment run on a Superose 6 5/150 column, in stoichiometric conditions. Only selected gels are displayed. The gray dotted box indicates the fraction loaded in the SDS-PAGE shown in panel B. (**B**) Comparison of the fractions of the expected DDS complex peak shown in panel A. A minus sign indicates adaptor-only runs, a plus indicates full complex runs. The arrow points at the expected position of ^mCh^Spindly. (**C-D**) Analytical SEC experiments on a Superose 6 5/150 column to assess complex formation (blue) between the RZZ complex (red) and the indicated Spindly constructs (green). (**C**) ^mCh^Spindly^Chimera^. (**D**) ^mCh^Spindly^CC2*^.

### The localization of Spindly to human kinetochores

Finally, we assessed the ability of the different Spindly mutants to reach kinetochores in human cells arrested in mitosis with a spindle poison. Endogenous Spindly was depleted by RNA interference (RNAi) and recombinant ^mCh^Spindly variants were introduced in cells through electroporation as summarized in Figure 7 – Supplement 1A-B. As shown previously (Gassmann et al., 2010), depletion of Spindly prevented kinetochore recruitment of Dynactin (Figure 7A-D). Electroporation of ^mCh^Spindly^WT^ largely rescued these effects, but both the ^mCh^Spindly^CC2*^ and the ^mCh^Spindly^Chimera^ constructs showed strongly reduced kinetochore levels and, accordingly, greatly reduced levels of Dynactin, as measured with an antibody against the p150^glued^ subunit. The Spindly and Dynactin signals at kinetochores were well correlated for all four constructs (Figure 7E and Figure 7 – Supplement 1C-G). This argues that the reduction in kinetochore levels of Dynactin arises due to a reduction in the kinetochore levels of the ^mCh^Spindly^CC2*^ and ^mCh^Spindly^Chimera^ constructs, rather than to an inability of Dynactin to interact with them. This conclusion is also in line with our biochemical data showing that these Spindly mutants interact with Dynein and Dynactin. Because Spindly^Chimera^ promoted RZZ filamentation with enhanced efficiency *in vitro* (Figure 5 – Supplement 1D), we reasoned that its reduction at kinetochores may reflect the assembly of ectopic complexes with DD no longer limited to kinetochores, thus reducing the pool available for kinetochore binding. Indeed, ^mCh^Spindly^Chimera^ formed ectopic corona-like filaments with Dynactin in interphase cells in the absence of kinetochores (Figure 7F).

**Figure 7.**
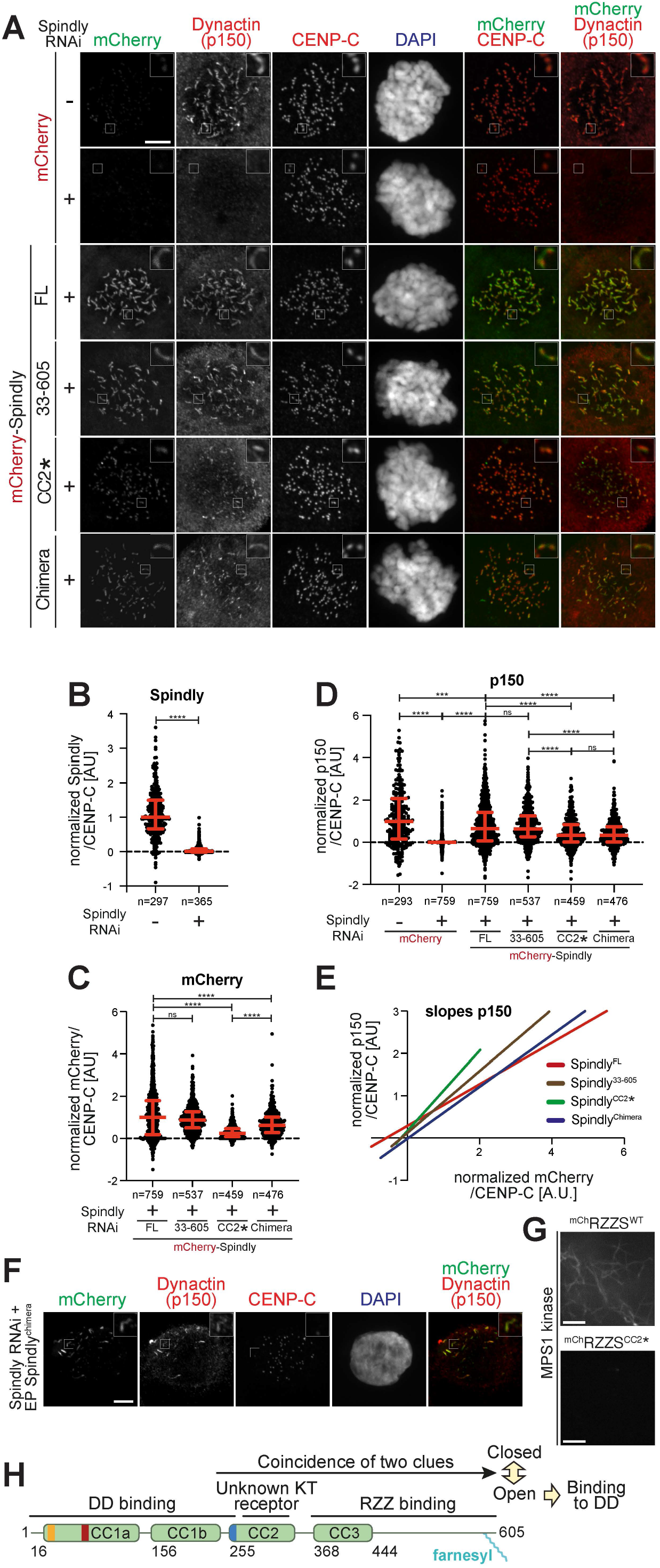
Kinetochore levels of Dynactin in presence of Spindly mutants. (**A**) Representative images showing the effects of a knockdown of the endogenous Spindly in HeLa cells on Dynactin recruitment monitored through the p150^glued^ subunit. RNAi treatment was performed for 48 h with 50 nM siRNA (see Figure 7 – Supplement 1A). Before fixation, cells were synchronized in G2 phase with 9 μM RO3306 for 16 h and then released into mitosis. Subsequently, cells were immediately treated with 3.3 μM nocodazole for an additional hour. CENP-C was used to visualize kinetochores and DAPI to stain DNA. Scale bar here and in panel F: 5 µm. (**B**) Quantification of residual Spindly levels at kinetochores. A representative image is shown in Figure 7 – Supplement 1B. (**C**) Quantification of kinetochore levels of the indicated electroporated ^mCh^Spindly proteins. (**D**) Kinetochore levels of Dynactin in cells depleted of endogenous Spindly and electroporated with the indicated Spindly proteins. (**E**) Least square linear fitting through the distribution of data points reporting for each kinetochore the CENP-C-normalized ^mCh^Spindly intensity on the X-axis and the CENP-C normalized p150^glued^ intensity on the Y-axis. The individual distributions are shown in Figure 7 – Supplement 1C-G. (**F**) Electroporated ^mCh^Spindly^Chimera^ is observed forming polymers in Spindly-depleted cells in interphase, causing ectopic recruitment of p150^glued^. (**G**) Spinning-disk confocal fluorescence microscopy-based filamentation assay at 561 nm with the indicated ^mCh^RZZS species (4 µM RZZ, 8 µM Spindly) at 20°C in presence of MPS1 kinase. Scale bar: 5 µm. (**H**) Model for the activation of Spindly.

Kinetochore localization by Spindly^CC2*^ was evidently severely impaired but this construct did not form ectopic corona-like filaments (unpublished results). Filamentation assays with RZZ showed it was almost unable to promote MPS1-dependent filament assembly (Figure 7G). Thus, it appears that the CC2* mutations (R295E-K297E) affect at the same time Spindly’s filament formation and kinetochore recruitment, without impairing DD binding. CC2* shares these characteristics with Spindly^Δ276-306^ (Figure 3E and (Raisch et al., 2021)). Collectively, these observations suggest that the region of CC2 containing these residues is at the same time implicated in kinetochore binding and filament formation, suggesting a possible mechanism of kinetochore regulation of corona expansion.

^mCh^Spindly^33-605^ not only decorated kinetochores to levels that were indistinguishable from those of ^mCh^Spindly^WT^, but also promoted recruitment of comparable levels of Dynactin (Figure 7A-D and Figure 7 – Supplement 1G). By suggesting that binding of the CC1 box to LIC1 is not required for robust DD recruitment at human kinetochores, this result may seem at odd with the observation that the Spindly^A23V^ mutant (where the CC1 box is mutated rather than absent) strongly impairs kinetochore recruitment of Dynactin (Sacristan et al., 2018). Our ^mCh^Spindly^AA/VV^ mutant could not be used to further investigate the issue, as – for unclear reasons – it was unable to reach kinetochores (unpublished results), preventing us from comparing it to ^mCh^Spindly^33-605^ in the same assay. Nonetheless, the new results with ^mCh^Spindly^33-605^ suggest that the deletion of the CC1 box or its mutation result in fundamentally distinct behaviors, and support a role of the LIC subunits as triggers of adaptor opening more than as decisive contributors to the binding affinity of the interaction, a speculative conclusion that will require further investigations.

## Discussion

Spindly has been mainly studied as a mitotic adaptor of Dynein-Dynactin (DD), but recent observations suggest that its role might extend to interphase cells (Del Castillo et al., 2020). Here, we have dissected a mechanism of conformational control that addresses the functions of Spindly at kinetochores, and that might inform future studies of Spindly in other cellular locales and cell cycle phases. More generally, our studies have implications for the control of DD in time and space. Adaptors are known to activate the motility and processivity of Dynein by stabilizing a complex with Dynactin (Hoogenraad and Akhmanova, 2016; McKenney et al., 2014; Schlager et al., 2014b). Studies so far have focused on N-terminal segments of adaptors that overcome intramolecular regulation, exemplified by the BICD2^1-400^ construct. Previous studies had demonstrated that the C-terminal cargo-binding region decreases the affinity of BICD2 for DD (Hoogenraad et al., 2003; Liu et al., 2013; Splinter et al., 2012; Terawaki et al., 2015). This effect was clearly evident also with Spindly, whose full-length form failed to bind DD in a variety of assays. However, we demonstrate that Spindly undergoes an apparently more complex regulation, that is not limited to cargo binding, but also includes a folded-back conformation of the N-terminal coiled-coil.

The RZZ complex, considered the Spindly cargo at kinetochores, binds directly to farnesylated Spindly (Mosalaganti et al., 2017; Raisch et al., 2021). The observation that binding of RZZ to Spindly was insufficient to unleash a conformational change compatible with DD binding motivated our investigation of the detailed mechanisms that hold Spindly in a closed conformation. A Spindly fragment encompassing residues 354-605 is sufficient to bind RZZ (Henen et al., 2021; Raisch et al., 2021), but the autoinhibited conformation requires a fragment of Spindly comprised between residues 276 and 309, and therefore positioned upstream of the minimal RZZ-binding region (Figure 7H). AF2 modeling (Evans et al., 2021; Jumper et al., 2021), which became available in the late phases of our work and proved to be instrumental as a validation tool, suggests that Spindly^1-275^ adopts an open conformation that is closely reminiscent of that of BICD2^1-400^, whereas Spindly^1-309^ may adopt a closed conformation. While these predictions must be taken with caution in the absence of supporting experimental evidence, they reinforce conclusions from our detailed biochemical and biophysical analysis, which are entirely consistent with this idea. Specifically, this model is supported by crosslinking analyses that predict the Spindly CC1 box to be in close proximity with residues 295-305, and by evidence that the closed conformation correlates with a break in the CC1 coiled-coil that allows Spindly to bend back on itself around residue 155. This model also explains a pattern of intra-molecular contacts, revealed by crosslinking-mass spectrometry, that would otherwise not be expected for a parallel coiled-coil like the one in Spindly. Our mutational analysis, combined with a binding assay monitoring the interaction with the pointed-end complex of Dynactin, and thus measuring the accessibility of the Spindly motif, was entirely consistent with the hypothesis that these two structural determinants, namely the 2-residue insertion in CC1 and the 295-305 region, are crucial for the auto-inhibitory mechanism of Spindly. This model was later confirmed by demonstrating binding of Spindly mutants to the entire Dynein-Dynactin complex. For these experiments, we established the expression of a recombinant form of Dynactin, a highly desirable development that will enable the production of significantly more homogeneous samples and the design of mutants, as well as enabling the use of tags for the individual subunits.

If RZZ binding is insufficient for unleashing the potential of Spindly to bind DD, what else might be required? Previous work has demonstrated that the RZZ complex is necessary for the recruitment of Spindly to the kinetochore and that Spindly, in turn, recruits DD (Barisic et al., 2010; Chan et al., 2009; Cheerambathur et al., 2013; Gassmann et al., 2008; Gassmann et al., 2010; Griffis et al., 2007; Raaijmakers et al., 2013; Starr et al., 1998; Yamamoto et al., 2008). Because the RZZ complex does not appear to be sufficient to promote binding of Spindly to DD, our analysis implies that a second, unknown interaction in the kinetochore elicits the opening of Spindly. The identity of the binding partner is unknown and will represent the focus of our future investigations. A possible working model is that the still unknown kinetochore binder interacts with the region comprised between residues 276-306, relieving it from its intra-molecular control of the Spindly closed conformation, and promoting the transition to the open conformation. Evidence supporting this idea is that mutations in the 276-306 region, including the deletion of this entire fragment or the introduction of charge-inverting point mutations at residues 295 and 297 respectively abolish or largely decrease the kinetochore recruitment of Spindly ((Raisch et al., 2021) and this study), implying that they impinge directly not only on the closed-open transition but also on the Spindly recruitment mechanism. *In vitro*, the 276-306 region is also required for the assembly of RZZ-Spindly filaments (this study and (Raisch et al., 2021)). However, this is unlikely to explain the kinetochore localization defect, because corona expansion is not required for robust recruitment of Spindly to the kinetochore (Raisch et al., 2021; Rodriguez-Rodriguez et al., 2018).

Our recent report of the cryo-EM structure of the RZZ complex (Raisch et al., 2021) represents a milestone of mechanistic studies of corona assembly at the kinetochore. We also investigated the mechanism of corona assembly and the role of Spindly in this process, identifying important roles for protein phosphorylation, in agreement with previous studies (Barbosa et al., 2020; Rodriguez-Rodriguez et al., 2018). Nonetheless, how Spindly promotes assembly of the corona, and the detailed organization of the corona, remain poorly understood. The RZZ is evolutionarily and structurally related to precursors of the coats that surround membrane vesicles during intra-cellular transport (Civril et al., 2010; Mosalaganti et al., 2017), and is thus likely to polymerize through similar mechanisms. Plausibly, the solution to this conundrum will require biochemical reconstitutions addressing the spectrum of interactions that this protein establishes at the kinetochore.

In conclusion, we have studied in mechanistic detail the basis of Spindly activation at the kinetochore and shown it to reflect a structural transition from a closed to an open conformation. We identified several structural determinants likely involved in the transition, and proposed a 2-step activation mechanism comprising RZZ binding and binding to an unknown second kinetochore receptor ultimately required for binding of Spindly to Dynein-Dynactin. These studies have important general implications for the mechanism of activation of adaptors, as they imply that the definition of “cargo” must be nuanced to every particular situation in which an adaptor is involved. In our particular analysis, the RZZ complex continues to be legitimately considered an element of the adaptor’s cargo, but our data imply that a second, equally important component remains to be identified. We speculate that a similar 2-step or multistep mechanism may apply to additional cargo-adaptor systems.

## Acknowledgments

We thank Giuseppe Ciossani, Stefan Raunser, and Thomas Surrey for helpful discussions, Dongqing Pan for help with crosslinking-mass spectrometry, Malte Metz for mass spectrometry data analysis, and Raphael Gasper-Schönenbrücher for help with biophysical experiments. This work was supported by the Max Planck Society, the Marie-Curie Training Network DivIDE (project number 675737), the European Research Council (ERC) through Synergy Grant 951439 (BIOMECANET), the Deutsche Forschungsgemeinschaft (DFG, German Research Foundation) through SFB1430 (Project-ID 424228829). Coordinates of Spindly^1-100^ will be deposited to the PDB upon publication. The authors declare no competing financial interests.

## Author contributions (following CRediT model)

**Conceptualization:** Ennio d’Amico, Andrea Musacchio, Anastassis Perrakis

**Funding acquisition:** Andrew Carter, Andrea Musacchio, Anastassis Perrakis

**Investigation:** Verena Cmentowski, Ennio d’Amico, Franziska Müller, Stefano Maffini, Misbha Ud Din Ahmad, Ingrid R. Vetter

**Project Administration:** Andrea Musacchio, Anastassis Perrakis

**Resources:** Andreas Brockmeyer, Andrew Carter, Petra Janning, Mathias Girbig, Sabine Wohlgemuth

**Supervision:** Andrew Carter, Andrea Musacchio, Anastassis Perrakis,

**Validation:** Andrea Musacchio, Anastassis Perrakis, Ingrid R. Vetter

**Visualization:** Verena Cmentowski, Ennio d’Amico, Andrea Musacchio, Ingrid R. Vetter

**Writing – original draft:** Ennio d’Amico, Andrea Musacchio

**Writing – review & editing:** All authors

## Materials and Methods

### Mutagenesis and cloning

cDNA segments encoding for wildtype Spindly or truncated constructs were subcloned into a pET28-mCherry plasmid, with an intervening PreScission cleavage site, or into a pLib plasmid for insect cell expression, with a 5’ insert coding for a His_6_-tag. Mutations were introduced by site-directed mutagenesis by Gibson assembly (Gibson et al., 2009). All constructs were sequence-verified. The BicD2-Spindly chimeric construct (^mCh^Spindly^Chimera^) was created by Gibson Assembly. We generated a sense primer containing the last 25bp of the coding sequence for the BicD2 (1-292) segment, and the first 25bp of the coding sequence for the Spindly (251-605) segment, as well as an antisense primer containing the same region. We used the former to expand the Spindly (251-605) sequence with an additional 3’ overlap with the MCS of a pLib-mCherry plasmid, and the latter to expand the BicD2 (1-292) coding sequence with an additional 5’ overlap with the pLib-mCherry plasmid MCS. The two PCR products were then assembled by Gibson Assembly into a pLib-mCherry plasmid opened by restriction cloning (BamHI-HindIII). A plasmid for the expression of the Dynactin pointed end was generated using the biGBac system (Weissmann et al., 2016). Coding sequences for Arp11, p62, p25-6His and p27 were subcloned into individual pLib plasmids, which were then used to build a pBiG1 plasmid, which was then used for expression of the entire complex. For the GST-LIC2 construct, cDNA encoding for human LIC2 was subcloned into a pLib vector containing an N-terminal TEV-cleavable GST.

The following Dynactin coding genes were codon-optimized for protein expression in insect cells and synthesized (Epoch Life Science): DCTN1 (Protein name p150, Uniprot Isoform 1, NCBI Reference Sequence NM_004082.4), DCTN2(p50, Isoform 1, NM_006400.4), DCTN3 (p24, Isoform 1, NM_007234.4), DCTN4 (p62, Isoform 1, NM_016221.3), DCTN5 (p25, Isoform 1, NM_032486.3), DCTN6 (p27, NM_006571.3), CAPZA1 (CapZα-1, NM_006135.2), CAPZB (CapZβ, Isoform 2, NM_004930.4), ACTR1A (Arp1, NM_005736.3), ACTR10 (Arp11, NM_018477.2), ACTB (β-actin, NM_001101.4). All genes were cloned into the pACEBac1 vector (Geneva Biotech) in between the polyhedron (polH) promoter and Simian virus 40 (SV40) poly A sequences by Gibson cloning. A sequence encoding the ZZ affinity tag plus a TEV-cleavage site was fused in-frame to the 5’-end of the DCTN1 gene. For cloning of mammalian cell expression vectors, the genes were cloned out of pACEBac1 into pcDNA™4/TO (Thermo Fisher Scientific) in between the CMV promoter and the BGH poly A signal. To assemble the dynactin genes into a single expression plasmid, biGBac cloning was used and adapted for mammalian expression. The biGBac cloning plasmids (pBig1a, pBig1b, pBig1c, pBig2abc) were generated by Gibson cloning and using the pACEBac1 vector as the backbone. To amplify the gene expression cassettes (GECs) by PCR and to assemble them into pBig1 plasmids, a set of CasMam oligonucleotides was designed and synthesized (Sigma-Aldrich/Merck, desalt purification). The oligonucleotides contained the optimized linker sequences (α/β/γ/δ/ε/ω) as described in Weissmann et al., 2016 and were modified to be complementary to the 5’-end of CMV promoter sequence (GTTGACATTGATTATTGACTAG – forward oligonucleotide) and reverse-complementary to the 3’-end of the BGH polyA sequence (CCATAGAGCCCACCGCATCC – reverse oligonucleotide). The GECs were then used to build three pBiG1 plasmids: pDCTN A, containing the ZZ-TEV-tagged p150^glued^, p50, and p24; pDCTN B, containing Arp1, Arp11, β-actin, CapZα, and CapZβ; and pDCTN C, containing p25, p27, and p62 subunits. The thee pBiG1 plasmids were then used to build a single pDCTN FL plasmid. Successful assembly of pDCTN FL was confirmed by complete plasmid sequencing (CCIB DNA Core Facility at Massachusetts General Hospital (Cambridge, MA)).

### Expression and purification of RZZ, mCherry-Spindly and Spindly constructs

The RZZ complex was expressed and purified using the biGBac system, with an mCherry N-terminally fused tag on the ROD subunit, as previously described (Sacristan et al., 2018). Expression of all mCherry-Spindly constructs and mutants except the ^mCh^Spindly^Chimera^ was carried out in *E. coli*. BL21 CodonPlus cells were transformed with the plasmid, and grown in TB at 37° C to an OD_600_ of 0.5. Expression was induced with 0.4 mM IPTG. The culture was then transferred into an incubator pre-cooled to 18° C, and grown overnight before harvesting. The pellet was then snap-frozen and stored at −80° C until purification. ^mCh^Spindly mutants containing the unnatural amino acid Bpa were expressed in *E. coli* BL21 strains containing the pEVOL-pBpF plasmid (Chin et al., 2002). Cells were cultured in selective (kanamycin, chloramphenicol) TB media, supplemented with 0.2% arabinose, to trigger expression of the tRNA synthetase/tRNA pair. Cells were grown at 37° C until an OD_600_ of 0.6 was reached. Expression was induced with 0.5 mM IPTG, and the Bpa was added to the bacterial culture at a concentration of 1 mM. The culture was then transferred into an incubator pre-cooled to 18° C, and grown overnight before harvesting. The pellet was then snap-frozen and stored at −80° C until purification. Spindly constructs without the mCherry tag and the ^mCh^Spindly^Chimera^ were expressed using the biGBac system as His_6_ fusions. Baculovirus was generated in Sf9 culture and used to infect TnAO38 cells, which were grown for 72 hours at 27° C before harvesting. The pellet was then snap-frozen and stored at −80° C until purification. All ^mCh^Spindly and Spindly constructs were purified using the same protocol. Pellets were resuspended in lysis buffer (50 mM HEPES pH 8.0, 250 mM NaCl, 50 mM imidazole, 2 mM TCEP) supplemented with protease inhibitor cocktail and lysed by sonication. The lysate was clarified by centrifugation followed by sterile filtration, and loaded onto a HisTrap HP column (Cytiva), which was then washed with at least 10 column volumes (CV) lysis buffer. Elution was performed with lysis buffer with 300 mM imidazole. The eluate was diluted 1:5 in no salt buffer (50 mM HEPES pH 8.0, 2 mM TCEP), and applied to a 6 ml Resource Q anion exchange column (Cytiva). Elution was then performed over a 50-500 mM NaCl gradient. Fractions of the peak were analysed by SDS-PAGE and those containing the protein of interest were pooled and concentrated. The concentrated sample was loaded onto a Superdex 200 10/300 pre-equilibrated in Spindly buffer (50 mM HEPES pH 8.0, 250 mM NaCl, 2 mM TCEP). The eluate was concentrated to 10 mg/ml, flash-frozen and stored at −80° C until use. The Spindly^250-275_307-605^ construct was expressed in bacteria as an mCherry fusion, and the mCherry tag was removed after the SEC purification step by overnight incubation with PreScission protease purified in-house. The protease and the cleaved mCherry tag were then separated from the Spindly sample by a further run of SEC on a Superdex 200 10/300 column pre-equilibrated in Spindly buffer, and the eluate was concentrated, flash-frozen and stored at −80° C until use.

### Expression and purification of Spindly^1-100^

The N-terminal 1-100 residues of Spindly were cloned in the pET-NKI-His-3C-LIC vector (Luna-Vargas et al., 2011) for expression in *E. coli* BL21 (DE3) cells. Cells were grown in LB medium at 37°C to an OD_600_ of 0.6. Overexpression was induced by adding isopropyl β-d-1-thiogalactopyranoside (IPTG) to a final concentration of 0.2 mM. Post induction, the cells were further grown for 18 hours at 18°C. For purification, cell pellet from a 2 l culture was resuspended in lysis buffer (40 mM HEPES/HCl pH 7.5, 500 mM NaCl, 10 mM Imidazole, 2 mM Tris(2-carboxyethyl)phosphine (TCEP, ThermoFisher)) supplemented with a protease inhibitor tablet (Roche) and 0.1 mM phenylmethylsulfonyl fluoride (PMSF). Cells were lysed by sonication (10 sec ON / 30 sec OFF; 70% Amplitude; 180 sec) and the lysate was centrifuged at 53000 g for 30 minutes. The supernatant was filtered through a 0.45 µM filter (Millipore) and incubated with 1 ml of Ni-Sepharose beads (Qiagen) on a rotator for 1 hour at 4°C. The protein was eluted from the beads with elution buffer (40 mM HEPES pH 7.5, 100 mM NaCl, 500 mM Imidazole, 2 mM TCEP. Protein containing fractions were pooled together and incubated with 3 mg/ml of 3C protease (1:100 molar ratio) overnight at 4°C to cleave off the N-terminal 6x-His tag. The protein was further purified by ion-exchange chromatography using a 6ml Porous XQ column and eluted with a linear NaCl gradient (50-1000 mM). As a final purification step, the protein was loaded onto a S200 10/300 SEC column equilibrated with buffer containing 40 mM HEPES/HCl pH 7.5, 100 mM NaCl, 2 mM TCEP. The elution fractions were analyzed by SDS-PAGE and the protein containing fractions were pooled together, concentrated to 11 mg/ml and stored at −80°C.

### Expression and purification of GST-LIC2

Baculoviruses for GST-LIC2 expression were generated in Sf9 culture and used to infect TnAO38 cells, which were grown for 72 hours at 27° C before harvesting. The pellet was then snap-frozen and stored at −80° C until purification. The pellet was resuspended in Spindly buffer supplemented with protease inhibitor cocktail, and lysed by sonication. The lysate was clarified by centrifugation followed by sterile filtration, and loaded onto a GSTrap column. The column was washed with 10 CV Spindly buffer. Elution was performed in SEC buffer supplemented with 50 mM glutathione, pre-buffered to pH 8.0. The eluate was pooled and concentrated for gel filtration. Gel filtration was performed on a Superdex 200 10/300 column pre-equilibrated in Spindly buffer. The fractions of the peak were analysed by SDS-PAGE, and the ones containing the protein of interest were pooled, concentrated, snap-frozen and stored at −80° C until use.

### Expression and purification of the Dynactin pointed end

Baculoviruses for Pointed end complex expression were generated in Sf9 culture and used to infect TnAO38 cells, which were grown for 72 hours at 27° C before harvesting. The pellet was then snap-frozen and stored at −80° C until purification. The pellet was resuspended in lysis buffer supplemented with protease inhibitor cocktail and 1 mg/ml DNAse, and lysed by sonication. The lysate was clarified by centrifugation followed by sterile filtration, and loaded onto a HisTrap HP column, which was then washed with at least 10 CV lysis buffer. Elution was performed with lysis buffer with 250 mM imidazole. The eluate was diluted 1:5 in no salt buffer, and applied to a 6 ml Resource Q anion exchange column. Elution was performed over a 50-500 mM NaCl gradient. Fractions of the peak were analysed by SDS-PAGE and those containing the entire complex were pooled and concentrated. The concentrated sample was loaded onto a Superdex 200 16/60 column pre-equilibrated in Spindly buffer. The eluate was concentrated to 15 mg/ml, flash-frozen and stored at −80° C until use.

### Expression and purification of recombinant Dynein tail

Expression of the Dynein tail (residues 1-1455) construct was performed using a previously described plasmid and protocol (Schlager et al., 2014a; Urnavicius et al., 2015). Pellets were resuspended on ice in Dynein buffer (50 mM HEPES-KOH pH 7.3, 150 mM KCl, 5 mM MgCl_2_, 2 mM TCEP, 0.2 mM ATP, 10% v/v glycerol) supplemented with protease inhibitor cocktail and 1 mg/ml DNAse, and lysed by sonication. The lysate was cleared by centrifugation and sterile filtration. The lysate was loaded on a 5 ml IgG column three times, and the column was washed with 20 CV Dynein buffer. Elution was performed by TEV cleavage of the ZZ tag overnight at 4° C. The cleaved protein was eluted in Dynein lysis buffer with fractionation. The eluate was pooled and concentrated for gel filtration. The concentrated sample was loaded on a Superose 6 10/300 column pre-equilibrated in Dynein buffer. The eluate from gel filtration was pooled and concentrated to approximately 5 mg/ml, snap-frozen, and stored at −80° C until use.

### Purification of Dynactin from pig brain

Pig brain Dynactin was purified essentially as previously described (Zhang et al., 2017). Pig brains were homogenised by blending at 4° C in a Waring blender, in PMEE buffer (35 mM PIPES-KOH pH 7.2, 5 mM MgSO_4_, 1 mM EGTA, 0.5 mM EDTA, 0.1 mM ATP, 2 mM TCEP), supplemented with 1 mg/ml DNAse and protease inhibitor cocktail. The lysate was cleared by two rounds of centrifugation, first at 30,000 g for 20 min at 2° C, then at 235,000 g for 45 min at 4° C, followed by two rounds of filtration, first through a glass fibre filter, then through a 0.45 µm filter. The cleared lysate was loaded on 300 ml SP Sepharose XL resin packed in an XK 50/30 column (Cytiva) and equilibrated in PMEE buffer. The column was washed with 4 CV PMEE. Elution was then performed with a 0 to 250 mM KCl gradient. Fractions containing Dynactin were initially identified by blotting for p150^glued^, pooled, and diluted 1:1 in PMEE. The diluted eluate was loaded on a MonoQ HR 16/10 column equilibrated in PMEE. The column was then washed with 10 CV PMEE. Elution was performed over a 150-350 mM KCl gradient. Dynactin eluted with a peak around 34 mS/cm conductivity. The fractions containing Dynactin were identified by SDS-PAGE, pooled, concentrated, snap-frozen, and stored at −80° C. As a final step of purification, the products of three MonoQ runs were pooled together and loaded on a Superose 6 10/300 column equilibrated in Dynactin buffer (25 mM HEPES-KOH pH 7.4, 150 mM KCl, 1 mM MgCl_2_, 2 mM TCEP, 0.1 mM ATP). The fractions of the peak were analysed by SDS-PAGE, and those containing Dynactin were pooled and concentrated to a final concentration of 3 mg/ml. The sample was snap-frozen and stored at −80° C until use.

### Expression and purification of recombinant human Dynactin

Recombinant human Dynactin was expressed using Expi293F cells (Invitrogen) and Expi293 expression medium (Invitrogen). Cells were freshly thawed for each expression, and passed at least three times before infection. All Expi293F cell cultures were performed at 37° C, 8% CO_2_, on an orbital shaker set to 125 rpm. Baculovirus for expression was made fresh for every expression. Sf9 cells were transfected with bacmid produced in EMBacY cells to produce baculovirus, which was then amplified through three 4-day rounds of amplification. The final V_2_ virus to be used for expression was made by infecting 60 ml V_2_ Sf9 culture supplemented with 5% FBS, and incubating it for 4 days at 27° C. The culture was then centrifuged at low speed to remove Sf9 cells, and the supernatant was decanted. 20 ml of sterile-filtered PEG solution (32% PEG6000, 400 mM NaCl, 40 mM HEPES pH 7.4) was added to the supernatant, mixed thoroughly, and the solution was allowed to precipitate overnight at 4° C in the dark. The precipitated virus was pelleted by centrifugation at 4000 rpm at 4° C for 30 minutes. The pellet was solubilized in 10 ml Expi293 expression medium prewarmed to 37° C and immediately used to infect 500 ml Expi293F culture, at a density of 3.5-5 x 10^6^ cells per ml. After 8 h from infection, 5 ml of a 1 M stock solution of sterile sodium butyrate in PBS was added to 500 ml Expi293F culture. The expression was incubated at 37° C, 8% CO_2_ for 48 h after infection. Cells were harvested by centrifugation at 350 g for 15 minutes, the pellet was washed with PBS, snap-frozen, and stored at −80° C until purification.

The pellet was resuspended in Dynein buffer, supplemented with 1 mg/ml DNAse and protease inhibitor cocktail, and lysed by sonication. The lysate was cleared by centrifugation, followed by sterile filtration, and loaded on a 5 ml IgG column. The flowthrough from the IgG column was further loaded three times on 4x 1 ml IgG beads in gravity flow columns. The column and the beads were washed with 10 CV each Dynein buffer. Elution was performed by TEV cleavage of the ZZ-tag overnight at 4° C. Fractions containing Dynactin were identified by SDS-PAGE, pooled and diluted 1:4 in MonoQ buffer (50 mM HEPES pH 7.3, 100 mM KCl, 5 mM MgCl_2_, 2 mM TCEP, 0.1 mM ATP, 10% glycerol). The diluted eluate was loaded on a MonoQ HR 16/10 column. After loading, the column was washed with 10 CV MonoQ buffer. Elution was performed over a 100-500 mM KCl gradient. Recombinant Dynactin eluted around 34 mS/cm conductivity, matching the results for pig brain Dynactin. The eluate was then concentrated in an Amicon 0.5 ml concentrator with a 100 kDa molecular mass cut-off to a final concentration of around 3-4 mg/ml. Typical yield from a 500 ml culture was in the range of 100-200 µg.

### In vitro dephosphorylation and phosphorylation

RZZ and Spindly were dephosphorylated using λ-phosphatase. Proteins were diluted to 10 µM in Spindly buffer, and λ-phosphatase was added to a final concentration of 500 nM, and incubated on ice for 15 minutes. Afterwards, MnCl_2_ was added at 10 mM concentration. The reaction mixture was incubated overnight at 10° C, and loaded on a Superose 6 10/300 column equilibrated in SEC buffer. The eluate was pooled, concentrated, snap-frozen and stored at −80° C until use. Pre-dephosphorylated RZZ and Spindly were phosphorylated with the mitotic kinases CDK1:CyclinB, MPS1, and Aurora B, which were all purified in-house (Huis In ’t Veld et al., 2021; Raisch et al., 2021; Sessa et al., 2005). Proteins were diluted to 10 µM in Spindly buffer, and kinases were added at a final concentration of 500 nM, and incubated on ice for 15 minutes. Afterwards, ATP was added at 1 mM, together with 10 mM MgCl_2_. The reaction mixture was incubated overnight at 10° C, and loaded on a Superose 6 10/300 column equilibrated in SEC buffer. The eluate was pooled, concentrated, snap-frozen and stored at −80° C until use.

### In vitro farnesylation

Farnesyltransferase α/β mutant (W102T/Y154T) was expressed and purified as previously described (Mosalaganti et al., 2017). Spindly was diluted to 100 µM in farnesylation buffer (50 mM HEPES pH 8.0, 250 mM NaCl, 10 mM MgCl_2_, 2 mM TCEP), and farnesyltransferase was added to a final concentration of 30 µM. Farnesyl pyrophosphate was added stepwise to a final concentration of 300 µM. The reaction mixture was incubated at RT for 6 hours, after which it was centrifuged at 16,000 g for 10 minutes to remove precipitate that formed during the reaction. The cleared reaction mixture was then loaded on a Superose 6 column equilibrated in SEC buffer to remove the farnesyltransferase. The fractions containing Spindly were identified by SDS-PAGE and pooled, concentrated, snap-frozen, and stored at −80 °C until use.

### Analytical Size Exclusion Chromatography (SEC)

Analytical SEC was performed under isocratic conditions at 4° C in Spindly buffer on an ÄKTAmicro system. Elution profiles were obtained by monitoring absorbance at 280 nm wavelength. 50 µl fractions were collected and analysed by SDS-PAGE. Complex formation assays were performed by mixing the samples at the indicated concentrations in 60 µl Spindly buffer and incubating them for at least 1 h on ice before the SEC assay was performed.

### Mass Photometry (MP)

Mass photometry experiments were performed essentially as described in Sonn-Segev et al., 2020. Standard microscope coverslips were cleaned with MilliQ water and isopropanol, and dried under an air stream. Silicon buffer gaskets were attached to the glass slides, and the slides were then mounted on a Refeyn TwoMP mass photometer (Refeyn Ltd). For measurement, Spindly construct samples were diluted to 100 nM in Spindly buffer immediately before measurement. The gasket was filled with Spindly buffer and the focal plane was automatically estimated. Proteins were diluted 1:10 into the buffer-filled gasket, to a final concentration of 10 nM. A 60 second movie was then recorded through AcquireMP (Refeyn Ltd). Data was processed using the DiscoverMP program. Contrast-to-mass calibration was performed with several known-mass proteins. Mass distributions were plotted with DiscoverMP and mean mass peaks determined by Gaussian fitting.

### Analytical Ultracentrifugation (AUC)

Sedimentation velocity analytical ultracentrifugation was performed at 42,000 rpm at 20° C in an An-60 Ti rotor (Beckman Coulter), using standard charcoal double-sector centrepieces. Protein samples were diluted in buffer containing 40 mM HEPES pH 8.0, 200 mM NaCl, and 2 mM TCEP. Approximately 300 radial absorbance scans at 587 nm were collected per each run, with a time interval of 1 minute. Buffer density and viscosity, as well as the protein partial specific volume, was estimated using the program SEDNTERP. Sedimentation peaks were identified through an analysis using the SEDFIT suite (Brown and Schuck, 2006), in terms of continuous distribution function of sedimentation coefficients (c(S)). Frictional ratios were manually set to match the sedimentation coefficient of the main peak to the theoretical mass of the sample, at the stoichiometry predicted by mass photometry data. A fitting run was then performed within SEDFIT, and the resulting estimations for frictional ratio and Stokes’ radius were collected as output. Figures were produced with the program GUSSI.

### Size-exclusion chromatography-multi angle light scattering (SEC-MALS)

SEC-MALS was performed using a Heleos II 18 angle light scattering instrument (Wyatt), which was coupled to an Optilab rEX online refractive index detector (Wyatt). 100 µl of pig-or human dynactin (0.15 mg/ml) were loaded onto a TSKgel G4000SWXL column with a TSKgel SWXL guard column (TOSOH Bioscience) equilibrated in GF150 buffer (25 mM HEPES pH 7.4, 150 mM KCl, 1 mM MgCl2, 5 mM DTT, 0.1 mM ATP) at room temperature. The column outlet was directly connected to the light scattering instrument and the refractive index detector. Data collection and determination of molecular weight was performed using the ASTRA 5.3.4 software (Wyatt).

### Negative stain electron microscopy (EM)

For negative stain EM, 400-square-mesh copper grids (Electron Microscopy Sciences) were plasma cleaned, and 3 µl of protein (≈100 nM in GF150 buffer) were applied to the grid. 20 µl of 2 % (w/v) uranyl acetate were added to the sample, and excessive stain was removed with a filter paper. Micrographs were recorded on an FEI Tecnai G2 Spirit transmission (120 kV) transmission electron microscope equipped with a Gatan Ultrascan 1000 XP CCD detector. Image acquisition was performed at 26000x nominal magnification and 1.5 µm underfocus. For 2D classification of recombinant human dynactin, approximately 5000 particles were picked semi-automatically using EMAN2 (Tang et al., 2007) and 2D-classified using RELION 2.1 (Scheres, 2012).

### DSBU and UV crosslinking

For DSBU crosslinking, all proteins in the assay were diluted to 5 µM in Spindly buffer. DSBU was added at a final concentration of 3 mM, and the reaction was incubated for 1 hour at RT. At the end of the reaction, TRIS-HCl pH 8.0 at a final concentration of 100 mM was added and the solution was thoroughly mixed to quench the reaction. Successful crosslinking was evaluated by SDS-PAGE analysis. For UV crosslinking, the BPA-containing Spindly mutants were diluted in Spindly buffer to a final concentration of 5 µM. The samples were irradiated with LED UV light at 365 nm wavelength for 15 minutes to induce complete cross-linking. Successful crosslinking was evaluated by SDS-PAGE analysis.

### Processing for LC/MS

Crosslinked samples were precipitated by dilution into 4 volumes of acetone pre-cooled to −20° C, followed by overnight incubation at −20° C. Pellets were resuspended and denatured in 8 M urea, 1 mM DTT, and alkylated with 5.5 mM chloroacetamide. The concentration of urea was then lowered to 4 M by dilution in 20 mM ammonium bicarbonate, pH 8.0, and the protein solution was then digested overnight at room temperature with Trypsin. Digestion was stopped by adding TFA to a final concentration of 0.2%. Samples were subjected to SEC on a Superdex 30 Increase 3.2/300 (Cytiva), followed by purification on a tC18 Sep-Pak column (50 mg, Waters) as previously reported (Pan et al., 2018). For the ^mCh^Spindly^Y26BPA^ and ^mCh^Spindly^Q29BPA^, the SEC step was omitted.

### LC-MS/MS analysis

LC-MS/MS analysis was performed as described in Pan et al., 2018. Data analysis was performed for the DSBU-crosslinked samples in MeroX 2.0.0.8. The analysis of Bpa-crosslinked samples was performed in MeroX 2.0.1.4. An artificial amino acid “Z” was added with the composition C16H13NO2 (mass 251.09463), and Bpa was defined as a crosslinker with parameters: ‘Specificity site 1:Z’, ‘Specificity Site 2: [ABCDEFGHIKLMPQRSTVWY]’, and ‘Maximum Cα-Cα -distance: 30 Å’. For the Spindly^F258BPA^ sample, the analysis was performed within MeroX 2.0.1.4 with a False Discovery Rate (FDR) of 50%. Crosslinks were then exported into XiView (Graham et al., 2019), and crosslinks with a score lower than 50 were filtered out. Visual representations of crosslinking data were produced using the xVis website (https://xvis.genzentrum.lmu.de (Grimm et al., 2015)), and then edited to produce the final figures.

### Molecular modelling and bioinformatical methods

AlphaFold 2 (AF2) was used for all molecular modelling (Jumper et al., 2021). For the Spindly^1-275^, Spindly^1-309^, and Spindly^1-440^ constructs, the original version of AlphaFold Multimer was used (Evans et al., 2021). For all other constructs the ColabFold version of AlphaFold 2 was used (Mirdita et al., 2021). A general feature of AF2 Multimer is the higher sensitivity to intra- and intermolecular interactions compared to ColabFold and thus a tendency to predict extended protein structures as a more or less compact model, making it difficult to distinguish artificial contacts from structurally significant ones. This problem is worse for fragments containing long, disordered stretches. Therefore, the disordered C-terminal region of Spindly (441-605) was not included in the predicted sequences, unless otherwise specified. In absence of clear predicted intramolecular contacts from the PAE plots, the models of the adapters shown in Figure 2 - Supplement 1 (obtained with ColabFold) were manually “unfolded” with PyMol and Coot to extend them and make them comparable. To determine the putative interactions in the autoinhibited Spindly fragments, the pLDDT scores of the models and especially the “predicted alignment errors” (PAE) were scrutinized to see which parts of the models were predicted to have defined relative orientations to each other (Figure 1 – Supplement 2). Also, some features, like the “kink” at Spindly residues 155-156, were very consistently predicted in almost all models. Other features, like predicting extended conformations of e.g. the Spindly N-terminal coiled coil, were more or less frequent, depending on the presence of the interacting, autoinhibitory regions, which was interpreted as being indicative of the strength of the co-evolutionary signal present in those regions.

### Crystallographic structure of Spindly^1-100^

Crystallization trials were set up in MRC 2-well sitting drop plates by mixing the protein and the reservoir solution in a 1:1 ratio. Initial crystallization hits were optimized and final crystals were obtained in a condition containing Bis-Tris Propane/HCl pH 6.5, 19% PEG3350 and 0.2M KSCN. For data collection, crystals were cryoprotected in mother liquor containing 20% glycerol prior to flash cooling in liquid N_2_. Data for native crystals were collected at the MASSIF-1 beamline (Bowler et al., 2015) at the ESRF, Grenoble. For SelenoMethionine protein production, *E. coli* BL21 DE3 cells were grown in ready to use SelenoMethionine medium (Molecular Dimensions) and purified using the same protocol as for the native protein. SeMet crystals could be reproduced in similar crystallization conditions, and anomalous data were collected at I04 beamline of Diamond Light Source. Details for all steps are available in Table S1. Data processing was done with XDS (Kabsch, 2010) and the structure was solved by Se-SAD using the CRANK2 (Pannu et al., 2011) pipeline from the CCP4 (Winn et al., 2011) software suite. The initial model contained 206 residues built in 8 fragments and clearly showed the parallel dimeric coiled coil. Further model building was performed in Coot (Emsley et al., 2010) and subsequent refinement cycles were carried out in REFMAC (Murshudov et al., 2011) and PDB-REDO (Joosten et al., 2014) using non-crystallographic and jelly-body restraints. The structure was refined to 2.8 Å resolution to an R_free_ of 29%. All residues are in the favorable regions of the Ramachandran plot and the structure is in the 97^th^ percentile of Molprobity (Williams et al., 2018).

### Cell culture and drug treatment

HeLa cells were grown in Dulbecco’s Modified Eagle’s Medium (DMEM; PAN Biotech) supplemented with 10 % tetracycline-free FBS (PAN Biotech), and L-Glutamine (PAN Biotech). Cells were grown at 37°C in the presence of 5 % CO2.

### Cell transfection and electroporation

Depletion of endogenous Spindly was achieved through reverse transfection with 50 nM Spindly siRNA (5′-GAAAGGGUCUCAAACUGAA-3′ obtained from Sigma-Aldrich, (Gassmann et al., 2010)) for 48 hours with RNAiMAX (Invitrogen, Carlsbad, California, United States). For rescue experiments, 24 hours after Spindly depletion, we electroporated recombinant Spindly constructs labeled with an N-terminal mCherry, at a concentration of 7 μM in the electroporation slurry (as previously described in (Alex et al., 2019)) (Neon Transfection System, Thermo Fisher). Control cells were electroporated with mCherry. Following an 8 hour recovery, cells were treated with 9 µM RO3306 (Calbiochem) for 15 hours. Subsequently, cells were released into mitosis in presence of 3.3 µM Nocodazole (Sigma) for 1 hour.

### Immunofluorescence

Cells were grown on coverslips pre-coated with Poly-L-lysine (Sigma-Aldrich). Cells were fixated with 4% PFA in PHEM for 10 minutes. Subsequently, the cells were permeabilized for 10 minuets with PHEM supplemented with 0.5% Triton-X100 (PHEM-T). After blocking with 5% boiled goat serum (BGS) in PHEM buffer, the cells were incubated for 2 hours at room temperature with the following primary antibodies: CENP-C (guinea pig, MBL, #PD030, 1:1000), Dynactin-p150 (mouse, BD Trans. Lab., #610473, 1:400), Spindly (rabbit, Bethyl, A301-354A, 1:1000) diluted in 2.5 % BGS-PHEM-T (PHEM supplemented with 0.1% Triton-X100). Subsequently, cells were incubated for 1 hour at room temperature with the following secondary antibodies: Goat anti-mouse Alexa Fluor 488 (Invitrogen, A11001), donkey anti-rabbit Rhodamine Red (Jackson Immuno Research 711-295-152), goat anti-guinea pig Alexa Fluor 647 (Invitrogen, A-21450). All washing steps were performed with PHEM-T buffer. DNA was stained with 0.5 μg/ml DAPI (Serva) and Mowiol (Calbiochem) was used as mounting media.

### Cell imaging

Cells were imaged at room temperature using a spinning disk confocal device on the 3i Marianas system equipped with an Axio Observer Z1 microscope (Zeiss), a CSU-X1 confocal scanner unit (Yokogawa Electric Corporation, Tokyo, Japan), 100 × /1.4NA Oil Objectives (Zeiss), and Orca Flash 4.0 sCMOS Camera (Hamamatsu). Images were acquired as z sections at 0.27 μm (using Slidebook Software 6 from Intelligent Imaging Innovations or using LCS 3D software from Leica). Images were converted into maximal intensity projections, exported, and converted into 16-bit TIFF files. Automatic quantification of single kinetochore signals was performed using the software Fiji with background subtraction. Measurements were exported in Excel (Microsoft) and graphed with GraphPad Prism 9.0 (GraphPad Software). Statistical analysis was performed with a nonparametric t-test comparing two unpaired groups (Mann-Whitney test). Symbols indicate: n.s. = p > 0.05, ∗ = p ≤ 0.05, ∗∗ = p ≤ 0.01, ∗∗∗ = p ≤ 0.001, ∗∗∗∗ = p ≤ 0.000

### GST-LIC2 pulldown assay

GST and GST-LIC baits were added to 10 µL GSH beads (Serva) at a final concentration of 4 µM in Spindly buffer, in Pierce micro-spin columns (Thermo Scientific), and incubated at 4° C for 1 h. The unbound baits were removed by centrifugation, and the beads were washed twice. The Spindly preys were added at a final concentration of 8 µM in Spindly buffer. The bait-bound beads were incubated with the preys at 4° C for 1 h, after which the supernatant was removed, and the beads washed twice. After the final wash, both prey and bait were eluted in Spindly buffer supplemented with 50 mM glutathione. The inputs and eluates were analysed by SDS-PAGE, and the unstained gels were imaged in the 546 nm channel. The signal from the main band was quantified using ImageLab (Biorad), in comparison to the signal of the SpindlyFL band. The gels were then stained with Coomassie and imaged.

### RZZS filaments

RZZS filaments were formed and imaged essentially as described in (Raisch et al., 2021). 4 µM RZZ was incubated with 8 µM prefarnesylated Spindly in presence of 1 µM MPS1, in M-buffer (50 mM HEPES pH 7.5, 100 mM NaCl, 1 mM MgCl2, 2 mM ATP, 1 mM TCEP) at room temperature overnight. Flow chambers were assembled by placing two parallel strips of double-sided tape onto a glass slide, with a standard coverslip on top, creating a chamber of 5 to 10 µl volume. The filament sample was diluted to 0.5 µm and loaded into the flow chamber. Imaging was performed on a 3i Marianas system at 100X magnification in the 561 nm channel. Sample images were acquired as 5-stacks of z-sections at 0.27 µm, converted into maximal intensity projections, and processed in Fiji (Schindelin et al., 2012).

---

**Figure 1 – Supplement 1.**
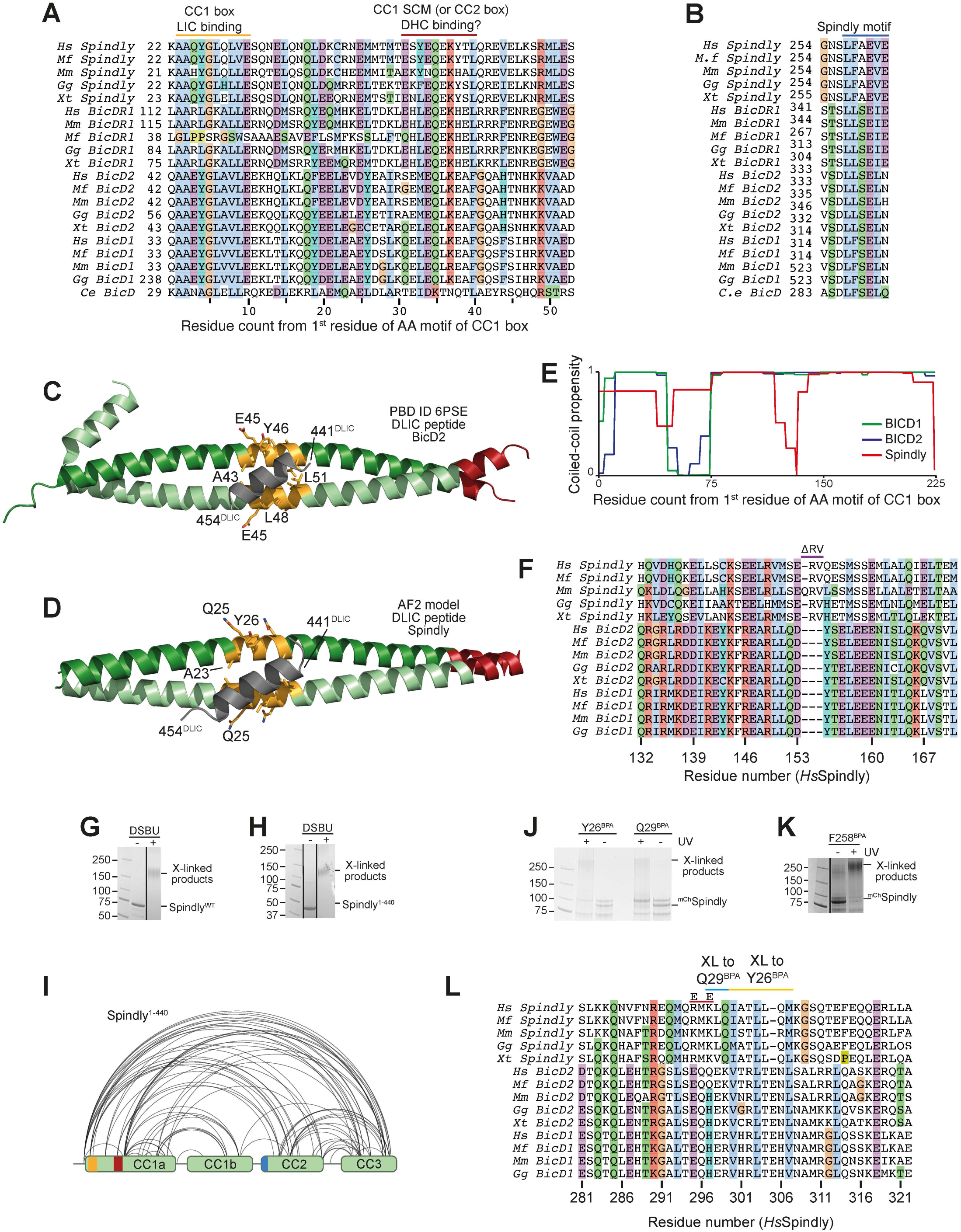
Additional analyses of Spindly motifs and their influence on Spindly conformation. (**A**) Multiple sequence alignment of the first part of the CC1 region of the indicated adaptors containing the CC1 box and the CC1 SCM (or CC2 box). Hs, Homo sapiens; Mf, Macaca fascicularis; Mm, Mus musculus; Gg, Gallul gallus; Xt, Xenopus tropicalis; Ce, Caenorhabditis elegans. (**B**) Multiple sequence alignment of the Spindly motif. (**C**) Cartoon model of PDB ID 6PSE (Lee et al., 2020) showing the mode of binding of a LIC peptide (grey) to the CC1 box (yelloworange). (**D**) ColabFold prediction model of the Spindly CC1:LIC peptide complex. Coloring as in panel C. (**E**) Coiled-coil propensity was measured with the COILS program within ExPasY suite (Duvaud et al., 2021) and displayed for all indicated adaptors from the first residue of the CC1 box (see panel A). The coiled-coil propensity for Spindly has a deep that corresponds to a 2-residue insertion shown in panel F. (**F**) Multiple sequence alignment of the region of CC1 around the 2-residue insertion in Spindly that causes a deep in the coiled-coil prediction profile (see panel E). (**G**) SDS-PAGE documenting crosslinking of the full-length Spindly proteins with DSBU. (**H**) SDS-PAGE documenting crosslinking of the Spindly^1-440^ proteins with DSBU. Panel G and H were obtained from the same original gel and the marker lane is the same in the two panels. (**I**) Summary of XL-MS data reporting Spindly intramolecular crosslinks. for ease of viewing, only crosslinks detected ≥3 times and involving sites ≥40 residues apart are depicted. See also Table S2. (**J**-**K**) Coomassie-stained SDS-PAGE gels documenting crosslinking of the indicated BPA mutants upon treatment with UV light. (**L**) Multiple sequence alignment of the indicated adaptors in the main region targeted by Q29^BPA^ and Y26^BPA^. The CC2* mutant discussed in the text is the charge reversal mutant (E-E) at the two indicated positively charged residues (R295 and K297 in human Spindly).

**Figure 1 – Supplement 2.**
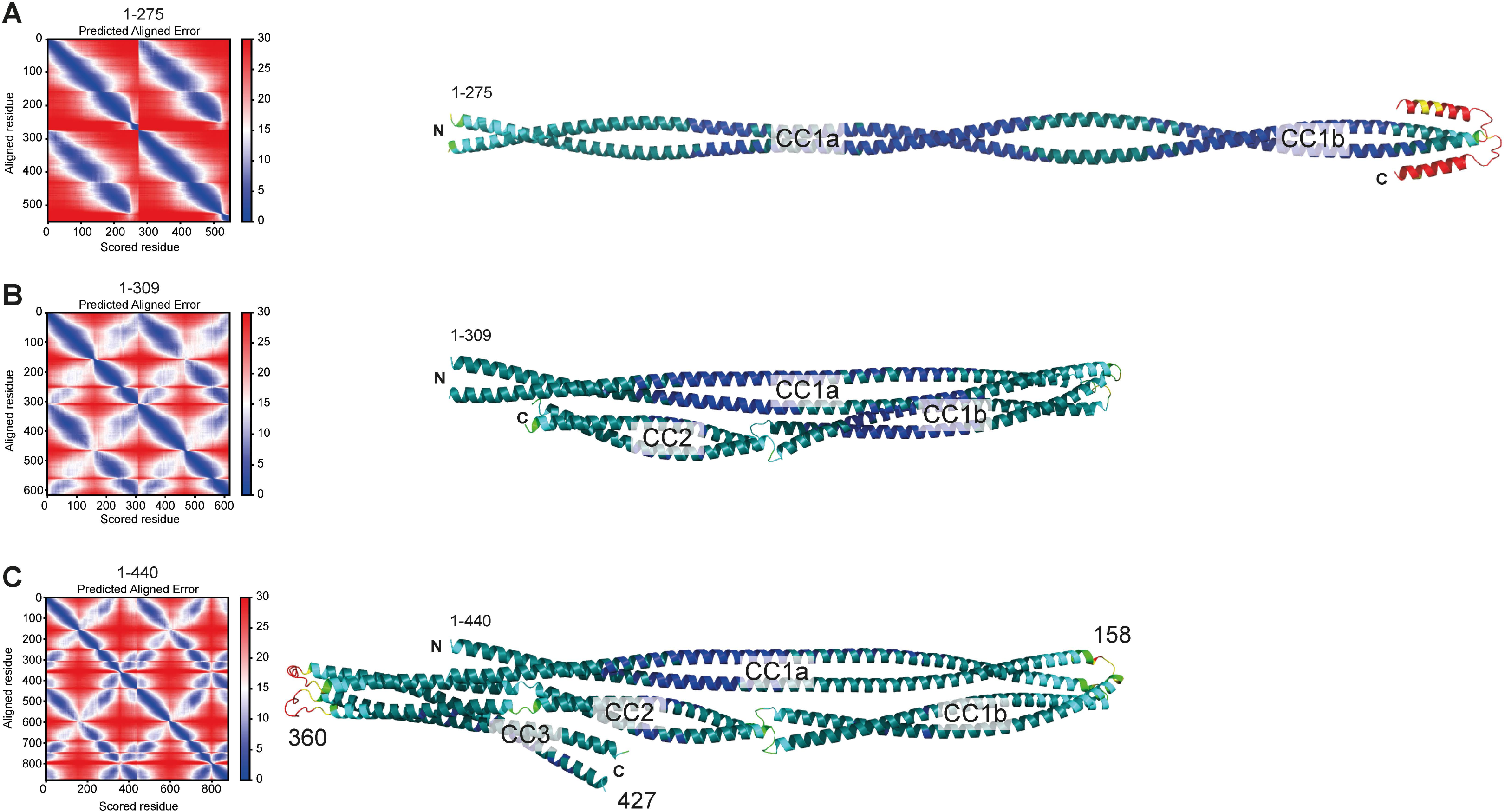
PAE plots and pLDDT scores. (**A**-**C**) Predicted Alignment Error (PAE) plots (left) and per-residue confidence score (pLDDT) of the three Spindly models discussed in the main text: (**A**) Spindly^1-275^. (**B**) Spindly^1-309^. (**C**) Spindly^1-440^. The pLDDT scores are displayed on the AF2 Multimer predictions of Spindly shown on the right (Blue: high confidence; Red: low confidence). The models are shown, with the same orientation, in Figure 1 and Figure 2. The PAE matrices refer to models of Spindly dimers, and correspondingly numbering of residues on the left and bottom of the plot is the number of residues in each chain multiplied by 2, and the second chain is plotted directly following the first. The parallel coiled-coils give rise to off-diagonal signals (blue) parallel to the main diagonal. Besides straight models like the one shown, a few predicted models of Spindly^1-275^ also showed a folded-back conformation. However, there are no additional off-diagonal signals for the Spindly^1-275^ construct in the PAE plots, suggesting that even if Spindly^1-275^ explores folded-back conformations, these are not stable. Off-diagonal signals perpendicular to the main diagonal are instead clearly visible in the Spindly^1-309^ and Spindly^1-440^ constructs, consistent with a folded back conformation. A predicted folded conformation of Spindly^1-440^ was also observed with orthologous sequences (unpublished results).

**Figure 2 – Supplement 1.**
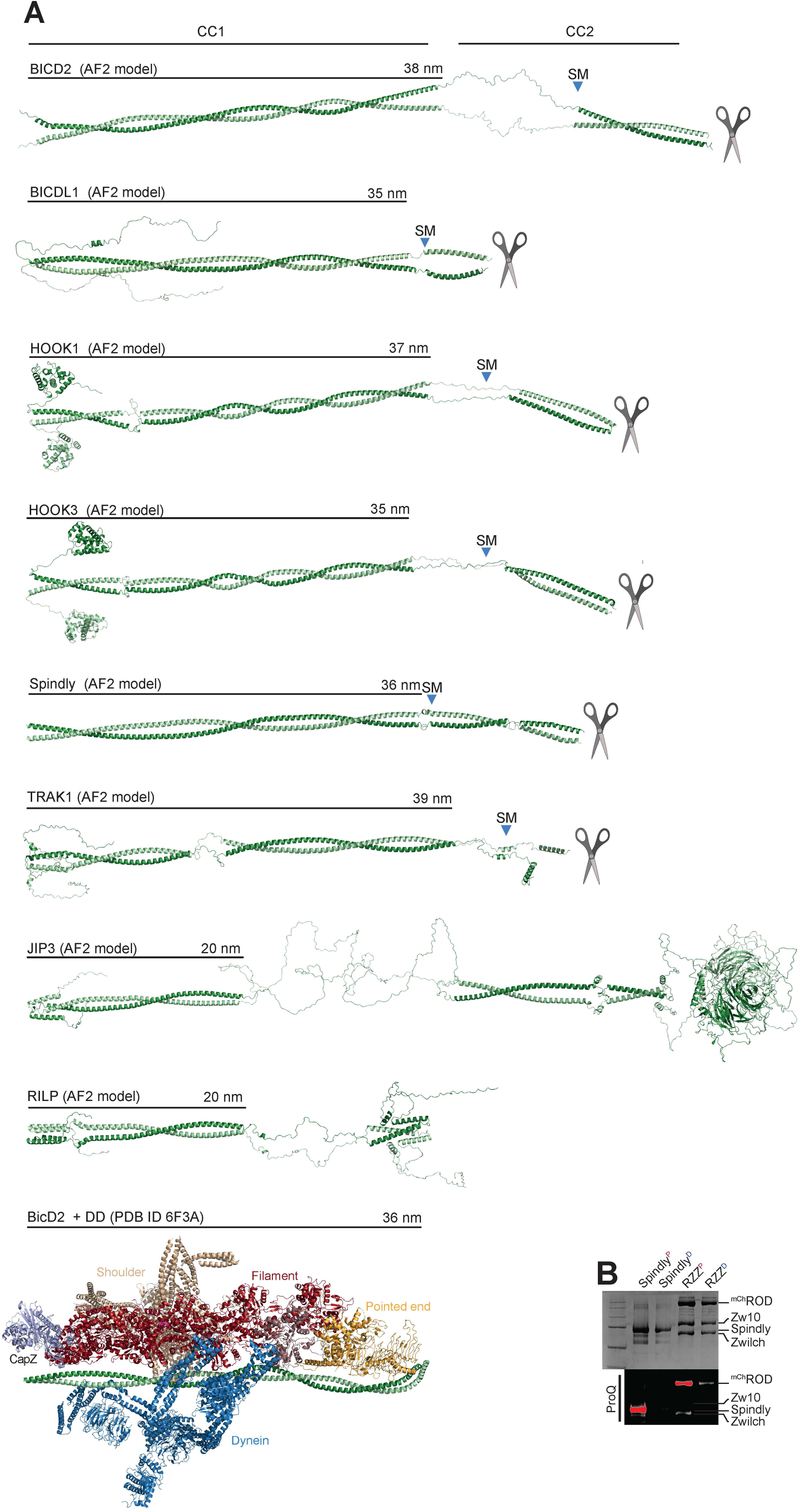
Gallery of AF2 predictions of the structure of representative adaptors. (A) AF2 ColabFold models of the indicated adaptors. The scissor symbol indicates that the displayed cartoon models were truncated after the CC2 region. SM is the Spindly motif. The length of CC1 coiled-coil is indicated. As indicated in the Methods section, AF2 and variants can predict different quaternary structures for the adaptors, including trimeric or tetrameric coiled coil formation, if three or four chains respectively are used as input. However, with trimers or tetramers, the PAE values estimated between the same positions on different protomers are high to very high (i.e. insignificant), indicating that the predictions are less likely to be accurate (unpublished results). We only present predictions of dimers here, as there is experimental evidence for several of them that their active conformation is the dimer (Isabet et al., 2009; Kelkar et al., 2000; Lee et al., 2018; Urnavicius et al., 2018; Urnavicius et al., 2015; Wu et al., 2005). For clarity, models were straightened as discussed in Methods. Even if Spindly is predicted to adopt a closed conformation when CC2 is present (see main text), here for comparison we show the extended open conformation expected to bind DD. The model of the DD complex with BicD2 (PDB ID 6F3A) (Urnavicius et al., 2018) shows that the length of the experimentally modelled BicD2 coiled-coil is approximately identical to the length of CC1 predicted by AF2 for many of the displayed adaptors. (B) Coomassie-stained SDS-PAGE and ProQ diamond staining of phosphorylated and dephosphorylated RZZ and Spindly. Samples were initially dephosphorylated with λ-PPase (samples indicated as ‘D’) and later re-phosphorylated with the mitotic kinases CDK1/CyclinB, MPS1, and Aurora B (samples indicated as ‘P’).

**Figure 3 – Supplement 1.**
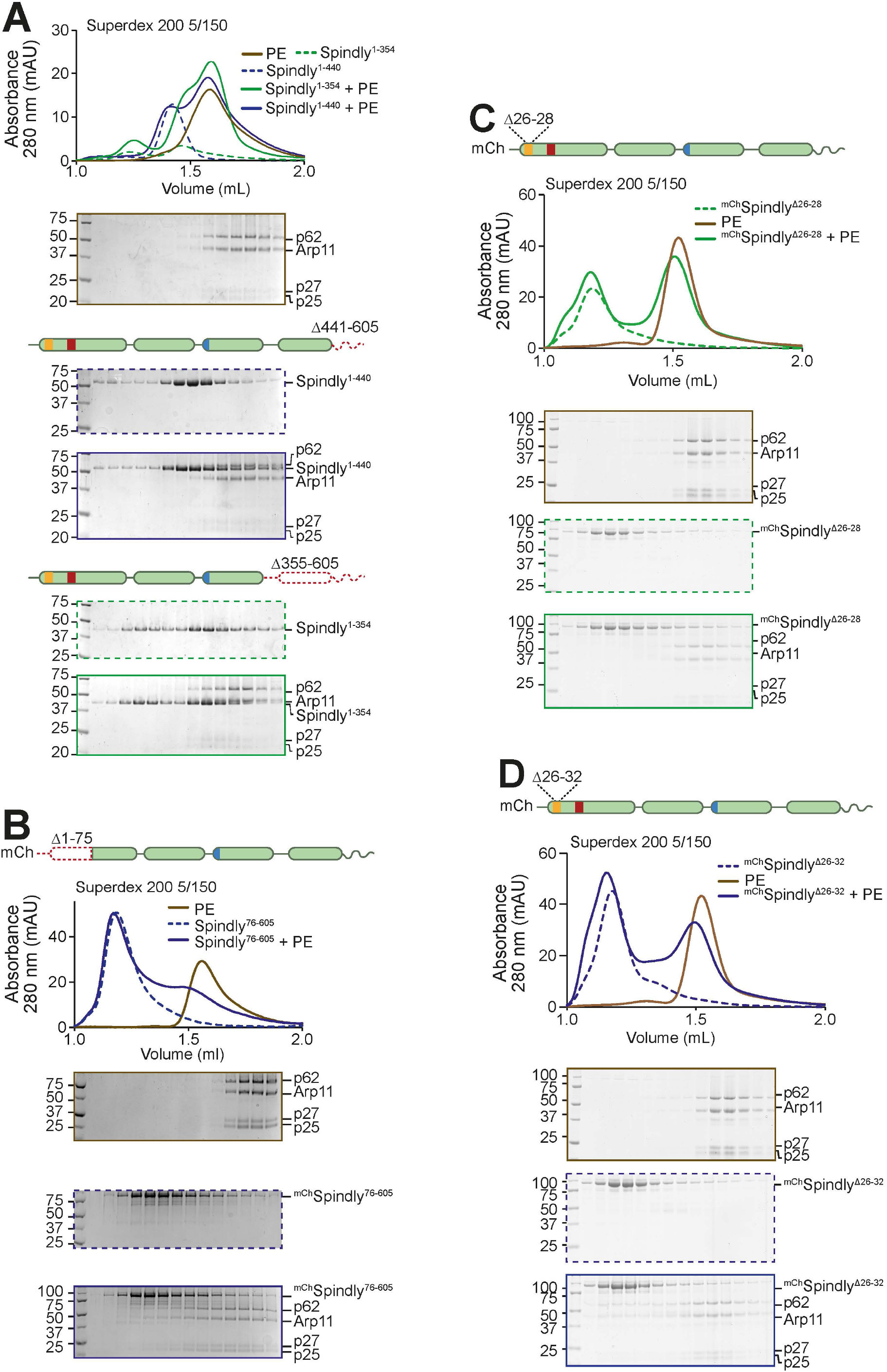
Additional analyses of the Spindly N-terminal autoinhibitory region. (**A**-**D**) Additional analytical SEC interaction assays between the Dynactin pointed end (brown) and the indicated Spindly constructs. The complex run is always represented with a continuous line, the Spindly construct with a dashed line. (**A**) Spindly^1-440^ (blue); Spindly^1-354^ (green); (**B**) ^mCh^Spindly^76-605^ (blue); (**C**) ^mCh^Spindly^Δ26-28^ (green); (**D**) ^mCh^Spindly^Δ26-32^ (blue). The PE alone control in panel A is shared with Figure 3D, between panels C and D, and between panel B and panels Figure 5 – Supplement 1A-B.

**Figure 5 – Supplement 1.**
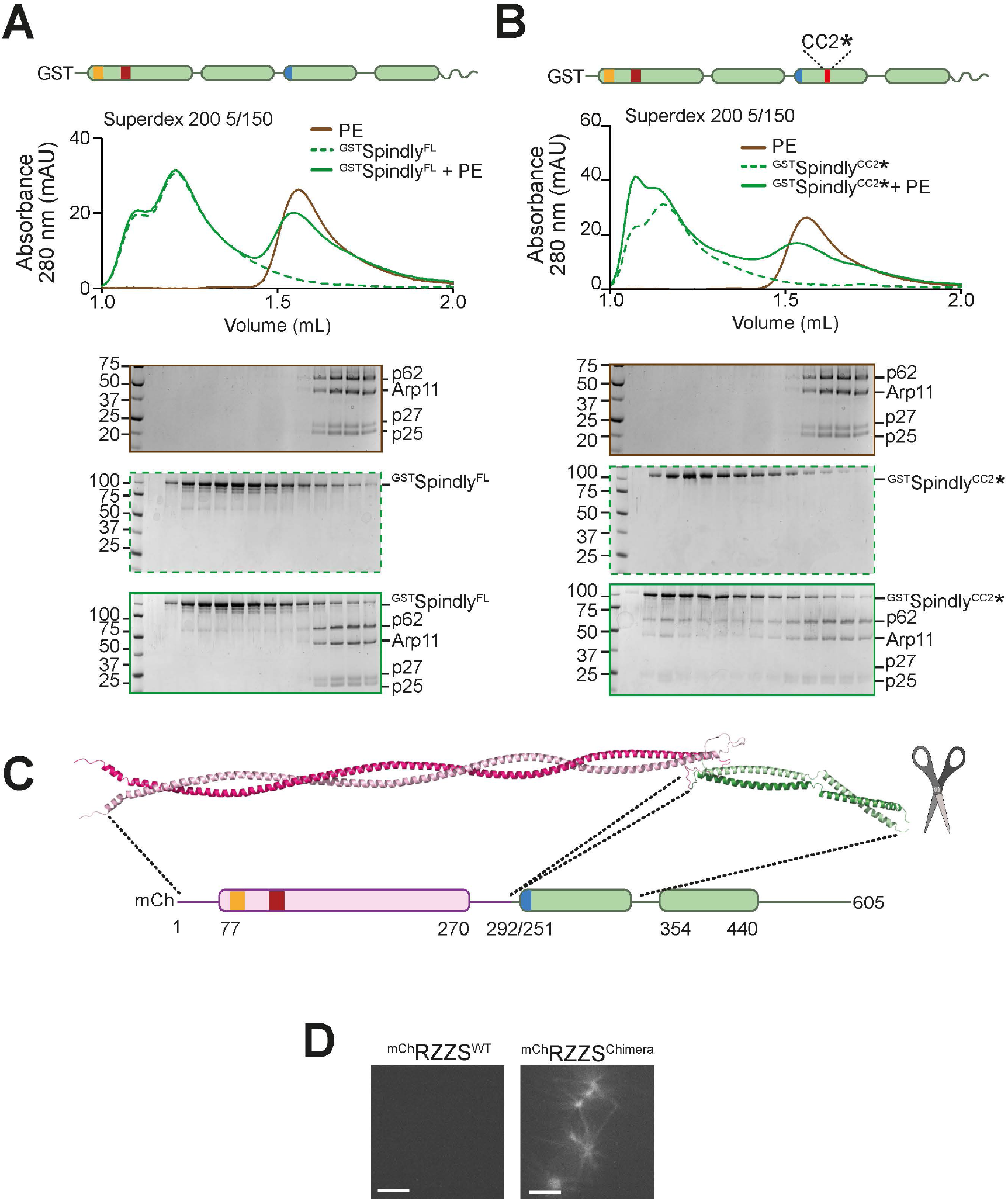
Additional characterization of Spindly binding to PE. (**A**) Size-exclusion chromatography (SEC) separation of ^GST^Spindly (dotted green line), the pointed end (PE) complex (brown line), and their mixture (continuous green line). (**B**) SEC separation of ^GST^Spindly^CC2*^ (dotted green line), the pointed end (PE) complex (brown line), and their mixture (continuous green line). The PE control is shared between the two shown experiments and with Figure 3 – Supplement 1B. (**C**) The AF2 ColabFold model of the BicD2-Spindly chimera shows CC1 is continuous. (**D**) Spinning-disk confocal fluorescence microscopy-based filamentation assay at 561 nm with the indicated ^mCh^RZZS species (4 µM RZZ, 8 µM Spindly) at 20°C in absence of MPS1 kinase. Scale bar: 5 µm.

**Figure 6 – Supplement 1.**
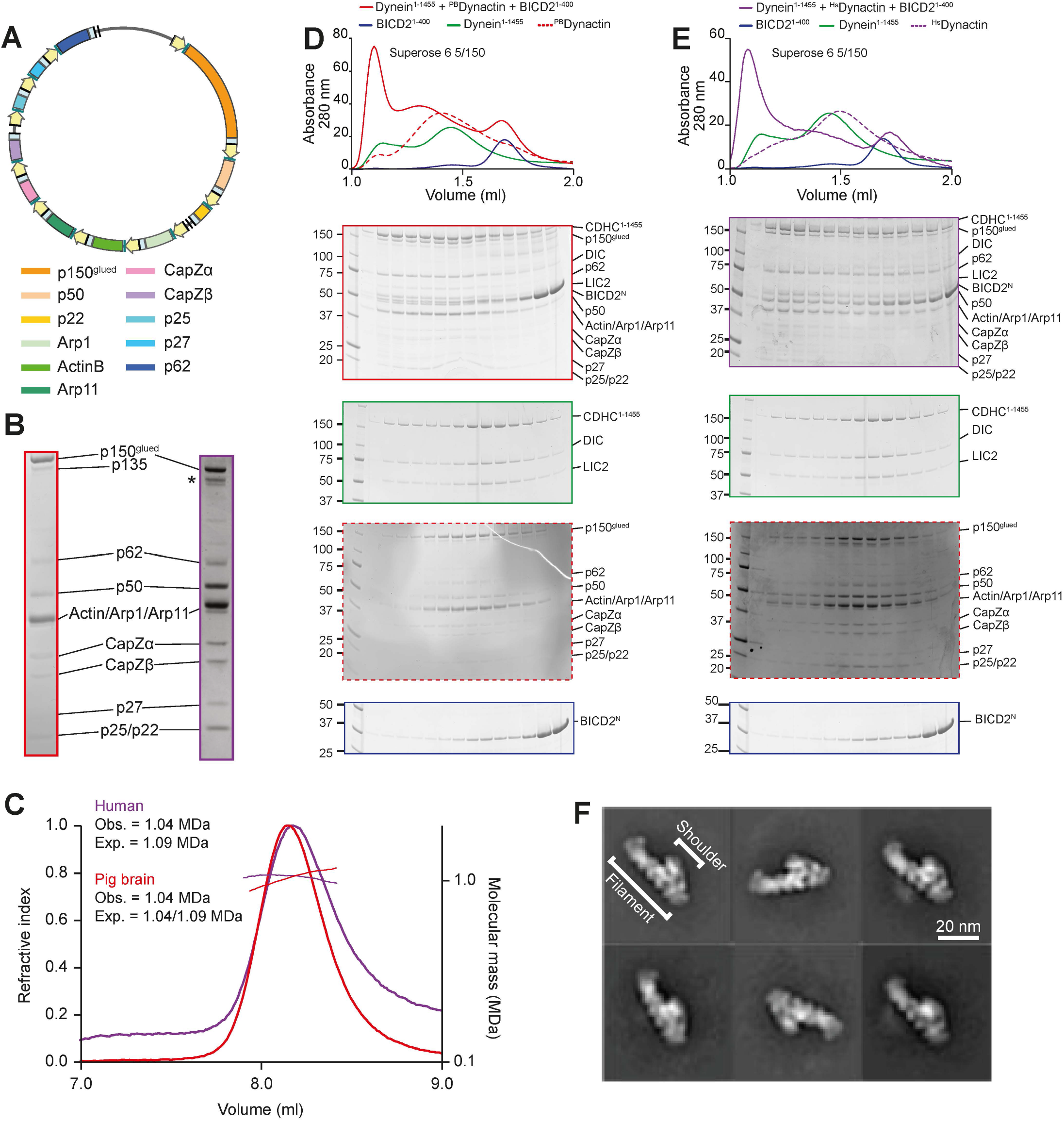
Recombinant human Dynactin. (**A**) Map of the Dynactin expression plasmid. Individual subunits are labelled according to list below. CMV promoters and enhancers are labelled in yellow. PolyA signals are labelled in light blue. (**B**) Comparison of SDS-PAGE of pig brain Dynactin and recombinant human Dynactin, in Coomassie staining. (**C**) SEC-MALS analysis of human (purple) and pig brain (red). Pig brain Dynactin contains a mix of the p150 isoforms p150^glued^ and p135, yielding two different expected masses. (**D**-**E**) SEC assays on a Superose 6 5/150 column to assess binding of Dynein tail, Dynactin, and BicD2^1-400^ with pig brain (**D**) and human (**E**) Dynactin. The Dynein tail and BicD2^1-400^ controls are shared between the two experiments. (**F**) 2D class averages from negative stain imaging of recombinant human Dynactin.

**Figure 7 – Supplement 1.**
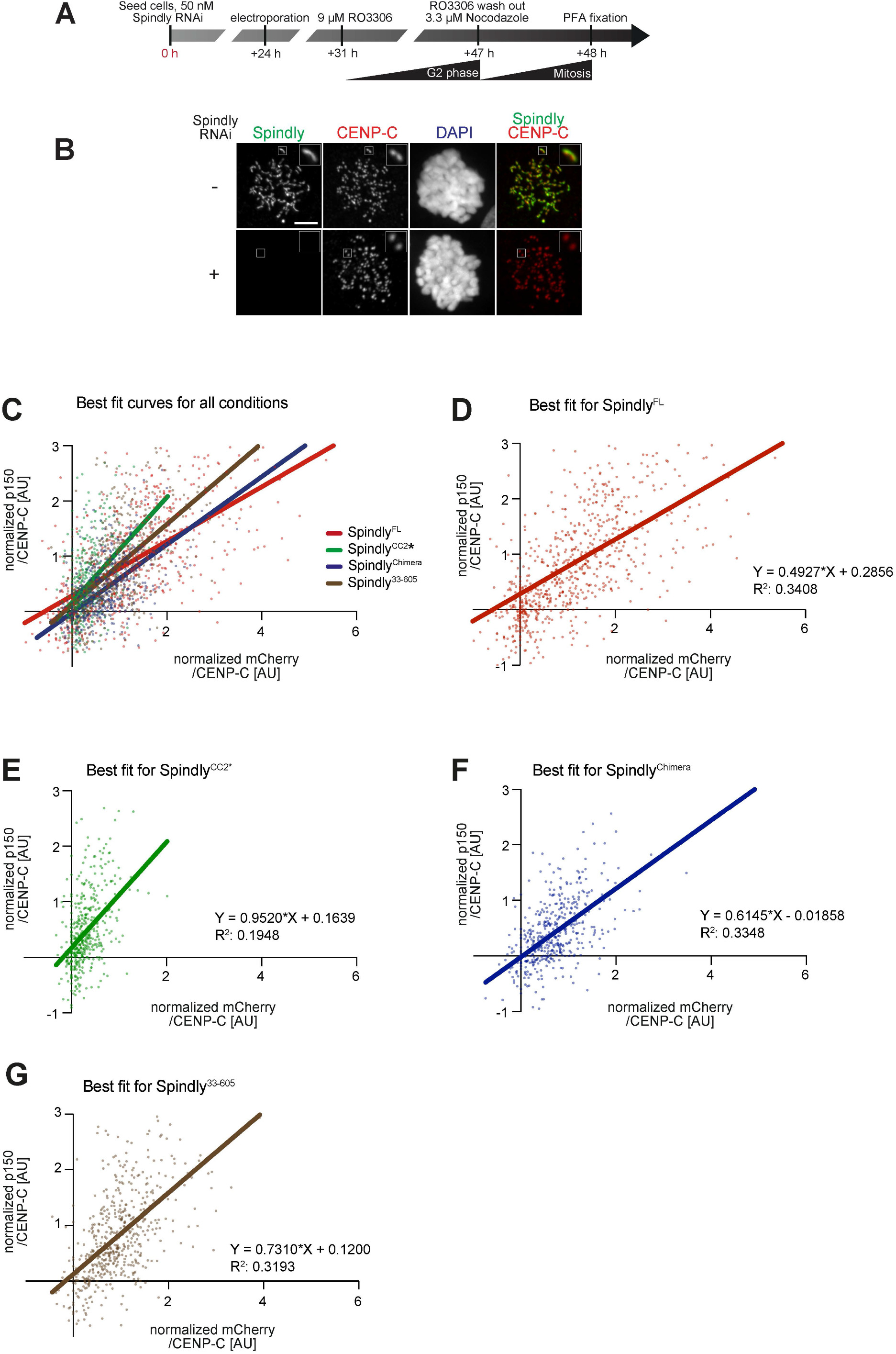
Additional data on kinetochore levels of Dynactin with Spindly mutants. (**A**) Schematic of the RNAi and complementation by electroporation with recombinant proteins. (**B**) Representative images of RNAi control cells and cells depleted of Spindly by RNAi. **C**-**G**) Least square fitting through the distribution of data points reporting for each kinetochore the CENP-C-normalized ^mCh^Spindly intensity on the X-axis and the CENP-C normalized p150^glued^ intensity on the Y-axis. (**C**) Fit curves with data points. (**D**) Individual best fit for ^mCh^Spindly^FL^. (**E**) Individual best fit for ^mCh^Spindly^CC2*^. (**F**) Individual best fit for ^mCh^Spindly^Chimera^. (**G**) Individual best fit for ^mCh^Spindly^33-605^.

**Table S1.**
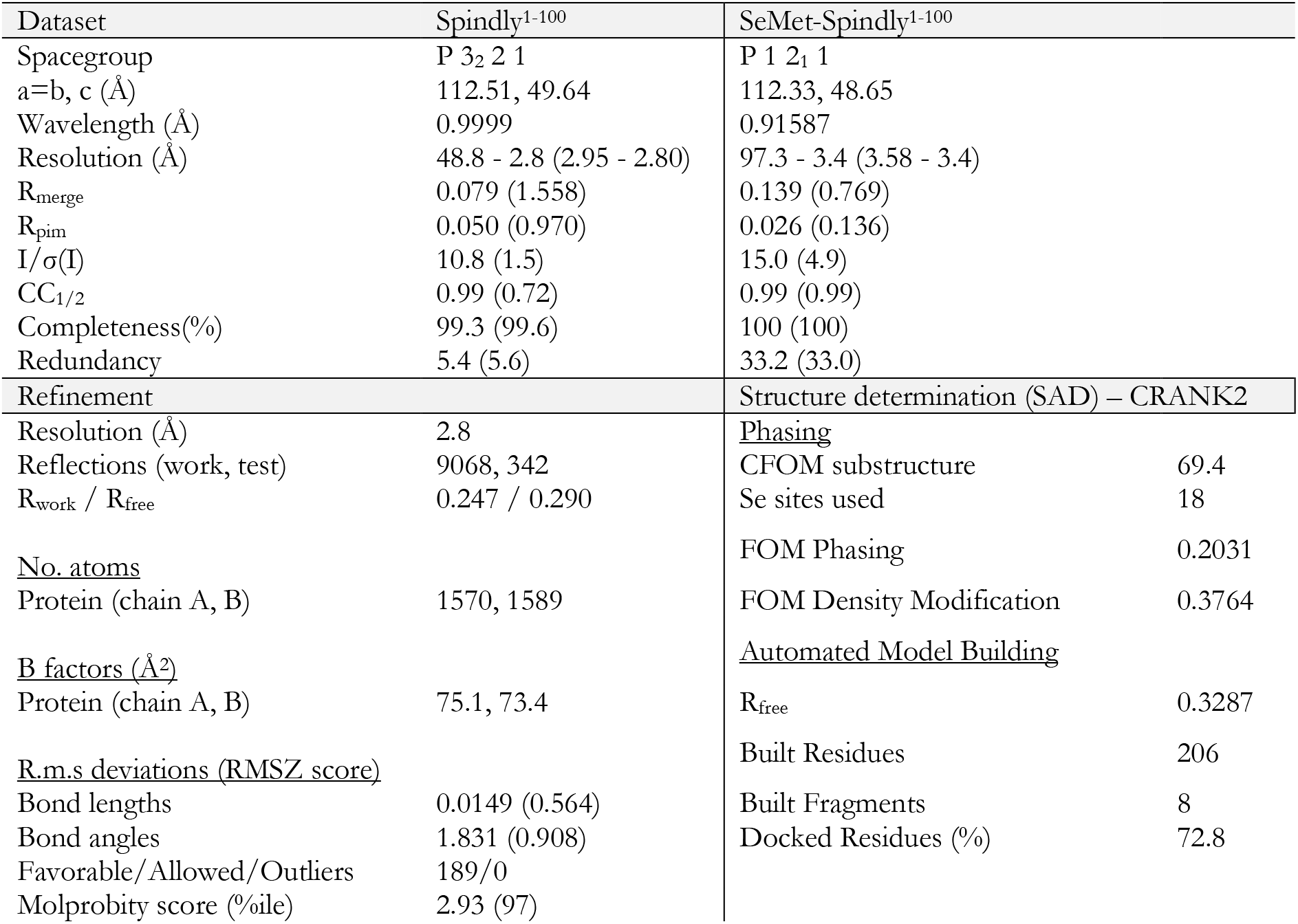
Crystallographic data.

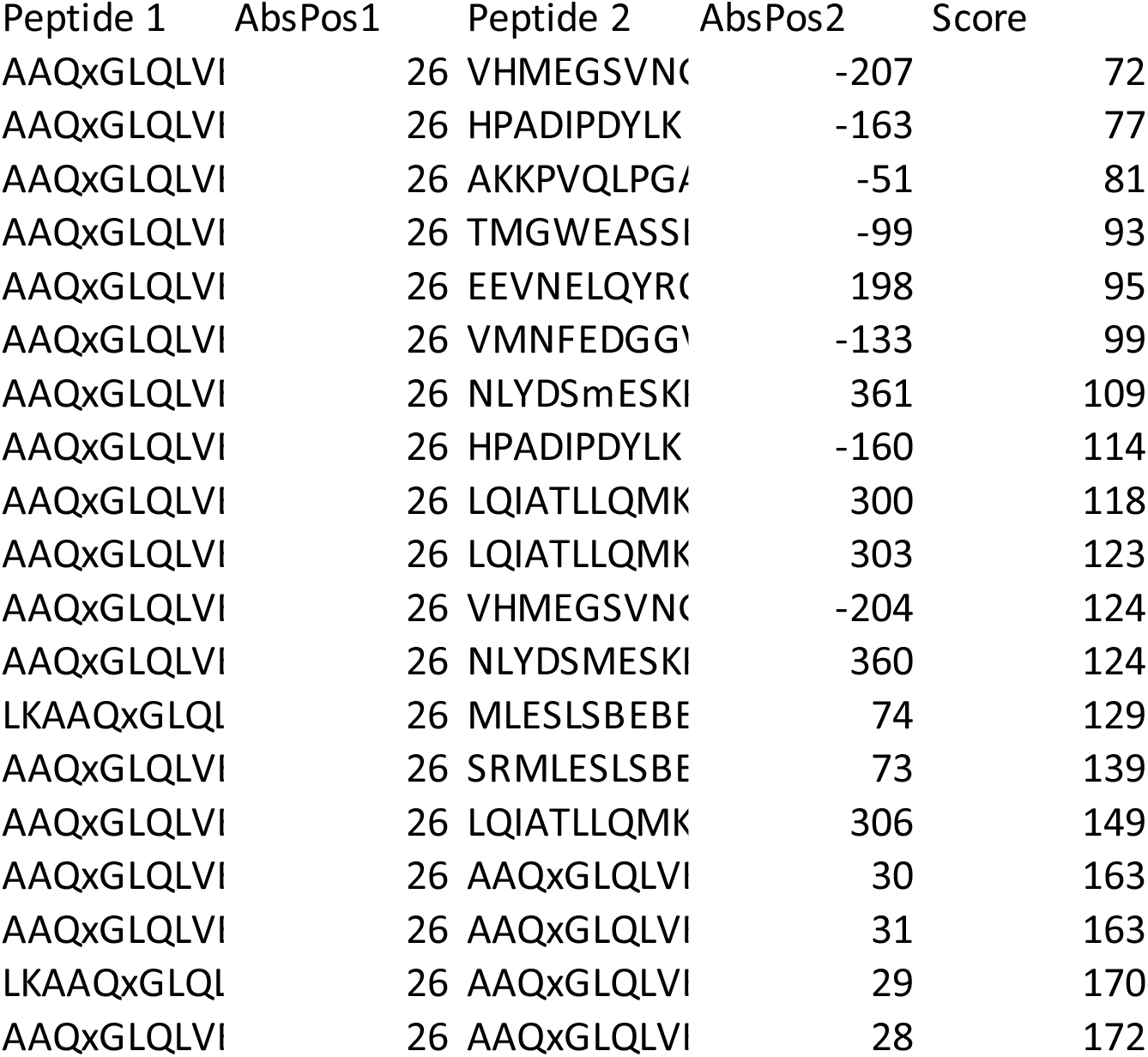

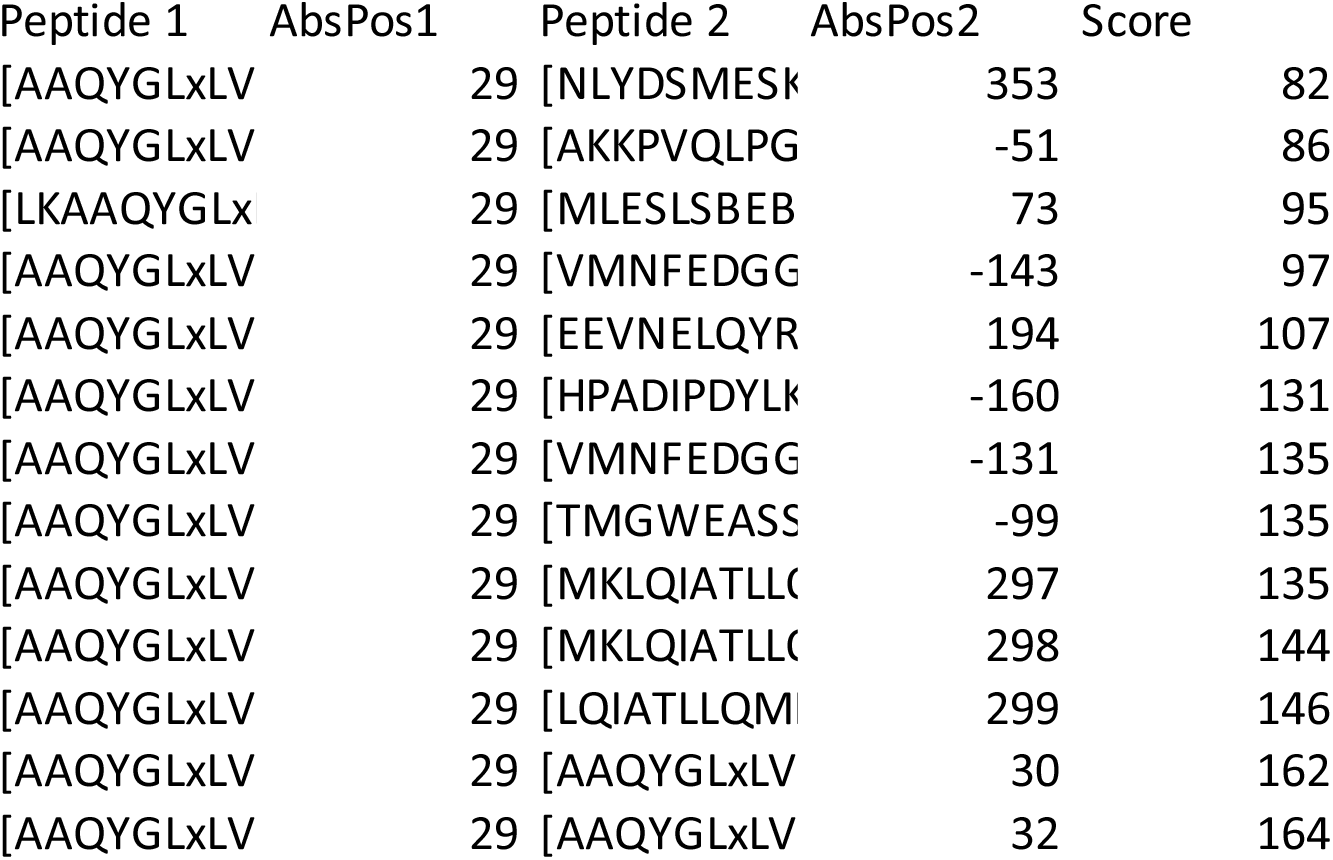

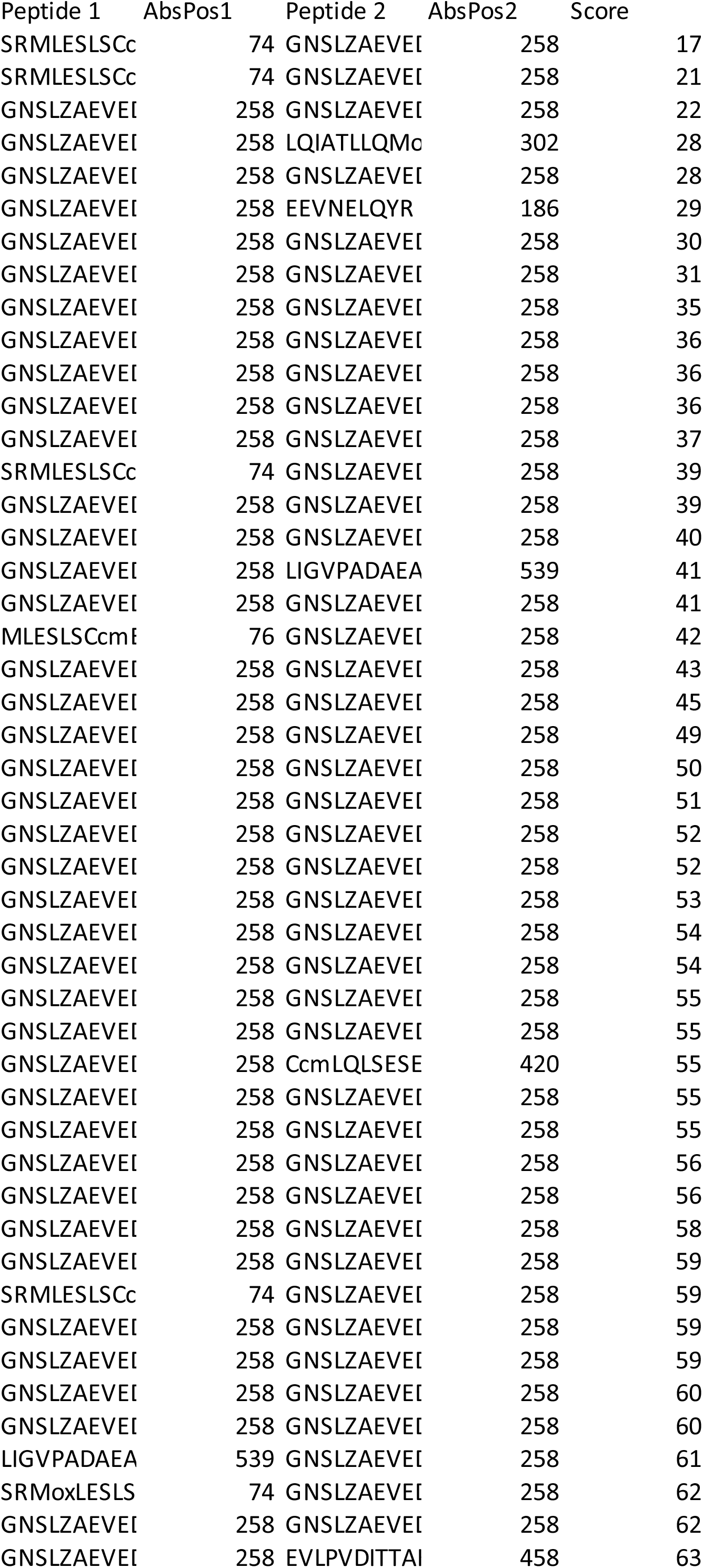
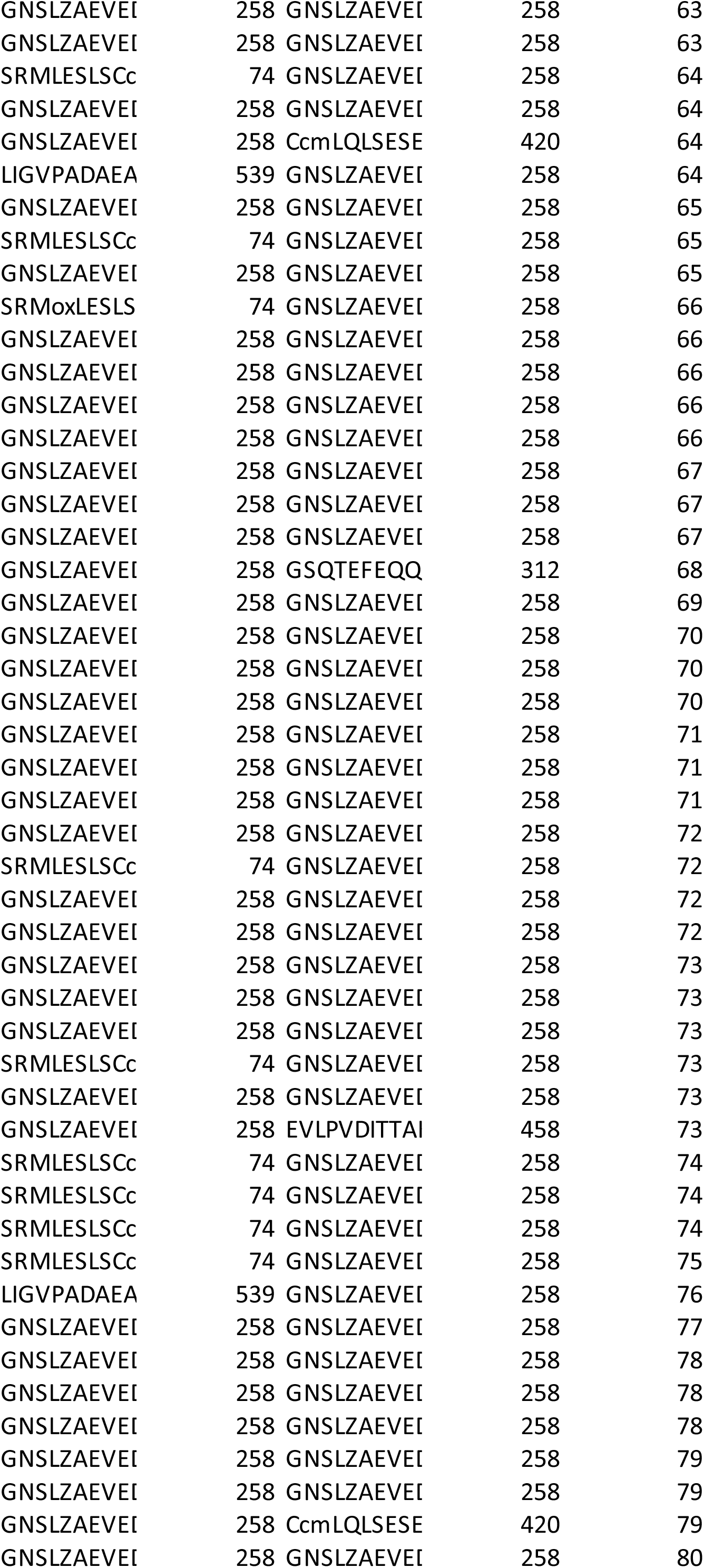
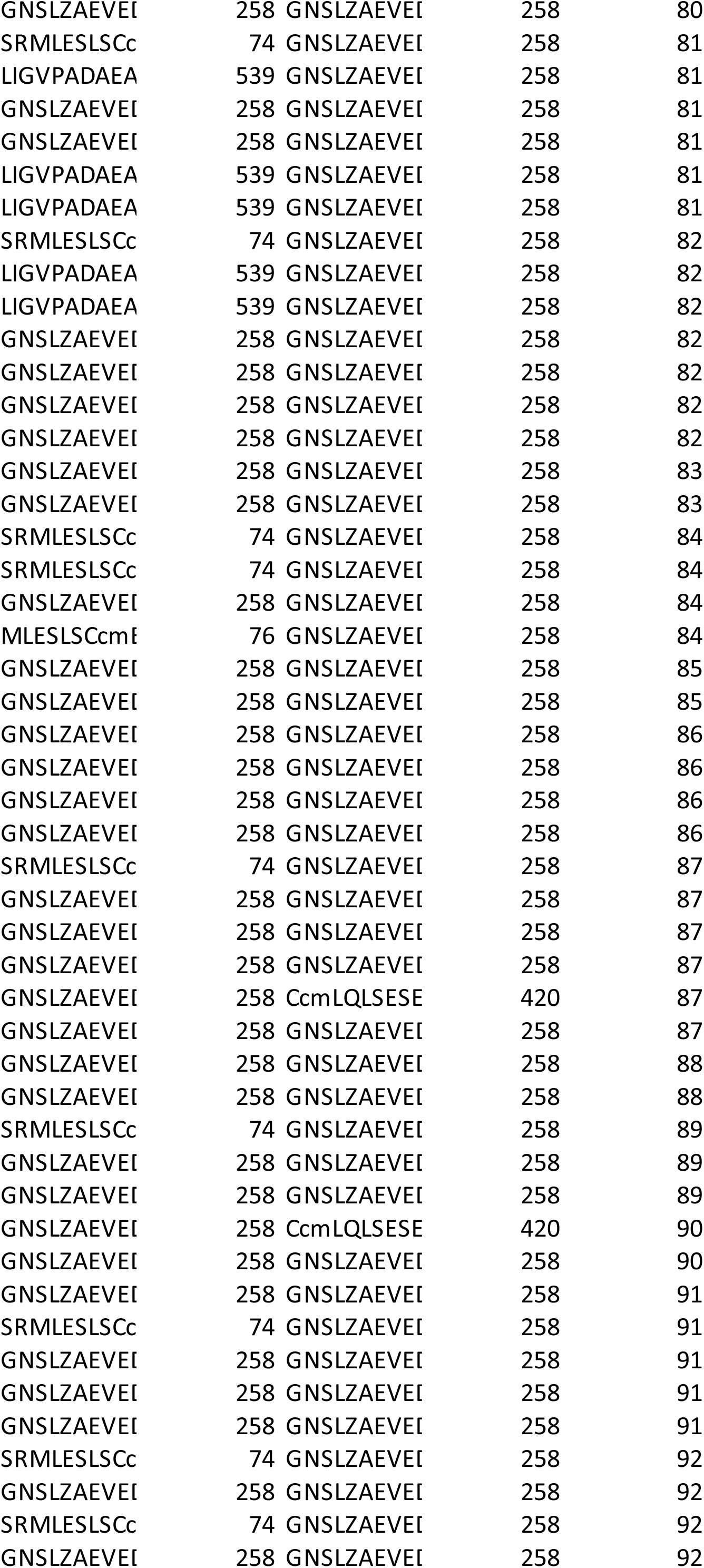
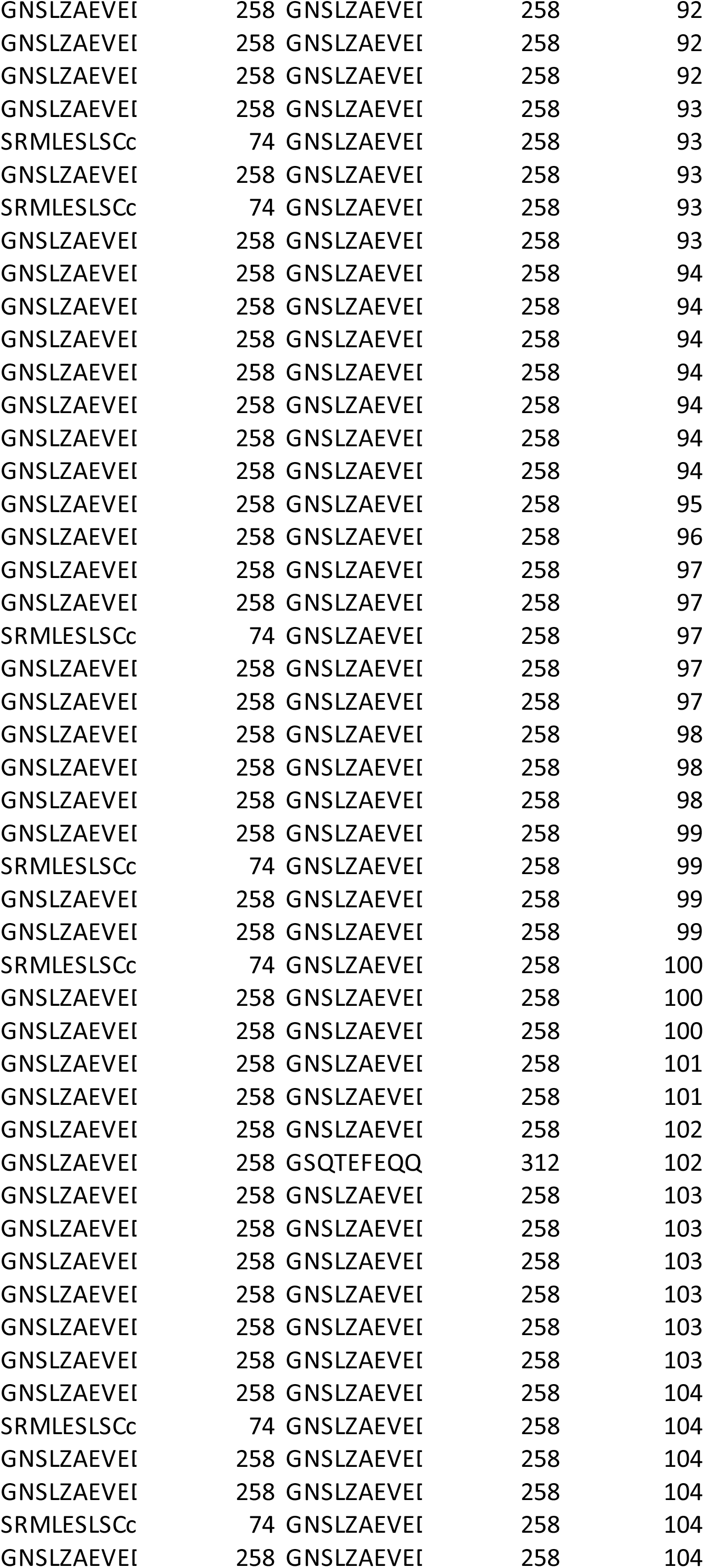
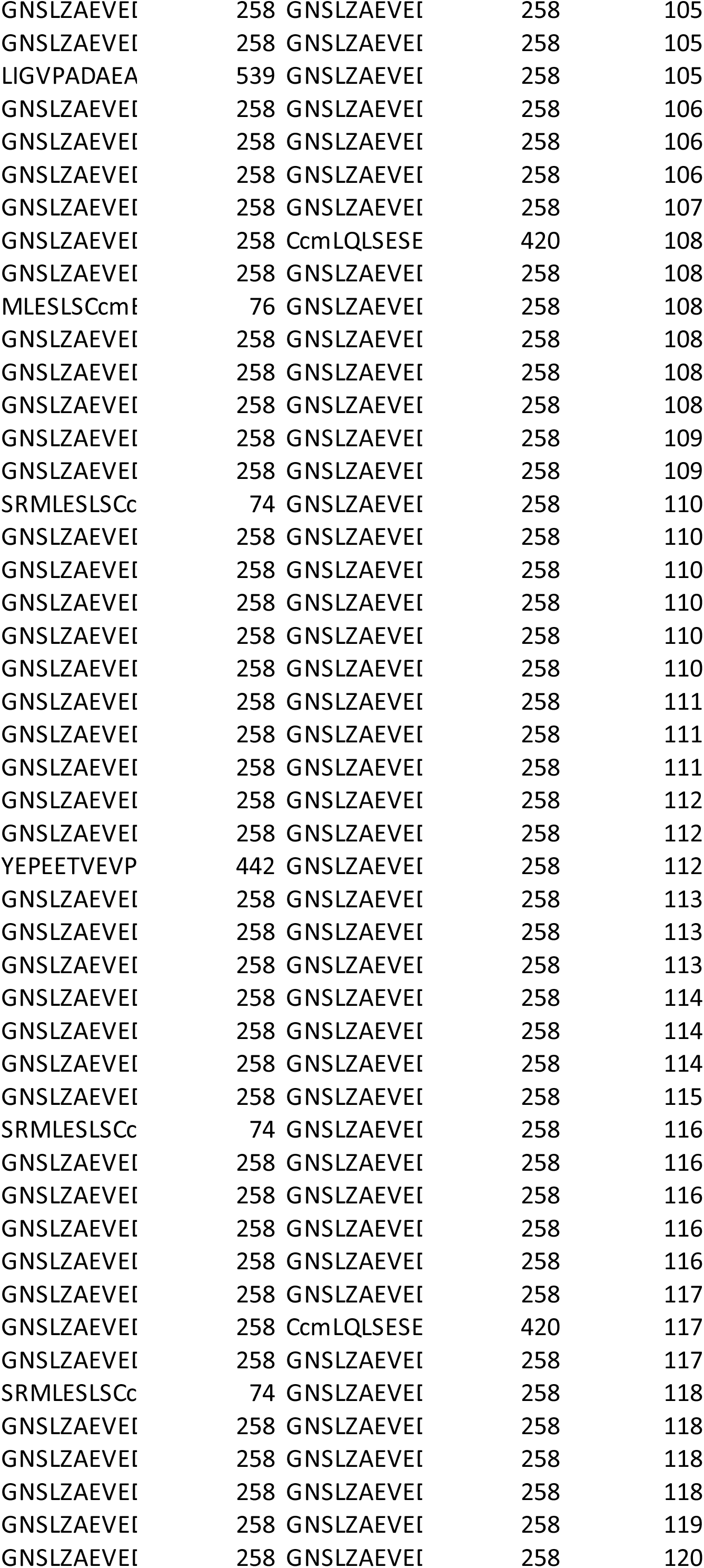
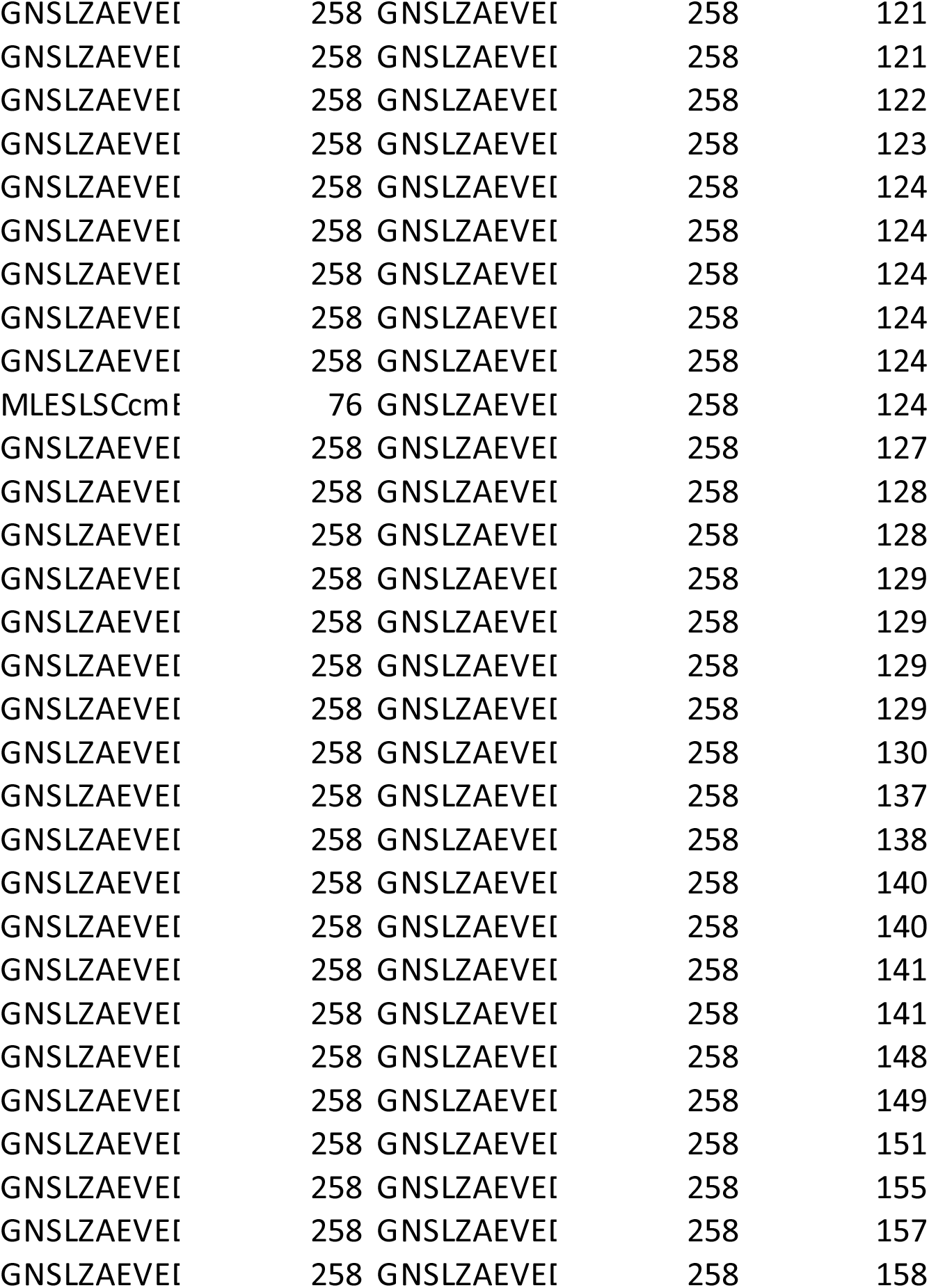

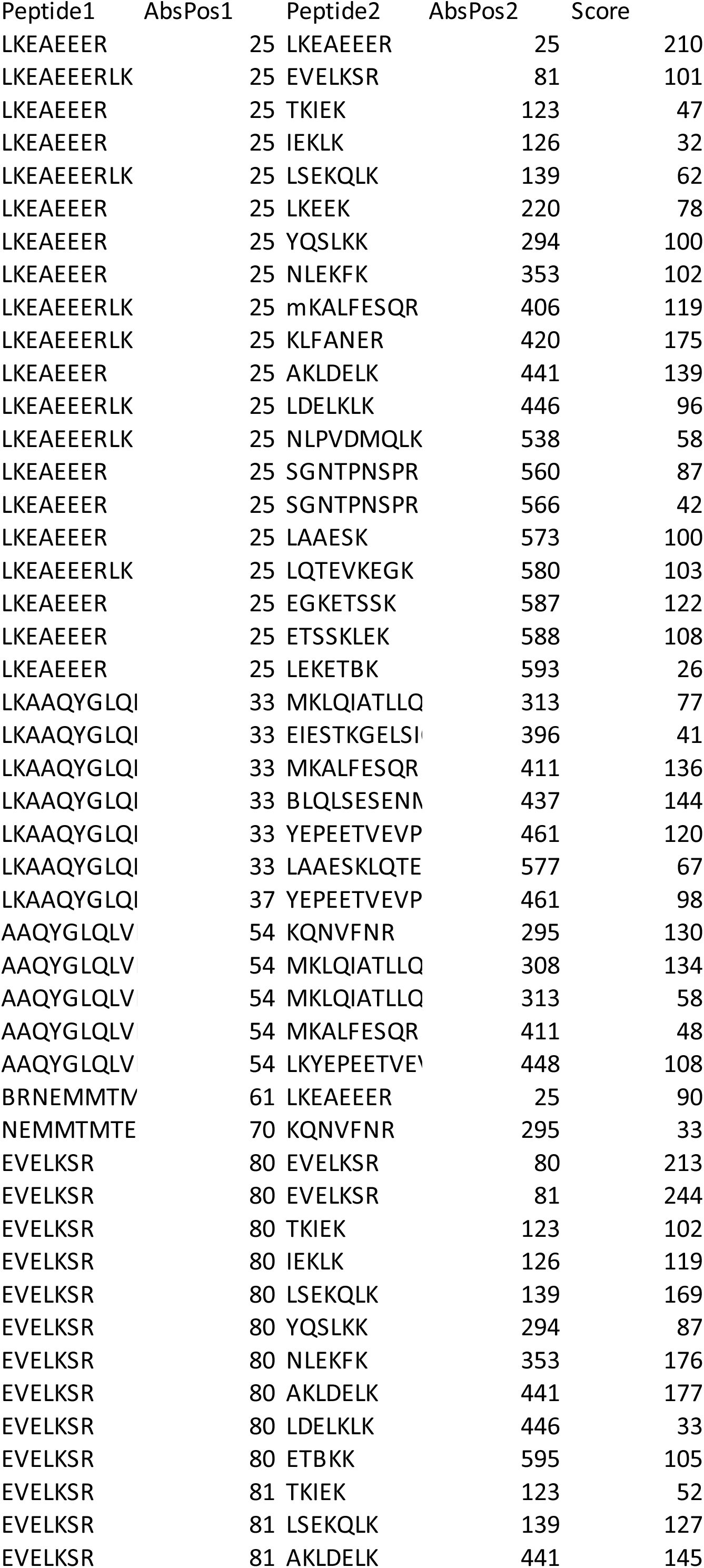
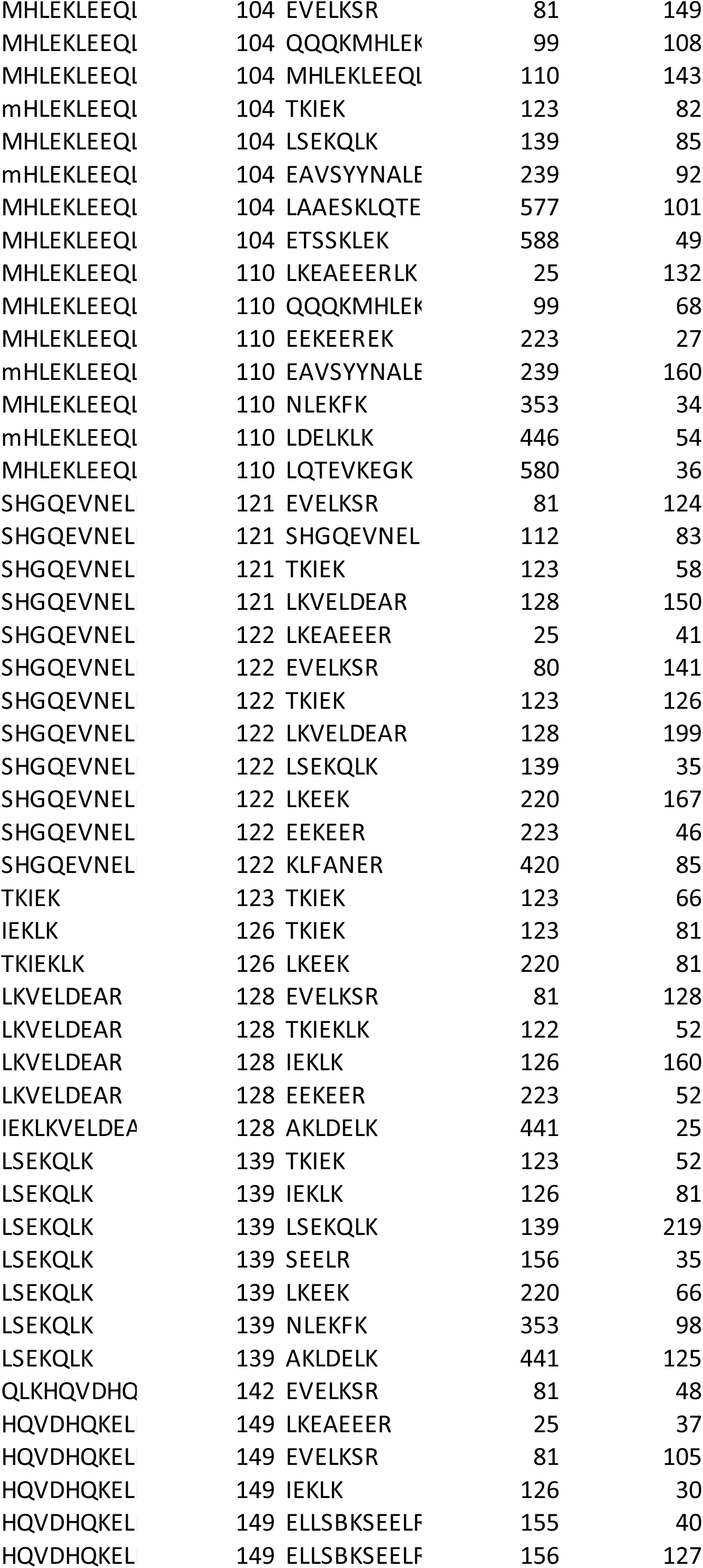
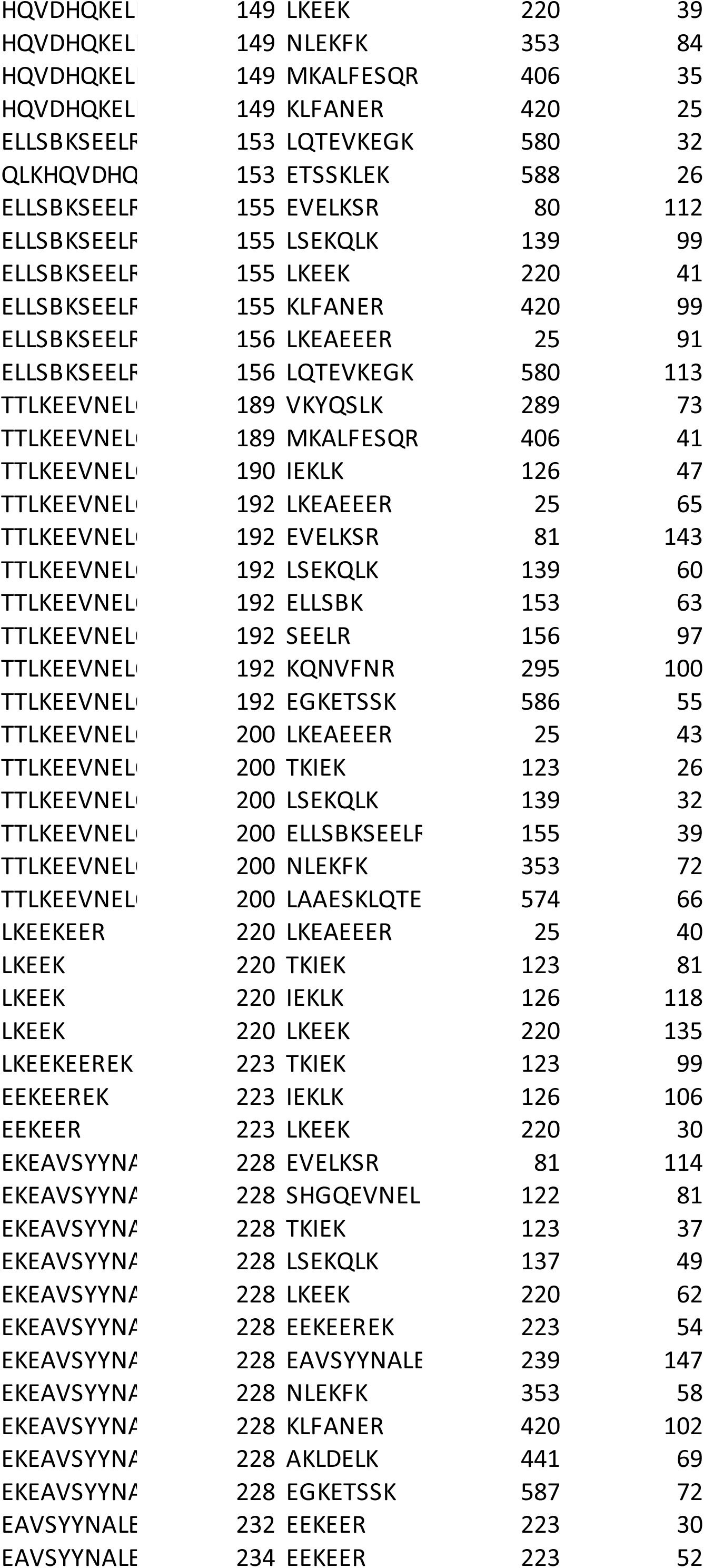
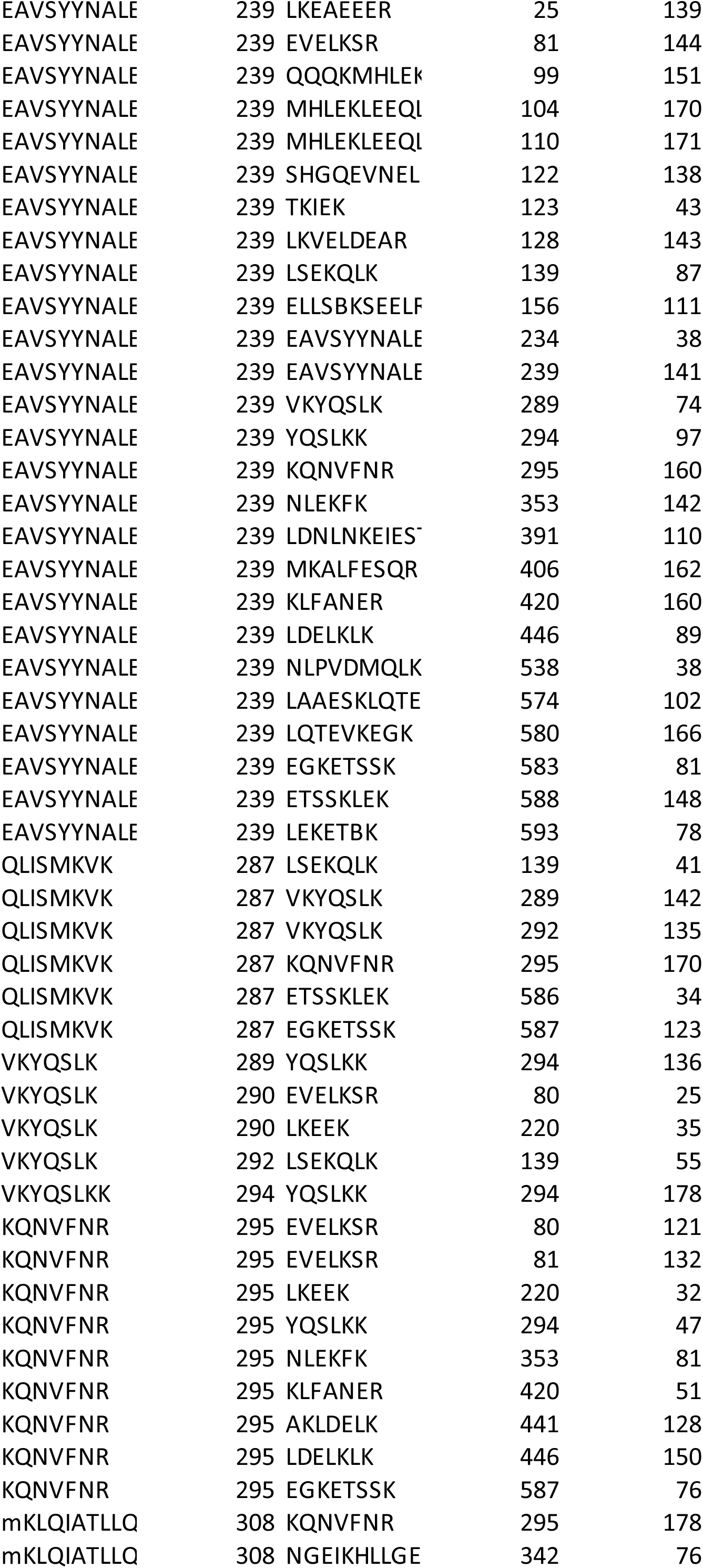
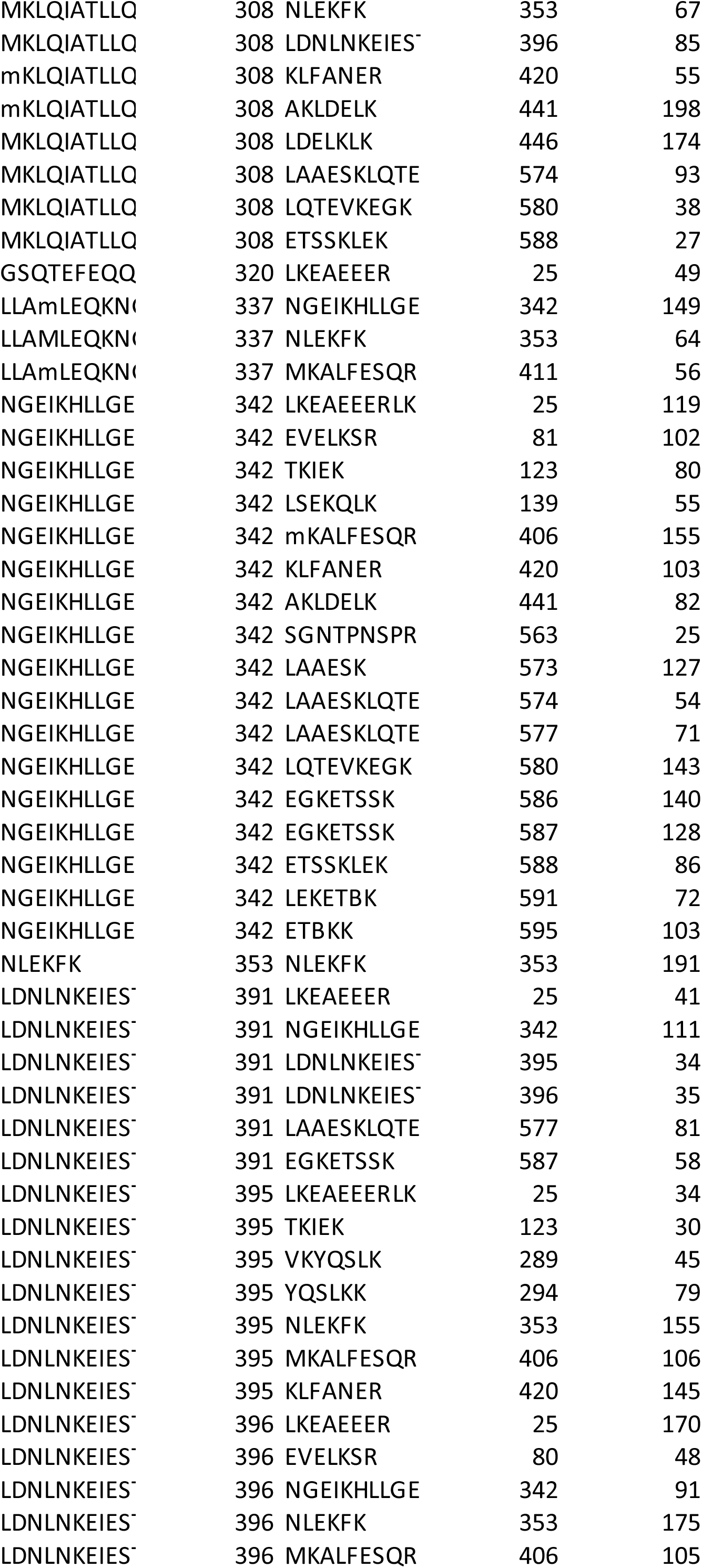
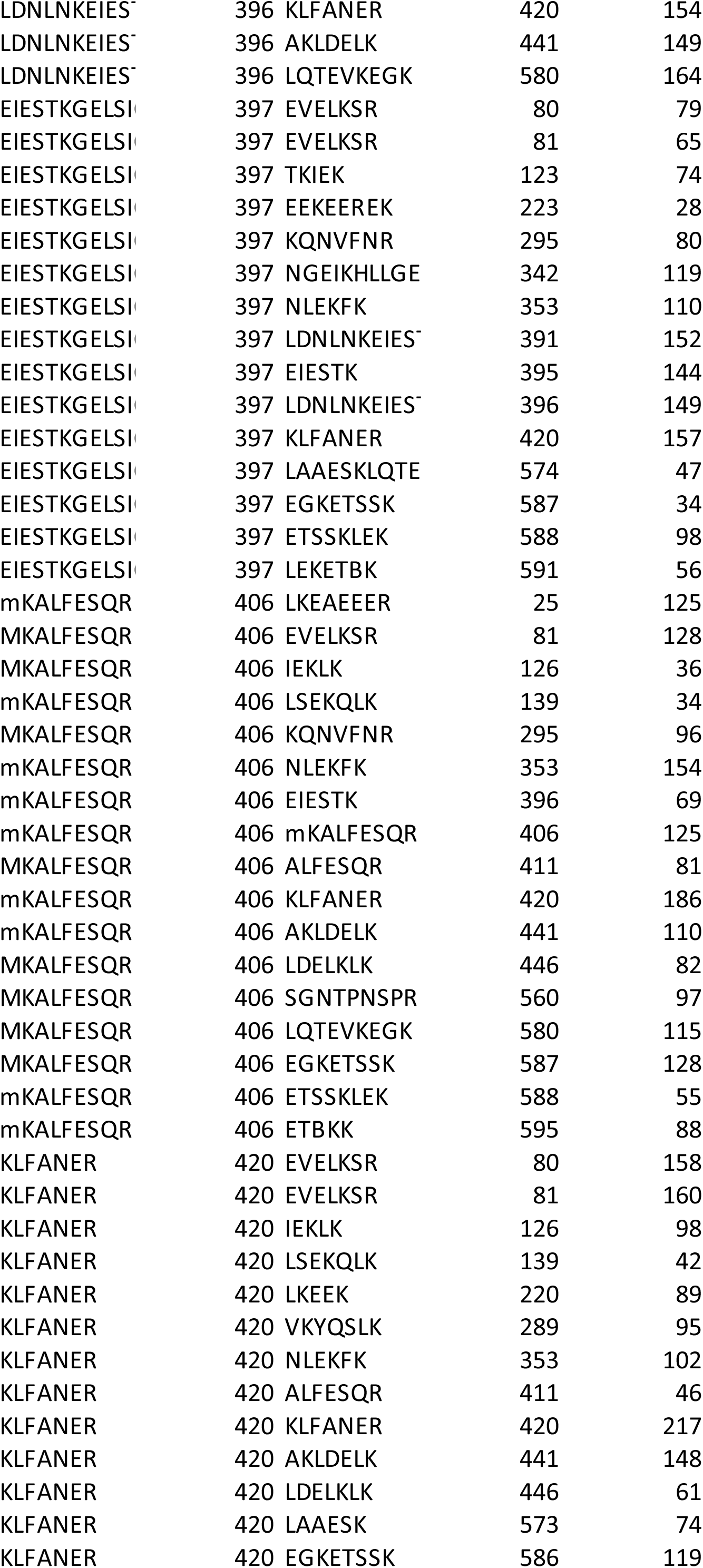
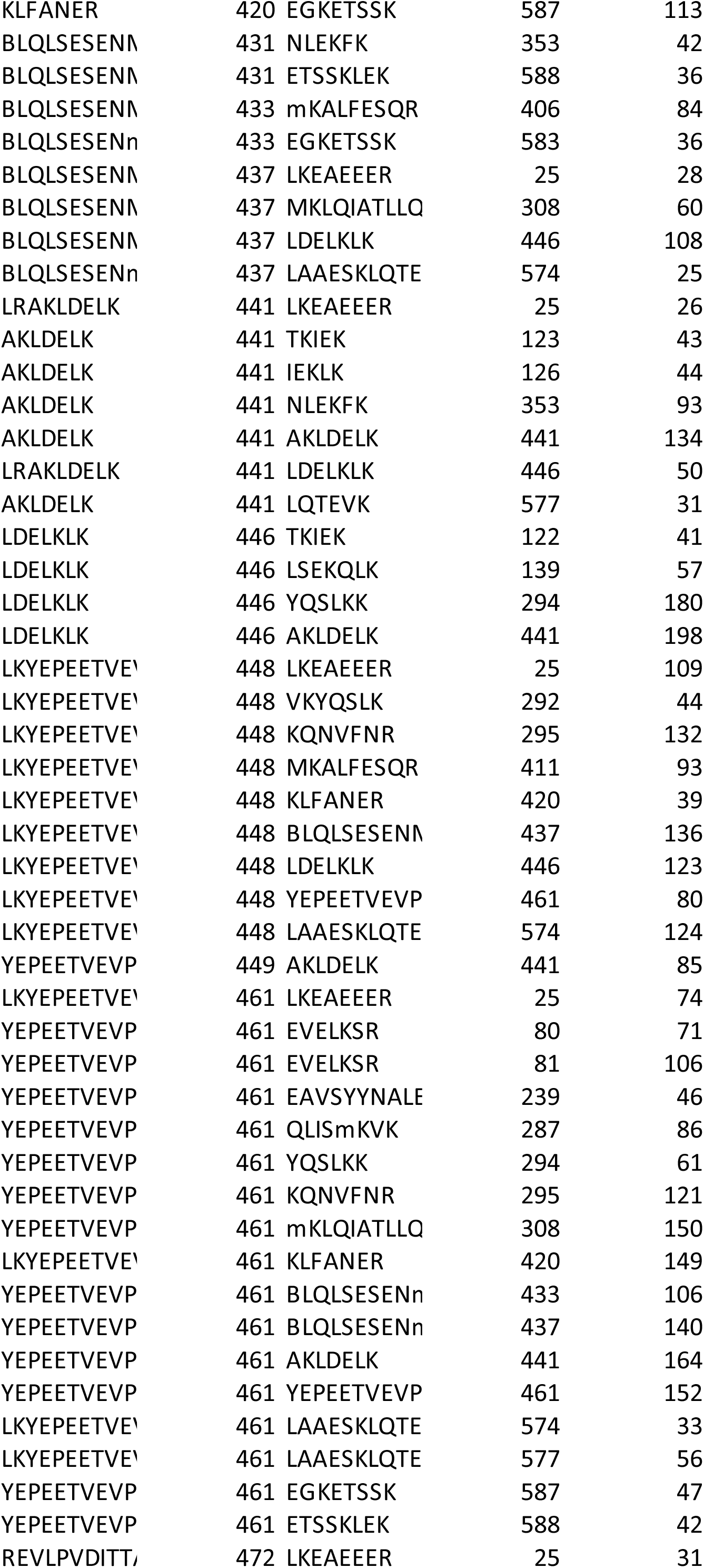
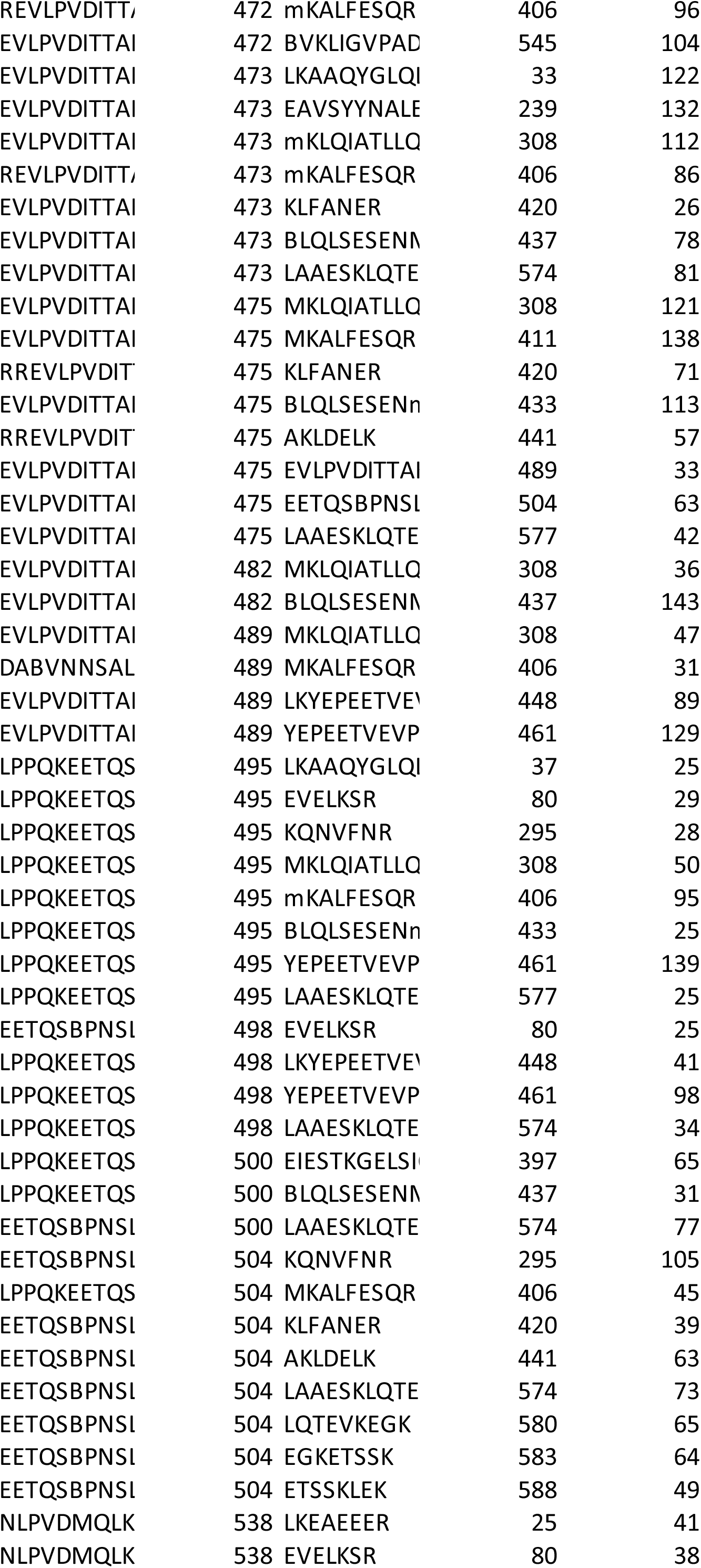
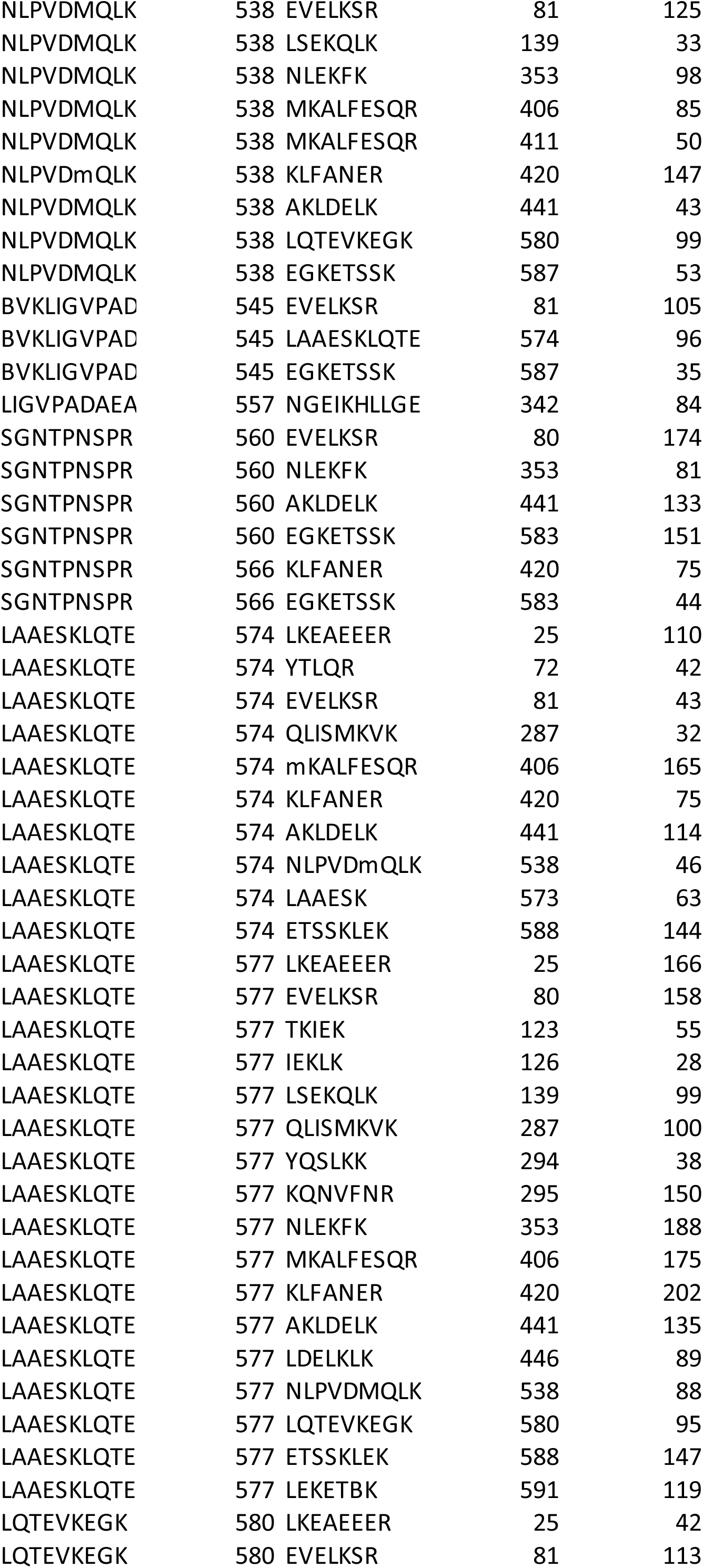
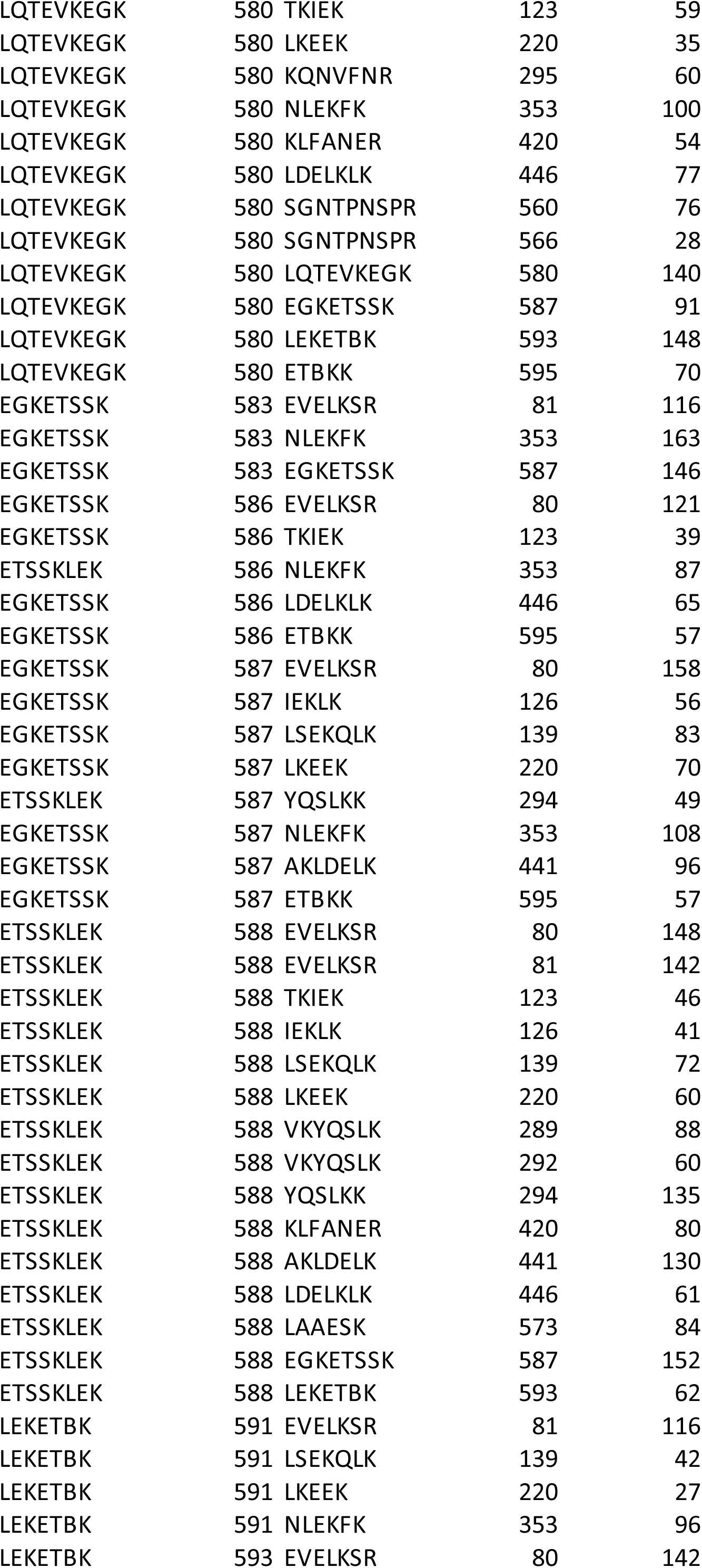
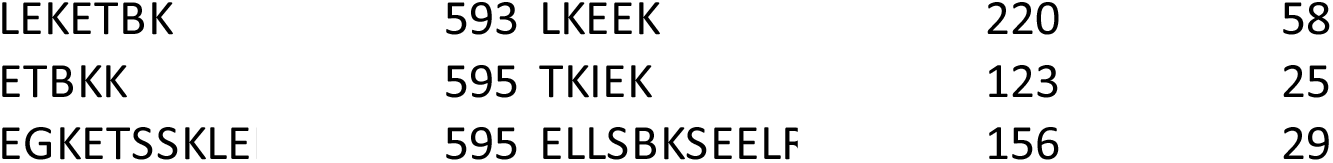

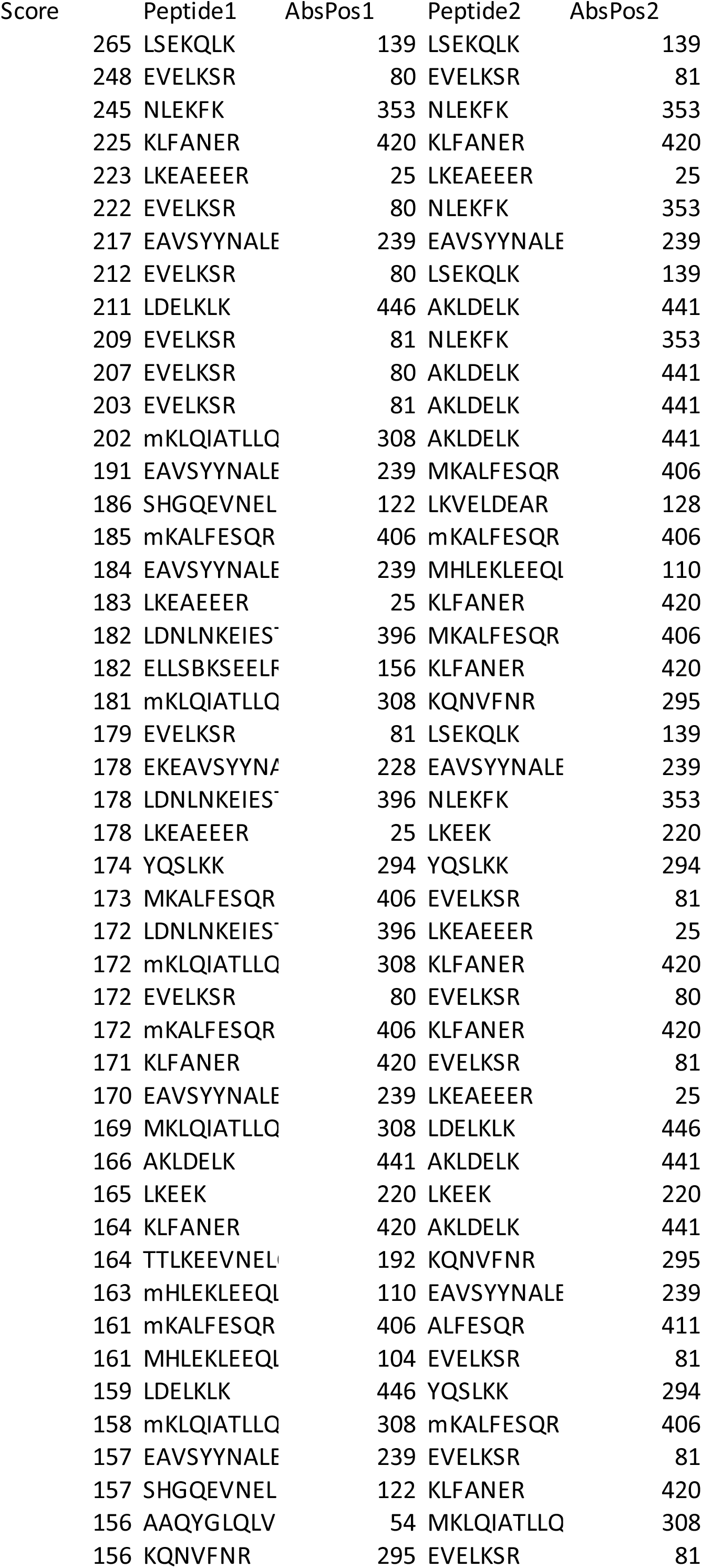
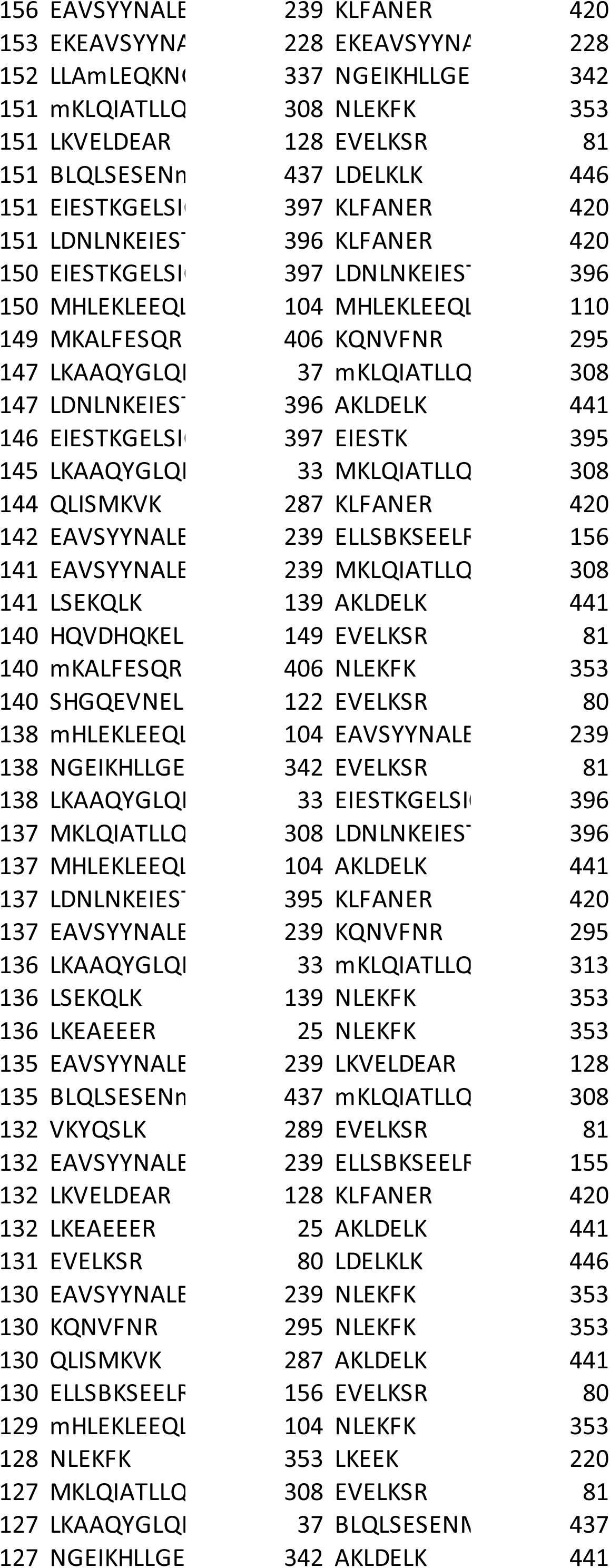
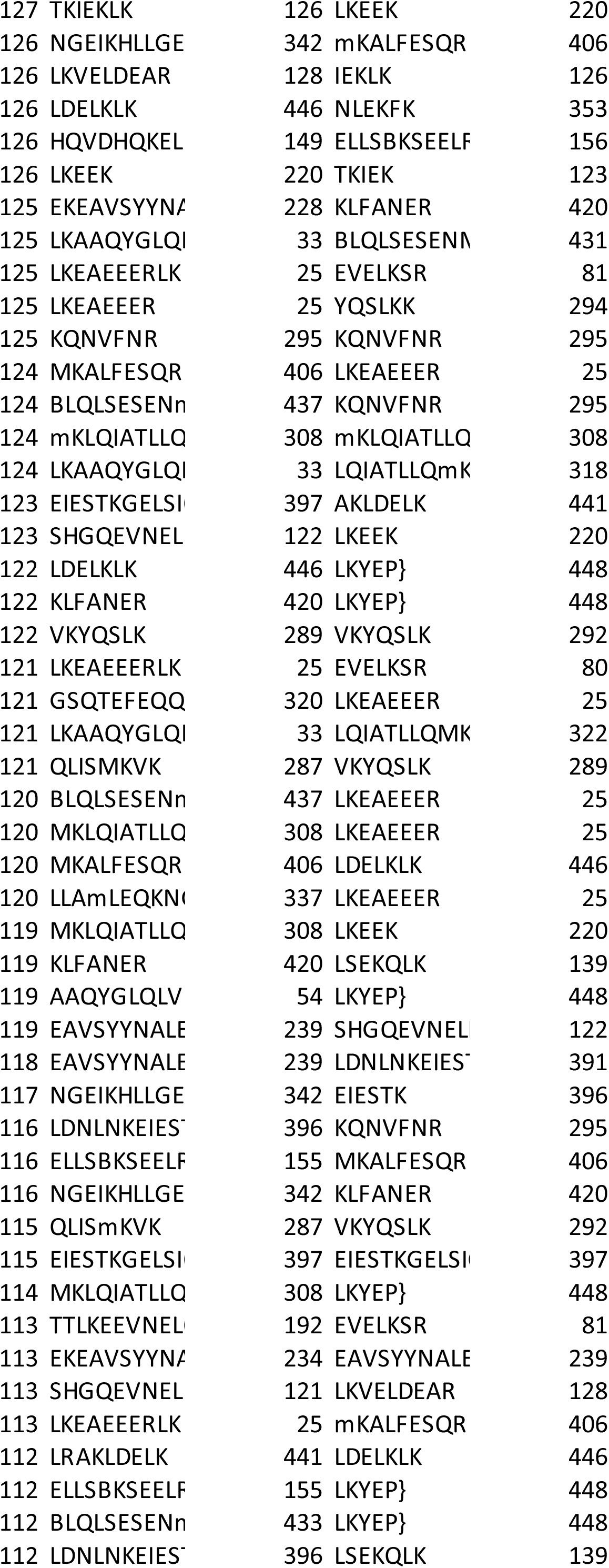
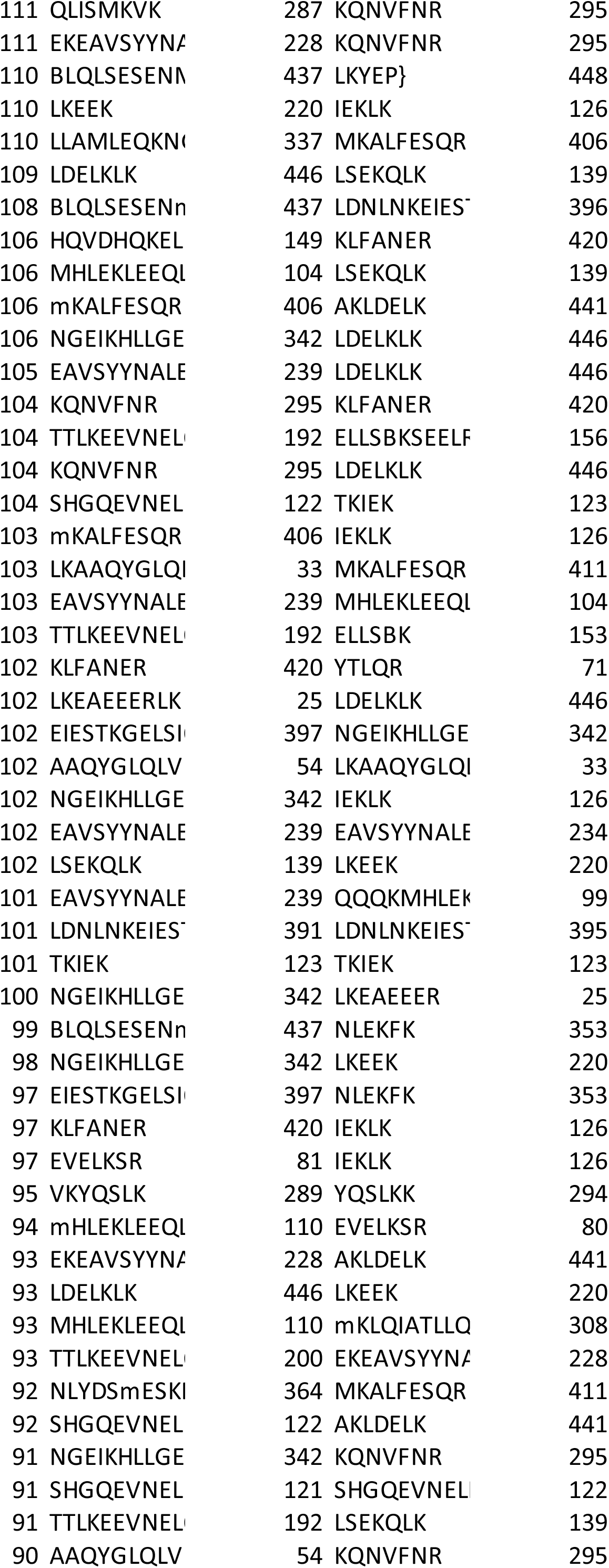
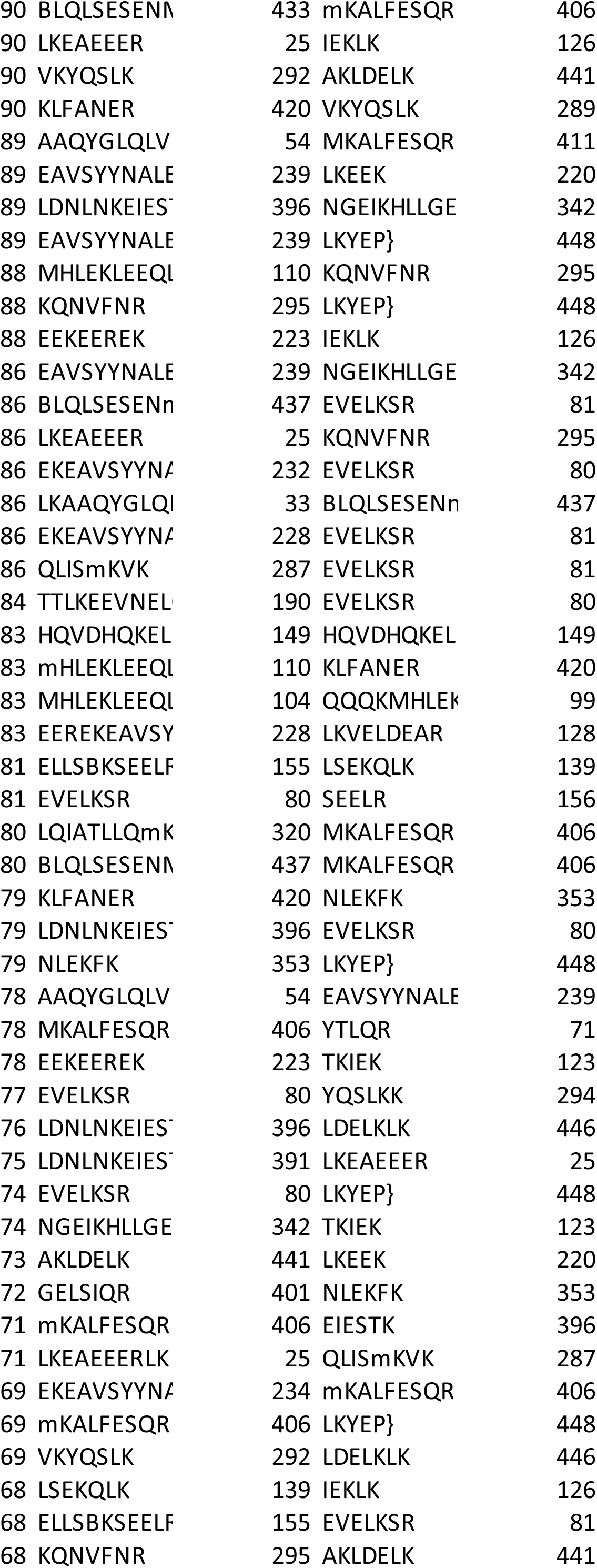
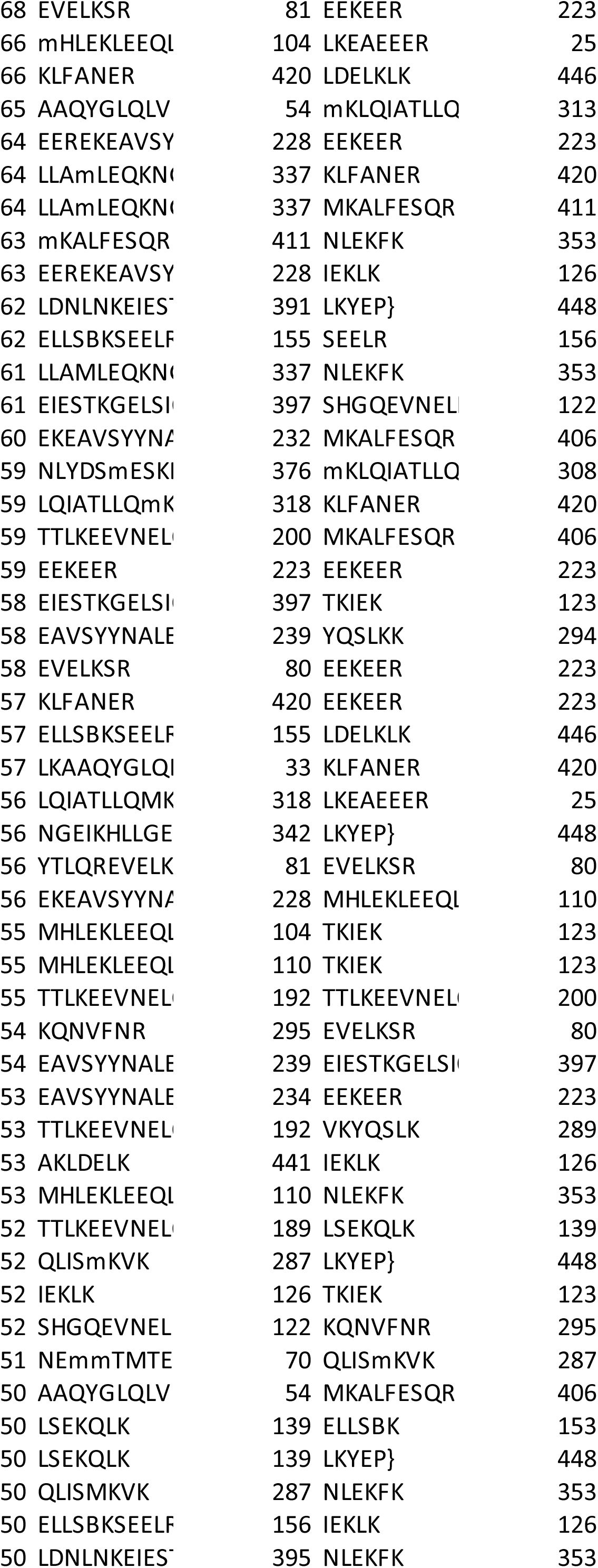
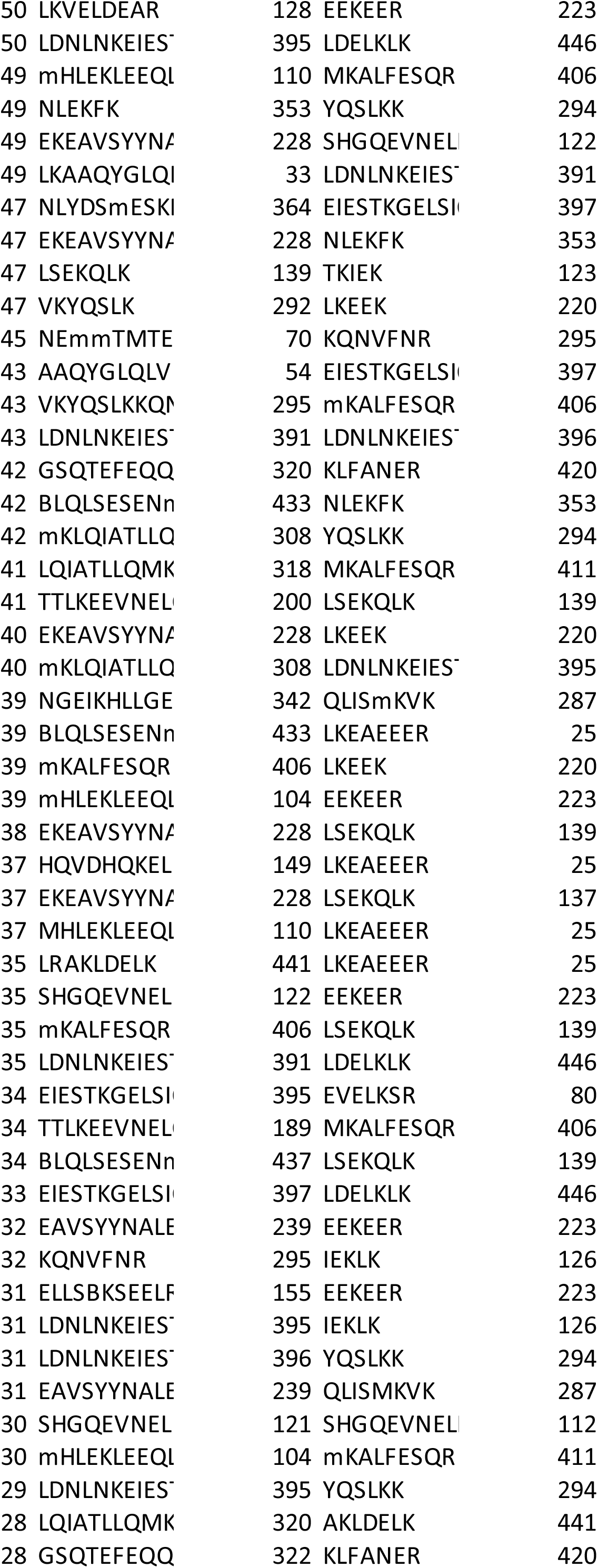
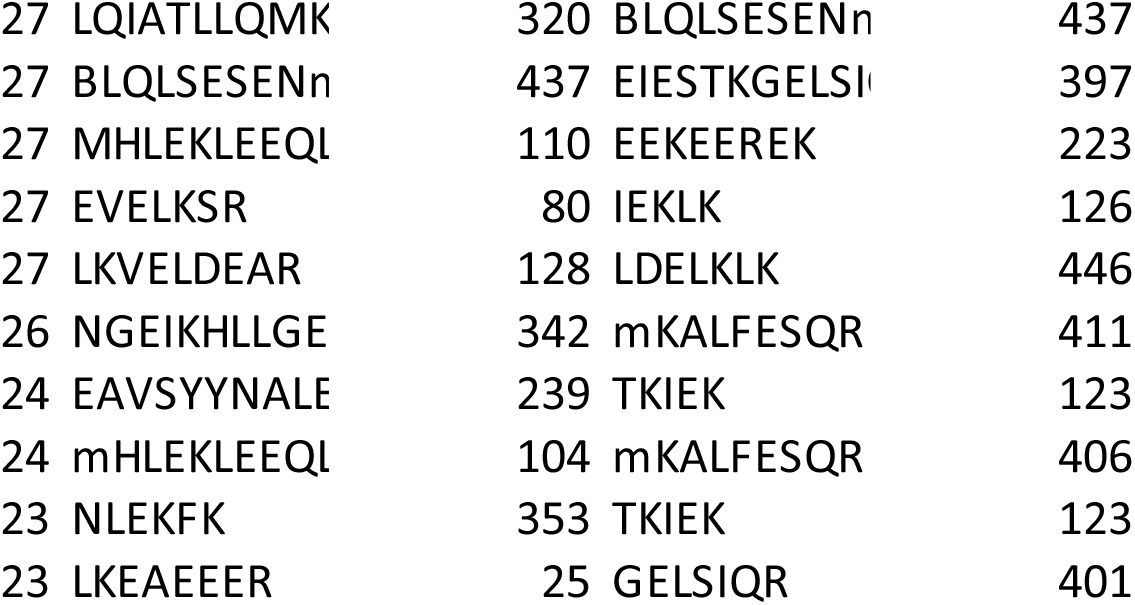

## Notes

### Competing Interest Statement

The authors have declared no competing interest.

## References

Ai, H.W., W. Shen, A. Sagi, P.R. Chen, and P.G. Schultz. 2011. Probing protein-protein interactions with a genetically encoded photo-crosslinking amino acid. Chembiochem. 12:1854–1857.

Akhmanova, A., and C.C. Hoogenraad. 2015. Microtubule minus-end-targeting proteins. Curr Biol. 25:R162–171.

Alex, A., V. Piano, S. Polley, M. Stuiver, S. Voss, G. Ciossani, K. Overlack, B. Voss, S. Wohlgemuth, A. Petrovic, Y. Wu, P. Selenko, A. Musacchio, and S. Maffini. 2019. Electroporated recombinant proteins as tools for in vivo functional complementation, imaging and chemical biology. Elife. 8.

Amos, L.A. 1989. Brain dynein crossbridges microtubules into bundles. J Cell Sci. 93 ( Pt 1):19–28.

Auckland, P., E. Roscioli, H.L.E. Coker, and A.D. McAinsh. 2020. CENP-F stabilizes kinetochore-microtubule attachments and limits dynein stripping of corona cargoes. J Cell Biol. 219.

Barbosa, J., T. Martins, T. Bange, L. Tao, C. Conde, and C. Sunkel. 2020. Polo regulates Spindly to prevent premature stabilization of kinetochore-microtubule attachments. EMBO J. 39:e100789.

Barisic, M., B. Sohm, P. Mikolcevic, C. Wandke, V. Rauch, T. Ringer, M. Hess, G. Bonn, and S. Geley. 2010. Spindly/CCDC99 is required for efficient chromosome congression and mitotic checkpoint regulation. Mol Biol Cell. 21:1968–1981.

Basto, R., F. Scaerou, S. Mische, E. Wojcik, C. Lefebvre, R. Gomes, T. Hays, and R. Karess. 2004. In vivo dynamics of the rough deal checkpoint protein during Drosophila mitosis. Curr Biol. 14:56–61.

Bowler, M.W., D. Nurizzo, R. Barrett, A. Beteva, M. Bodin, H. Caserotto, S. Delageniere, F. Dobias, D. Flot, T. Giraud, N. Guichard, M. Guijarro, M. Lentini, G.A. Leonard, S. McSweeney, M. Oskarsson, W. Schmidt, A. Snigirev, D. von Stetten, J. Surr, O. Svensson, P. Theveneau, and C. Mueller-Dieckmann. 2015. MASSIF-1: a beamline dedicated to the fully automatic characterization and data collection from crystals of biological macromolecules. J Synchrotron Radiat. 22:1540–1547.

Brown, P.H., and P. Schuck. 2006. Macromolecular size-and-shape distributions by sedimentation velocity analytical ultracentrifugation. Biophys J. 90:4651–4661.

Canty, J.T., R. Tan, E. Kusakci, J. Fernandes, and A. Yildiz. 2021. Structure and Mechanics of Dynein Motors. Annu Rev Biophys. 50:549–574.

Carter, A.P., A.G. Diamant, and L. Urnavicius. 2016. How dynein and dynactin transport cargos: a structural perspective. Curr Opin Struct Biol. 37:62–70.

Celestino, R., M.A. Henen, J.B. Gama, C. Carvalho, M. McCabe, D.J. Barbosa, A. Born, P.J. Nichols, A.X. Carvalho, R. Gassmann, and B. Vogeli. 2019. A transient helix in the disordered region of dynein light intermediate chain links the motor to structurally diverse adaptors for cargo transport. PLoS Biol. 17:e3000100.

Chan, Y.W., L.L. Fava, A. Uldschmid, M.H. Schmitz, D.W. Gerlich, E.A. Nigg, and A. Santamaria. 2009. Mitotic control of kinetochore-associated dynein and spindle orientation by human Spindly. J Cell Biol. 185:859–874.

Cheerambathur, D.K., R. Gassmann, B. Cook, K. Oegema, and A. Desai. 2013. Crosstalk between microtubule attachment complexes ensures accurate chromosome segregation. Science. 342:1239–1242.

Chowdhury, S., S.A. Ketcham, T.A. Schroer, and G.C. Lander. 2015. Structural organization of the dynein-dynactin complex bound to microtubules. Nat Struct Mol Biol. 22:345–347.

Civril, F., A. Wehenkel, F.M. Giorgi, S. Santaguida, A. Di Fonzo, G. Grigorean, F.D. Ciccarelli, and A. Musacchio. 2010. Structural analysis of the RZZ complex reveals common ancestry with multisubunit vesicle tethering machinery. Structure. 18:616–626.

Davis, L., and J.W. Chin. 2012. Designer proteins: applications of genetic code expansion in cell biology. Nat Rev Mol Cell Biol. 13:168–182.

Del Castillo, U., H.J. Muller, and V.I. Gelfand. 2020. Kinetochore protein Spindly controls microtubule polarity in Drosophila axons. Proc Natl Acad Sci U S A. 117:12155–12163.

Duvaud, S., C. Gabella, F. Lisacek, H. Stockinger, V. Ioannidis, and C. Durinx. 2021. Expasy, the Swiss Bioinformatics Resource Portal, as designed by its users. Nucleic Acids Res. 49:W216–W227.

Emsley, P., B. Lohkamp, W.G. Scott, and K. Cowtan. 2010. Features and development of Coot. Acta Crystallogr D Biol Crystallogr. 66:486–501.

Evans, R., M. O’Neill, A. Pritzel, N. Antropova, A. Senior, T. Green, A. Žídek, R. Bates, S. Blackwell, J. Yim, O. Ronneberger, S. Bodenstein, M. Zielinski, A. Bridgland, A. Potapenko, A. Cowie, K. Tunyasuvunakool, R. Jain, E. Clancy, P. Kohli, J. Jumper, and H. D. 2021. Protein complex prediction with AlphaFold-Multimer. BioRXiv.

Gama, J.B., C. Pereira, P.A. Simoes, R. Celestino, R.M. Reis, D.J. Barbosa, H.R. Pires, C. Carvalho, J. Amorim, A.X. Carvalho, D.K. Cheerambathur, and R. Gassmann. 2017. Molecular mechanism of dynein recruitment to kinetochores by the Rod-Zw10-Zwilch complex and Spindly. J Cell Biol. 216:943–960.

Gassmann, R., A. Essex, J.S. Hu, P.S. Maddox, F. Motegi, A. Sugimoto, S.M. O’Rourke, B. Bowerman, I. McLeod, J.R. Yates, 3rd, K. Oegema, I.M. Cheeseman, and A. Desai. 2008. A new mechanism controlling kinetochore-microtubule interactions revealed by comparison of two dynein-targeting components: SPDL-1 and the Rod/Zwilch/Zw10 complex. Genes Dev. 22:2385-2399.

Gassmann, R., A.J. Holland, D. Varma, X. Wan, F. Civril, D.W. Cleveland, K. Oegema, E.D. Salmon, and A. Desai. 2010. Removal of Spindly from microtubule-attached kinetochores controls spindle checkpoint silencing in human cells. Genes Dev. 24:957–971.

Gibson, D.G., L. Young, R.Y. Chuang, J.C. Venter, C.A. Hutchison, 3rd, and H.O. Smith. 2009. Enzymatic assembly of DNA molecules up to several hundred kilobases. Nat Methods. 6:343–345.

Graham, M., C. Combe, L. Kobowski, and J. Rappsilber. 2019. xiView: a common platform for the downstream analysis of crosslinking mass spectrometry data. BioRxiv.

Griffis, E.R., N. Stuurman, and R.D. Vale. 2007. Spindly, a novel protein essential for silencing the spindle assembly checkpoint, recruits dynein to the kinetochore. J Cell Biol. 177:1005–1015.

Grimm, M., T. Zimniak, A. Kahraman, and F. Herzog. 2015. xVis: a web server for the schematic visualization and interpretation of crosslink-derived spatial restraints. Nucleic Acids Res. 43:W362–369.

Grotjahn, D.A., S. Chowdhury, Y. Xu, R.J. McKenney, T.A. Schroer, and G.C. Lander. 2018. Cryo-electron tomography reveals that dynactin recruits a team of dyneins for processive motility. Nat Struct Mol Biol. 25:203–207.

Henen, M.A., W. Myers, L.R. Schmitt, K.J. Wade, A. Born, P.J. Nichols, and B. Vogeli. 2021. The Disordered Spindly C-terminus Interacts with RZZ Subunits ROD-1 and ZWL-1 in the Kinetochore through the Same Sites in C. Elegans. J Mol Biol. 433:166812.

Holland, A.J., R.M. Reis, S. Niessen, C. Pereira, D.A. Andres, H.P. Spielmann, D.W. Cleveland, A. Desai, and R. Gassmann. 2015. Preventing farnesylation of the dynein adaptor Spindly contributes to the mitotic defects caused by farnesyltransferase inhibitors. Mol Biol Cell. 26:1845–1856.

Hoogenraad, C.C., and A. Akhmanova. 2016. Bicaudal D Family of Motor Adaptors: Linking Dynein Motility to Cargo Binding. Trends Cell Biol. 26:327–340.

Hoogenraad, C.C., P. Wulf, N. Schiefermeier, T. Stepanova, N. Galjart, J.V. Small, F. Grosveld, C.I. de Zeeuw, and A. Akhmanova. 2003. Bicaudal D induces selective dynein-mediated microtubule minus end-directed transport. EMBO J. 22:6004–6015.

Howell, B.J., B.F. McEwen, J.C. Canman, D.B. Hoffman, E.M. Farrar, C.L. Rieder, and E.D. Salmon. 2001. Cytoplasmic dynein/dynactin drives kinetochore protein transport to the spindle poles and has a role in mitotic spindle checkpoint inactivation. J Cell Biol. 155:1159–1172.

Hueschen, C.L., S.J. Kenny, K. Xu, and S. Dumont. 2017. NuMA recruits dynein activity to microtubule minus-ends at mitosis. Elife. 6.

Huis In ’t Veld, P.J., S. Wohlgemuth, C. Koerner, F. Muller, P. Janning, and A. Musacchio. 2021. Reconstitution and use of highly active human CDK1:Cyclin-B:CKS1 complexes. Protein Sci.

Isabet, T., G. Montagnac, K. Regazzoni, B. Raynal, F. El Khadali, P. England, M. Franco, P. Chavrier, A. Houdusse, and J. Menetrey. 2009. The structural basis of Arf effector specificity: the crystal structure of ARF6 in a complex with JIP4. EMBO J. 28:2835–2845.

Joosten, R.P., F. Long, G.N. Murshudov, and A. Perrakis. 2014. The PDB_REDO server for macromolecular structure model optimization. IUCrJ. 1:213–220.

Jumper, J., R. Evans, A. Pritzel, T. Green, M. Figurnov, O. Ronneberger, K. Tunyasuvunakool, R. Bates, A. Zidek, A. Potapenko, A. Bridgland, C. Meyer, S.A.A. Kohl, A.J. Ballard, A. Cowie, B. Romera-Paredes, S. Nikolov, R. Jain, J. Adler, T. Back, S. Petersen, D. Reiman, E. Clancy, M. Zielinski, M. Steinegger, M. Pacholska, T. Berghammer, S. Bodenstein, D. Silver, O. Vinyals, A.W. Senior, K. Kavukcuoglu, P. Kohli, and D. Hassabis. 2021. Highly accurate protein structure prediction with AlphaFold. Nature. 596:583–589.

Kabsch, W. 2010. Xds. Acta Crystallogr D Biol Crystallogr. 66:125–132.

Kelkar, N., S. Gupta, M. Dickens, and R.J. Davis. 2000. Interaction of a mitogen-activated protein kinase signaling module with the neuronal protein JIP3. Mol Cell Biol. 20:1030–1043.

Klinman, E., and E.L.F. Holzbaur. 2018. Walking Forward with Kinesin. Trends Neurosci. 41:555–556.

Kops, G., and R. Gassmann. 2020. Crowning the Kinetochore: The Fibrous Corona in Chromosome Segregation. Trends Cell Biol. 30:653–667.

Lau, C.K., F.J. O’Reilly, B. Santhanam, S.E. Lacey, J. Rappsilber, and A.P. Carter. 2021. Cryo-EM reveals the complex architecture of dynactin’s shoulder region and pointed end. EMBO J. 40:e106164.

Lee, I.G., S.E. Cason, S.S. Alqassim, E.L.F. Holzbaur, and R. Dominguez. 2020. A tunable LIC1-adaptor interaction modulates dynein activity in a cargo-specific manner. Nat Commun. 11:5695.

Lee, I.G., M.A. Olenick, M. Boczkowska, C. Franzini-Armstrong, E.L.F. Holzbaur, and R. Dominguez. 2018. A conserved interaction of the dynein light intermediate chain with dynein-dynactin effectors necessary for processivity. Nat Commun. 9:986.

Liu, Y., H.K. Salter, A.N. Holding, C.M. Johnson, E. Stephens, P.J. Lukavsky, J. Walshaw, and S.L. Bullock. 2013. Bicaudal-D uses a parallel, homodimeric coiled coil with heterotypic registry to coordinate recruitment of cargos to dynein. Genes Dev. 27:1233–1246.

Luna-Vargas, M.P., E. Christodoulou, A. Alfieri, W.J. van Dijk, M. Stadnik, R.G. Hibbert, D.D. Sahtoe, M. Clerici, V.D. Marco, D. Littler, P.H. Celie, T.K. Sixma, and A. Perrakis. 2011. Enabling high-throughput ligation-independent cloning and protein expression for the family of ubiquitin specific proteases. J Struct Biol. 175:113–119.

Magidson, V., R. Paul, N. Yang, J.G. Ault, C.B. O’Connell, I. Tikhonenko, B.F. McEwen, A. Mogilner, and A. Khodjakov. 2015. Adaptive changes in the kinetochore architecture facilitate proper spindle assembly. Nat Cell Biol.

McKenney, R.J., W. Huynh, M.E. Tanenbaum, G. Bhabha, and R.D. Vale. 2014. Activation of cytoplasmic dynein motility by dynactin-cargo adapter complexes. Science. 345:337–341.

Mirdita, M., K. Schütze, L. Moriqwaki, S. Ovchinnikov, and M. Steinegger. 2021. ColabFold -Making protein folding accessible to all. BioRXiv.

Mische, S., Y. He, L. Ma, M. Li, M. Serr, and T.S. Hays. 2008. Dynein light intermediate chain: an essential subunit that contributes to spindle checkpoint inactivation. Mol Biol Cell. 19:4918–4929.

Mosalaganti, S., J. Keller, A. Altenfeld, M. Winzker, P. Rombaut, M. Saur, A. Petrovic, A. Wehenkel, S. Wohlgemuth, F. Muller, S. Maffini, T. Bange, F. Herzog, H. Waldmann, S. Raunser, and A. Musacchio. 2017. Structure of the RZZ complex and molecular basis of its interaction with Spindly. J Cell Biol. 216:961–981.

Moudgil, D.K., N. Westcott, J.K. Famulski, K. Patel, D. Macdonald, H. Hang, and G.K. Chan. 2015. A novel role of farnesylation in targeting a mitotic checkpoint protein, human Spindly, to kinetochores. J Cell Biol. 208:881–896.

Murshudov, G.N., P. Skubak, A.A. Lebedev, N.S. Pannu, R.A. Steiner, R.A. Nicholls, M.D. Winn, F. Long, and A.A. Vagin. 2011. REFMAC5 for the refinement of macromolecular crystal structures. Acta Crystallogr D Biol Crystallogr. 67:355–367.

Musacchio, A. 2015. The Molecular Biology of Spindle Assembly Checkpoint Signaling Dynamics. Curr Biol. 25:R1002–1018.

Musacchio, A., and A. Desai. 2017. A Molecular View of Kinetochore Assembly and Function. Biology (Basel). 6.

Olenick, M.A., and E.L.F. Holzbaur. 2019. Dynein activators and adaptors at a glance. J Cell Sci. 132.

Olenick, M.A., M. Tokito, M. Boczkowska, R. Dominguez, and E.L. Holzbaur. 2016. Hook Adaptors Induce Unidirectional Processive Motility by Enhancing the Dynein-Dynactin Interaction. J Biol Chem. 291:18239–18251.

Pan, D., A. Brockmeyer, F. Mueller, A. Musacchio, and T. Bange. 2018. Simplified Protocol for Cross-linking Mass Spectrometry Using the MS-Cleavable Cross-linker DSBU with Efficient Cross-link Identification. Anal Chem. 90:10990–10999.

Pannu, N.S., W.J. Waterreus, P. Skubak, I. Sikharulidze, J.P. Abrahams, and R.A. de Graaff. 2011. Recent advances in the CRANK software suite for experimental phasing. Acta Crystallogr D Biol Crystallogr. 67:331–337.

Pavin, N., and I.M. Tolic. 2021. Mechanobiology of the Mitotic Spindle. Dev Cell. 56:192–201.

Pereira, C., R.M. Reis, J.B. Gama, R. Celestino, D.K. Cheerambathur, A.X. Carvalho, and R. Gassmann. 2018. Self-Assembly of the RZZ Complex into Filaments Drives Kinetochore Expansion in the Absence of Microtubule Attachment. Curr Biol. 28:3408–3421 e3408.

Raaijmakers, J.A., M.E. Tanenbaum, and R.H. Medema. 2013. Systematic dissection of dynein regulators in mitosis. J Cell Biol. 201:201–215.

Raisch, T., G. Ciossani, E. D’Amico, V. Cmentowski, S. Carmignani, S. Maffini, F. Merino, S. Wohlgemuth, I.R. Vetter, S. Raunser, and A. Musacchio. 2021. Structure of the RZZ complex and molecular basis of Spindly-driven corona assembly at human kinetochores. BioRVix.

Reck-Peterson, S.L., W.B. Redwine, R.D. Vale, and A.P. Carter. 2018. The cytoplasmic dynein transport machinery and its many cargoes. Nat Rev Mol Cell Biol. 19:382–398.

Redwine, W.B., M.E. DeSantis, I. Hollyer, Z.M. Htet, P.T. Tran, S.K. Swanson, L. Florens, M.P. Washburn, and S.L. Reck-Peterson. 2017. The human cytoplasmic dynein interactome reveals novel activators of motility. Elife. 6.

Renna, C., F. Rizzelli, M. Carminati, C. Gaddoni, L. Pirovano, V. Cecatiello, S. Pasqualato, and M. Mapelli. 2020. Organizational Principles of the NuMA-Dynein Interaction Interface and Implications for Mitotic Spindle Functions. Structure. 28:820–829 e826.

Roberts, A.J., T. Kon, P.J. Knight, K. Sutoh, and S.A. Burgess. 2013. Functions and mechanics of dynein motor proteins. Nat Rev Mol Cell Biol. 14:713–726.

Rodriguez-Rodriguez, J.A., C. Lewis, K.L. McKinley, V. Sikirzhytski, J. Corona, J. Maciejowski, A. Khodjakov, I.M. Cheeseman, and P.V. Jallepalli. 2018. Distinct Roles of RZZ and Bub1-KNL1 in Mitotic Checkpoint Signaling and Kinetochore Expansion. Curr Biol. 28:3422–3429 e3425.

Sacristan, C., M.U.D. Ahmad, J. Keller, J. Fermie, V. Groenewold, E. Tromer, A. Fish, R. Melero, J.M. Carazo, J. Klumperman, A. Musacchio, A. Perrakis, and G.J. Kops. 2018. Dynamic kinetochore size regulation promotes microtubule capture and chromosome biorientation in mitosis. Nat Cell Biol. 20:800–810.

Scheres, S.H. 2012. RELION: implementation of a Bayesian approach to cryo-EM structure determination. J Struct Biol. 180:519–530.

Schindelin, J., I. Arganda-Carreras, E. Frise, V. Kaynig, M. Longair, T. Pietzsch, S. Preibisch, C. Rueden, S. Saalfeld, B. Schmid, J.Y. Tinevez, D.J. White, V. Hartenstein, K. Eliceiri, P. Tomancak, and A. Cardona. 2012. Fiji: an open-source platform for biological-image analysis. Nat Methods. 9:676–682.

Schlager, M.A., H.T. Hoang, L. Urnavicius, S.L. Bullock, and A.P. Carter. 2014a. In vitro reconstitution of a highly processive recombinant human dynein complex. EMBO J. 33:1855–1868.

Schlager, M.A., A. Serra-Marques, I. Grigoriev, L.F. Gumy, M. Esteves da Silva, P.S. Wulf, A. Akhmanova, and C.C. Hoogenraad. 2014b. Bicaudal d family adaptor proteins control the velocity of Dynein-based movements. Cell Rep. 8:1248–1256.

Schroeder, C.M., J.M. Ostrem, N.T. Hertz, and R.D. Vale. 2014. A Ras-like domain in the light intermediate chain bridges the dynein motor to a cargo-binding region. Elife. 3:e03351.

Schroeder, C.M., and R.D. Vale. 2016. Assembly and activation of dynein-dynactin by the cargo adaptor protein Hook3. J Cell Biol. 214:309–318.

Sessa, F., M. Mapelli, C. Ciferri, C. Tarricone, L.B. Areces, T.R. Schneider, P.T. Stukenberg, and A. Musacchio. 2005. Mechanism of Aurora B activation by INCENP and inhibition by hesperadin. Mol Cell. 18:379–391.

Siegel, L.M., and K.J. Monty. 1966. Determination of molecular weights and frictional ratios of proteins in impure systems by use of gel filtration and density gradient centrifugation. Application to crude preparations of sulfite and hydroxylamine reductases. Biochim Biophys Acta. 112:346–362.

Sivaram, M.V., T.L. Wadzinski, S.D. Redick, T. Manna, and S.J. Doxsey. 2009. Dynein light intermediate chain 1 is required for progress through the spindle assembly checkpoint. EMBO J. 28:902–914.

Splinter, D., D.S. Razafsky, M.A. Schlager, A. Serra-Marques, I. Grigoriev, J. Demmers, N. Keijzer, K. Jiang, I. Poser, A.A. Hyman, C.C. Hoogenraad, S.J. King, and A. Akhmanova. 2012. BICD2, dynactin, and LIS1 cooperate in regulating dynein recruitment to cellular structures. Mol Biol Cell. 23:4226–4241.

Starr, D.A., B.C. Williams, T.S. Hays, and M.L. Goldberg. 1998. ZW10 helps recruit dynactin and dynein to the kinetochore. J Cell Biol. 142:763–774.

Stuurman, N., M. Haner, B. Sasse, W. Hubner, B. Suter, and U. Aebi. 1999. Interactions between coiled-coil proteins: Drosophila lamin Dm0 binds to the bicaudal-D protein. Eur J Cell Biol. 78:278–287.

Tang, G., L. Peng, P.R. Baldwin, D.S. Mann, W. Jiang, I. Rees, and S.J. Ludtke. 2007. EMAN2: an extensible image processing suite for electron microscopy. J Struct Biol. 157:38–46.

Terawaki, S., A. Yoshikane, Y. Higuchi, and K. Wakamatsu. 2015. Structural basis for cargo binding and autoinhibition of Bicaudal-D1 by a parallel coiled-coil with homotypic registry. Biochem Biophys Res Commun. 460:451–456.

Urnavicius, L., C.K. Lau, M.M. Elshenawy, E. Morales-Rios, C. Motz, A. Yildiz, and A.P. Carter. 2018. Cryo-EM shows how dynactin recruits two dyneins for faster movement. Nature. 554:202–206.

Urnavicius, L., K. Zhang, A.G. Diamant, C. Motz, M.A. Schlager, M. Yu, N.A. Patel, C.V. Robinson, and A.P. Carter. 2015. The structure of the dynactin complex and its interaction with dynein. Science. 347:1441–1446.

Varma, D., P. Monzo, S.A. Stehman, and R.B. Vallee. 2008. Direct role of dynein motor in stable kinetochore-microtubule attachment, orientation, and alignment. J Cell Biol. 182:1045–1054.

Weissmann, F., G. Petzold, R. VanderLinden, P.J. Huis In ’t Veld, N.G. Brown, F. Lampert, S. Westermann, H. Stark, B.A. Schulman, and J.M. Peters. 2016. biGBac enables rapid gene assembly for the expression of large multisubunit protein complexes. Proc Natl Acad Sci U S A. 113:E2564–2569.

Williams, B.C., M. Gatti, and M.L. Goldberg. 1996. Bipolar spindle attachments affect redistributions of ZW10, a Drosophila centromere/kinetochore component required for accurate chromosome segregation. J Cell Biol. 134:1127–1140.

Williams, C.J., J.J. Headd, N.W. Moriarty, M.G. Prisant, L.L. Videau, L.N. Deis, V. Verma, D.A. Keedy, B.J. Hintze, V.B. Chen, S. Jain, S.M. Lewis, W.B. Arendall, 3rd, J. Snoeyink, P.D. Adams, S.C. Lovell, J.S. Richardson, and D.C. Richardson. 2018. MolProbity: More and better reference data for improved all-atom structure validation. Protein Sci. 27:293–315.

Winn, M.D., C.C. Ballard, K.D. Cowtan, E.J. Dodson, P. Emsley, P.R. Evans, R.M. Keegan, E.B. Krissinel, A.G. Leslie, A. McCoy, S.J. McNicholas, G.N. Murshudov, N.S. Pannu, E.A. Potterton, H.R. Powell, R.J. Read, A. Vagin, and K.S. Wilson. 2011. Overview of the CCP4 suite and current developments. Acta Crystallogr D Biol Crystallogr. 67:235–242.

Wojcik, E., R. Basto, M. Serr, F. Scaerou, R. Karess, and T. Hays. 2001. Kinetochore dynein: its dynamics and role in the transport of the Rough deal checkpoint protein. Nat Cell Biol. 3:1001–1007.

Wu, M., T. Wang, E. Loh, W. Hong, and H. Song. 2005. Structural basis for recruitment of RILP by small GTPase Rab7. EMBO J. 24:1491–1501.

Yamamoto, T.G., S. Watanabe, A. Essex, and R. Kitagawa. 2008. SPDL-1 functions as a kinetochore receptor for MDF-1 in Caenorhabditis elegans. J Cell Biol. 183:187–194.

Yeh, T.Y., N.J. Quintyne, B.R. Scipioni, D.M. Eckley, and T.A. Schroer. 2012. Dynactin’s pointed-end complex is a cargo-targeting module. Mol Biol Cell. 23:3827–3837.

Zhang, K., H.E. Foster, A. Rondelet, S.E. Lacey, N. Bahi-Buisson, A.W. Bird, and A.P. Carter. 2017. Cryo-EM Reveals How Human Cytoplasmic Dynein Is Auto-inhibited and Activated. Cell. 169:1303–1314 e1318.

